# Stable isotopomers of *myo*-inositol to uncover the complex MINPP1-dependent inositol phosphate network

**DOI:** 10.1101/2022.08.29.505671

**Authors:** Minh Nguyen Trung, Stefanie Kieninger, Zeinab Fandi, Danye Qiu, Guizhen Liu, Adolfo Saiardi, Henning Jessen, Bettina Keller, Dorothea Fiedler

## Abstract

The water-soluble inositol phosphates (InsPs) represent a functionally diverse group of small-molecule messengers central to a myriad of cellular processes. However, we have an incomplete understanding of InsP metabolism because the available analytical toolset for inositol phosphates is rather limited. Here, we have synthesized and utilized fully and unsymmetrically ^13^C-labeled *myo*-inositol and inositol phosphates. These probes were applied in combination with nuclear magnetic resonance spectroscopy (NMR) and capillary electrophoresis mass spectrometry (CE-MS) to further annotate central aspects of InsP metabolism in human cells. The labeling strategy provided detailed structural information *via* NMR – down to individual enantiomers – which overcomes a crucial blind spot in the analysis of InsPs. We uncovered a novel branch of InsP dephosphorylation in human cells which is dependent on MINPP1, a phytase-like enzyme, that contributes to cellular homeostasis. Full characterization of MINPP1 activity *in vitro* and in cells, provided a clear picture of this multifunctional phosphatase. Metabolic labeling with stable isotopomers thus constitutes a powerful tool for investigating InsP networks in a variety of different biological contexts.

## Introduction

Inositol polyphosphates (InsPs) are ubiquitous, water-soluble small molecules, found in all eukaryotes. InsPs are involved in a wide spectrum of biological functions as they are key to fundamental physiological processes. A well-characterized example is inositol-1,4,5-trisphosphate (Ins(1,4,5)P_3_) as a Ca^2+^ release factor. More recently, InsPs were shown to regulate the activity of class I histone deacetylases, as well as Bruton’s tyrosine kinase (Btk), which implies a wider role for InsPs in transcriptional regulation and in governing intracellular signal transduction. [1–3]

The InsPs vary greatly with respect to their phosphorylation patterns and over 20 different InsPs are currently thought to be part of mammalian InsP metabolism.[4–7] The most abundant InsPs in mammalian cells are inositol-1,3,4,5,6-pentakisphosphate (InsP_5_[2OH]) and inositol hexakisphosphate (also called phytic acid, InsP_6_), with cellular concentrations ranging from the lower µM range to >100 µM in human cells and even in the sub-mM range in slime molds.[8,9] InsP_5_[2OH] and InsP_6_ are precursors for the biosynthesis of inositol pyrophosphates (PP-InsPs) which have recently drawn increasing attention due to their dense phosphorylation patterns and their involvement in central signaling processes.[10] InsP_6_ is also found in a growing number of proteins and protein complexes as a structural cofactor or as a “molecular glue” for protein-protein interactions.[11–14]

While the kinase-mediated biosynthetic pathways of InsP metabolism are fairly well studied, there is limited information on dephosphorylation of InsPs in mammalian cells,[15] especially with respect to the higher phosphorylated members. To date, MINPP1 (Multiple Inositol Polyphosphate Phosphatase 1) is the only recognized enzyme in the human genome capable of dephosphorylating InsP_6_. [16,17] MINPP1 is related to phytases, a highly conserved group of enzymes in many other organisms that can dephosphorylate various InsPs.[18] MINPP1 has been shown to play a role in apoptosis, ER-related stress, and bone and cartilage tissue formation [17,19]. Recently, MINPP1 was connected to a genetic disorder: patients with loss-of-function mutations in MINPP1 exhibit pontocerebellar hypoplasia, a neurodegenerative disease severely impacting cognitive functions and life expectancy.[20,21] Therefore, it is important to understand the molecular mechanisms of MINPP1-governed functions in health and disease.

Although MINPP1 is annotated as a 3-phosphatase, i.e. it predominantly removes the phosphoryl group at the 3-position on InsP_6_,[22] MINPP1 is known to be able to dephosphorylate several InsPs at different positions with varying affinities and kinetics [23,24]. The current assumption is that MINPP1 can dephosphorylate InsP_6_ to hitherto only sparsely annotated InsP_4/3_ species (Figure 1a).[6,7,25–27] However, this activity has only been demonstrated *in vitro* thus far, and whether this is relevant *in vivo*, and which InsP intermediates are involved, is still not clear.

**Figure 1:**
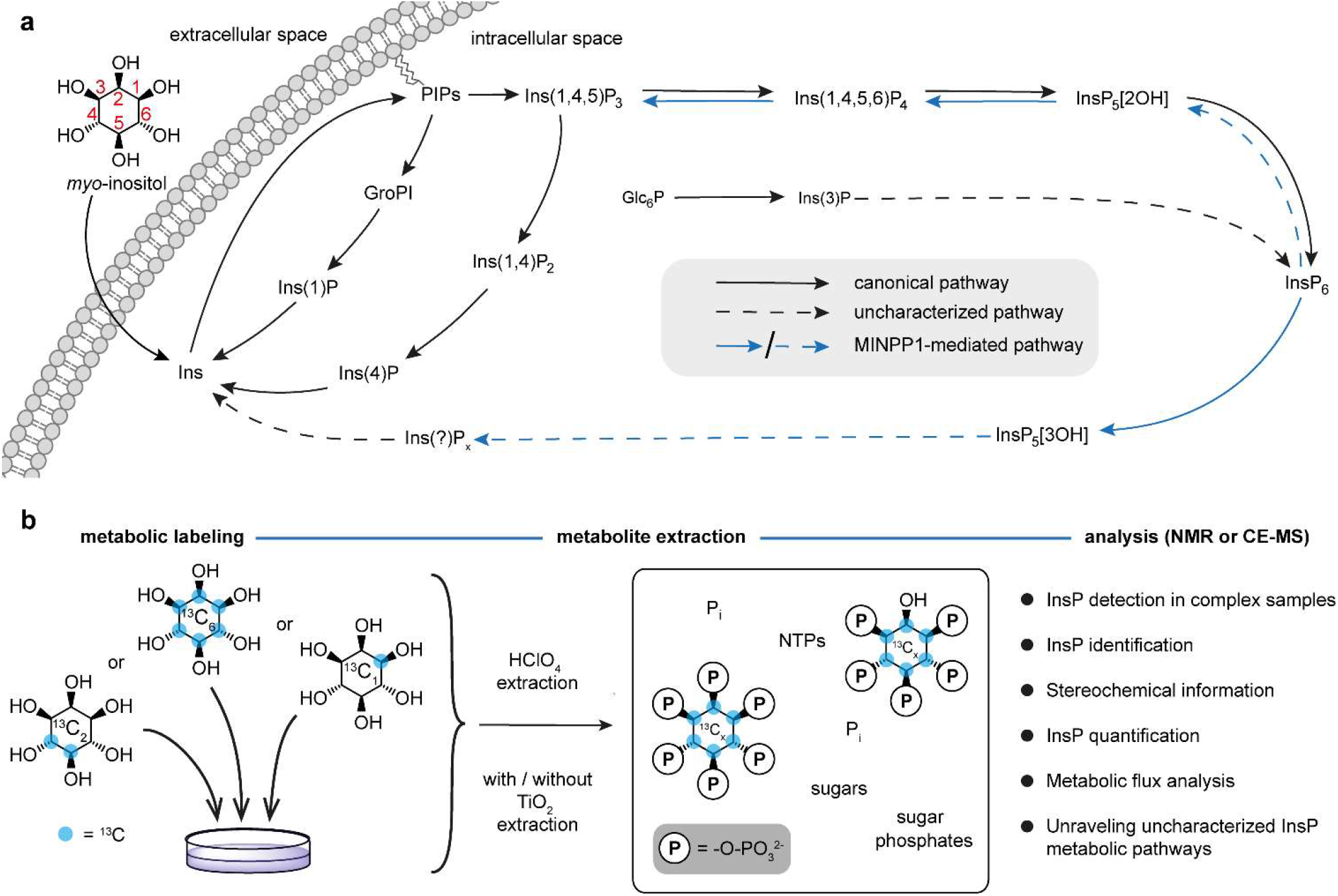
Probing InsP metabolism with *myo*-inositol isotopomers. (**a**) Simplified overview of InsP metabolism with MINPP1-mediated processes highlighted. It is assumed that MINPP1 dephosphorylates InsP_6_ and various other InsPs down to sparsely annotated InsP_3_ isomers. PIPs: phosphatidylinositol phosphates, GroPI: glycerophosphoinositol, Glc6P: glucose-6-phosphate, MINPP1: multiple inositol polyphosphate phosphatase1, Ins(X,Y)P_z_: *myo*-inositol with z phosphoryl groups at positions X,Y; InsP_5_[XOH]: inositol pentakisphosphate with a hydroxyl group at position X. The IUPAC numbering convention of the positions on the inositol scaffold is shown in red. (**b**) Workflow for the analysis of cellular InsP pools through metabolic labeling: human cells are grown in medium devoid of non-labeled *myo-*inositol but supplemented with an isotopomer of *myo-*inositol ([^13^C_6_]Ins, 4,5[^13^C_2_]*myo-*inositol or 1[^13^C_2_]*myo-*inositol) which are incorporated into the cellular InsP pool. The metabolites are then extracted resulting in a complex sample containing all water-soluble biomolecules, such as nucleotide triphosphates (NTPs), inorganic phosphate (Pi) and the labeled InsPs. This mixture can be analyzed *via* NMR or CE-MS exploiting NMR activity and mass difference of the ^13^C label.

Probing and quantifying InsP metabolites and their interconversion is still a challenging task due to the limitations of current analytical tools for InsPs. Many established methods for the detection and analysis of InsPs rely on some form of physico-chemical separation of different InsPs from a complex mixture. The most common methods are strong-anion exchange chromatography (SAX-HPLC)-based fractionation in combination with radiolabeling and scintillation counting, high-density polyacrylamide electrophoresis with cationic staining, or more recently, capillary electrophoresis coupled to mass spectrometry (CE-MS).[26,28–33] Most of these methods are sensitive and powerful for the analysis of highly phosphorylated InsPs, but the separation and detection of lower InsPs (i.e. InsP_1_, InsP_2_, and InsP_3_ species) in a mixture with isobaric sugar phosphates remain difficult. Our group recently established a metabolic labeling strategy using fully isotopically labeled [^13^C_6_]*myo*-inositol. Analysis of the extracted metabolites by 2D nuclear magnetic resonance (NMR) spectroscopy enabled the quantification of higher phosphorylated InsPs (InsP_6_, InsP_5_[2OH], and PP-InsPs; see Figure 1b) without the need for analytical separation.[34] In addition, 2D-NMR measurements provide important information on the InsP phosphorylation patterns and should be able to detect the whole range of InsP metabolites, including the lower phosphorylated species.

Here, we combined fully ^13^C-labeled and asymmetrically ^13^C-labeled isotopomers of *myo*-inositol and InsPs in both biochemical and cellular metabolic labeling experiments. Making use of their inherent properties (position-specific NMR activity and different molecular masses) we uncovered an uncharacterized branch of human InsP metabolism. Ins(2,3)P_2_ and Ins(2)P were identified as major InsPs species in human cells and their levels are dependent on MINPP1 activity towards InsP_6_ *in vitro* and *in cellula*. Through *in vitro* characterization, computational kinetic modeling and metabolic flux *via* CE-MS analysis, we dissect the complex reactivity of MINPP1. We envision that this combined application of *myo-*inositol isotopomers in NMR and CE-MS experiments will help unravel complex InsP networks in different biological contexts in the future.

## Results

### InsP phosphorylation patterns are well resolved by BIRD-{^1^H-^13^C}HMQC NMR spectra

The analysis of complex mixtures of InsP metabolites still constitutes a significant analytical challenge. To identify inositol-derived signals in biological samples *via* NMR in a methodical way, BIRD-{^1^H-^13^C}HMQC-NMR spectra of 19 different InsPs and PP-InsPs (commercially available or synthesized) were recorded and assigned. The collective data of these spectra illustrate that the NMR signals of InsPs cluster in a systematic manner (Figures 2 and S1). NMR signals corresponding to methine group signals adjacent to a non-phosphorylated hydroxyl substituent (**CH**-OH) are separated from methine groups signals with a phosphate substituent (**CH**-O-PO_3_^2-^), which are collectively shifted downfield in both ^1^H and ^13^C dimensions. Within these two groups, clusters for the different positions on the *myo*-inositol ring are apparent. The 2- and 5-positions form clusters of their own, while positions 1 and 3, as well as positions 4 and 6, are intertwined, due to the symmetry plane of the *myo*-inositol ring. Taken together, these combined spectra illustrate that a complete set of NMR signals of an InsP can be used to determine the phosphorylation pattern, and thus the identity, of a given InsP. In the case of chiral InsPs, their NMR spectra cannot be used for a definitive assignment but can narrow the identity down to a pair of enantiomers. For distinguishing two InsP enantiomers, a desymmetrization strategy has to be employed, such as unsymmetrical isotopic labeling of the *myo-*inositol ring with ^13^C, as will be discussed below.

**Figure 2:**
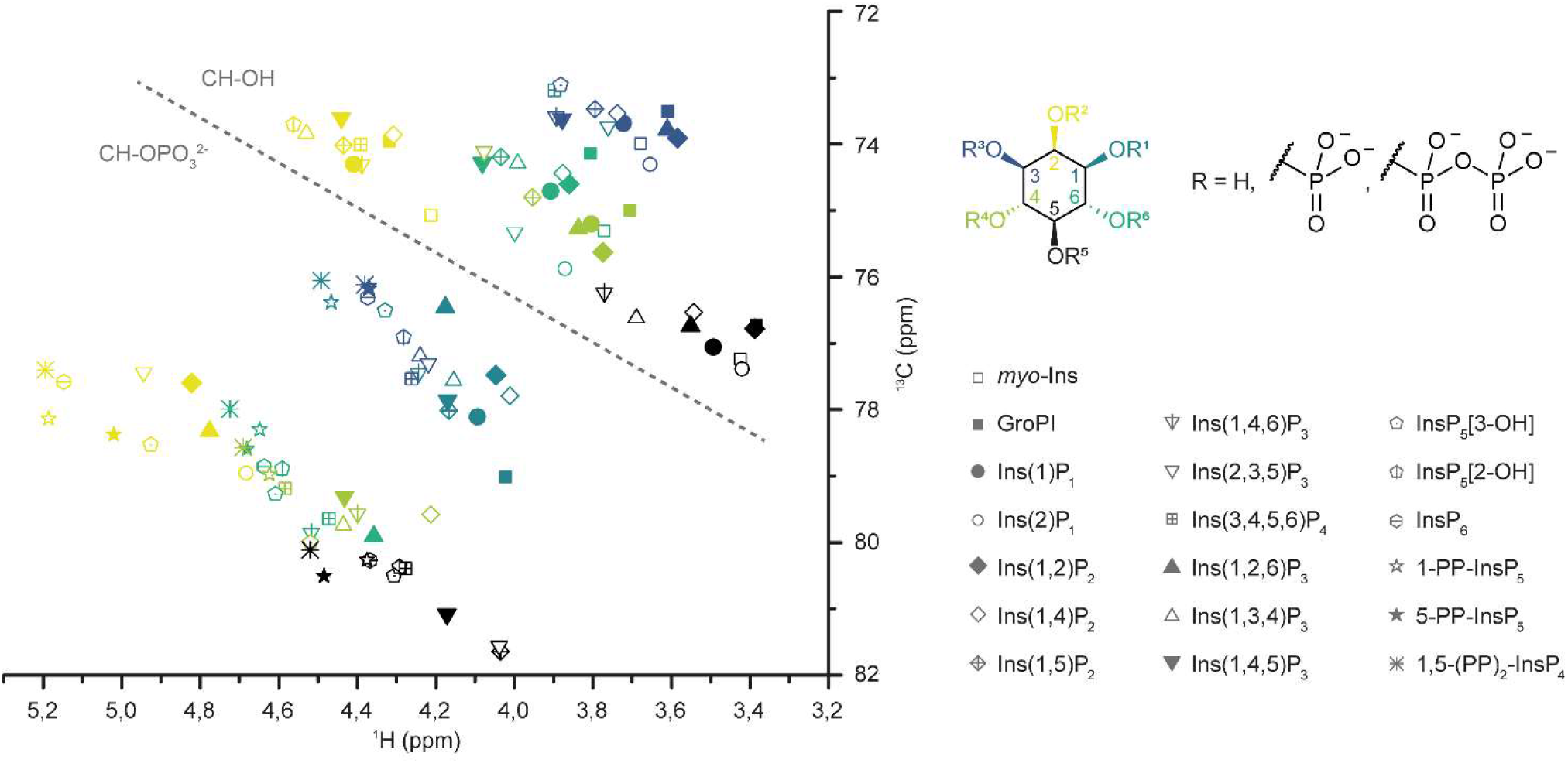
HMQC signals of InsPs with different phosphorylation patterns cluster systematically. Collection of BIRD-{^1^H,^13^C}HMQC NMR data of various InsP standards in metabolic extract buffer conditions (saturated KClO_4_ in D_2_O, pH* 6.0). The HMQC-signals of different InsPs are represented with symbols, while the position on the inositol ring is color-coded. The HMQC-signals cluster together depending on phosphorylation status (dotted line) and position on the inositol ring. The CH groups bearing the pyrophosphate moiety of PP-InsPs or the 1-glycerylphosphate group of GroPI cluster with the phosphorylated CH groups and were treated accordingly for creating bagplots (Figure S1).

### Ins(2,3)P_2_ and Ins(2)P are major mammalian metabolites

We next performed metabolic labeling of human cell lines (HEK293, HCT116, HT29, H1Hela, H1975) with [^13^C_6_]*myo-* inositol (Figure 1b).[34] In brief, cells were grown in a custom medium based on DMEM which contains no natural [^12^C]*myo-*inositol but is instead supplemented with [^13^C_6_]*myo-*inositol or an isotopomer of choice (see below). After the cells incorporated the ^13^C-label into their InsP pool to equilibrium (over 2 passages), cells were harvested and their water-soluble metabolites extracted and analyzed by BIRD-{^1^H-^13^C}HMQC-NMR. This NMR experiment detects ^13^CH groups selectively over non-labeled CH groups, making it particularly suitable for measuring the ^13^C-labeld InsP pool over a complex background. The information from Figure 2 allowed us to annotate all detectable ^13^C-labeled species from such extracts. Quantification was performed through relative integration of the signal corresponding to the 2-position against an internal standard and back-calculated to packed cell volumes. The annotation of the different InsPs in an HCT116 metabolic extract is shown exemplarily in Figure 3a (for full annotation see Figure S2) The same set of InsP species was observed in all other cell lines as well (Figure S3): the major labeled species include InsP_6_, InsP_5_[2OH], 1/3-glycerophospho-*myo*-inositol (1/3-GroPI), inositol 1-or 3-monophosphate (Ins(1/3)P), inositol 1,2-or 2,3-bisphosphate (Ins(1/3,2)P_2_), inositol 2-monophosphate (Ins(2)P) and *myo-*inositol. All of these metabolite assignments were validated through spike-in experiments with commercially available InsP standards into ^13^C-labeled metabolic extracts (Figure S4). In order to differentiate the possible enantiomers in the InsP pool, we synthesized asymmetrically ^13^C-labeled *myo*-inositols following our previously published protocol.[34] Using the singly-labeled isotopomer 1[^13^C_1_]*myo-*inositol and doubly labeled 4,5[^13^C_2_]Ins, respectively, we repeated the metabolic labeling in HEK293 and HCT116 cells. Focusing on the 1[^13^C_1_]*myo-*inositol labeling, the resulting spectra (Figures 3b, S5a) show that the signals that correspond to the phosphorylated 1/3-positions of 1/3-GroPI and Ins(1/3)P are labeled; i.e. the enantiomers present in mammalian cells are 1-GroPI and Ins(1)P. The phosphorylated 1/3-position signal of Ins(1/3,2)P_2_ is not labeled, which identifies Ins(2,3)P_2_ as the prevalent enantiomer, an observation that was reproducible in both cell lines.

**Figure 3:**
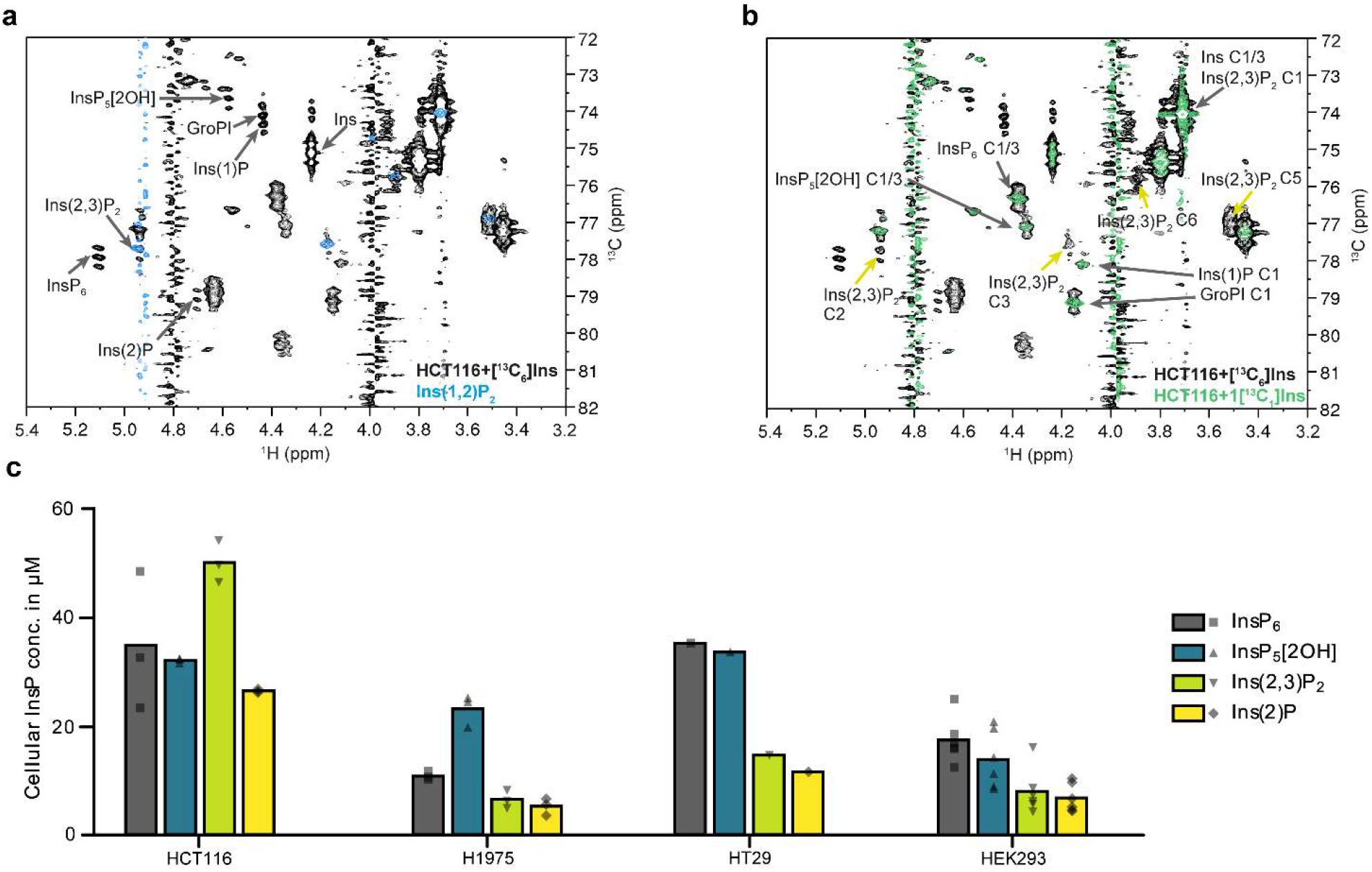
Identification and quantification of major InsPs in human cells. (**a**) Overlay of BIRD-{^1^H,^13^C}HMQC-NMR spectra of metabolic extracts from HCT116 cells which were labeled with [^13^C_6_]*myo-*inositol (black spectrum) and a reference spectrum of Ins(1,2)P_2_ (blue). The annotation of identified InsPs was limited to the NMR signals of the 2-positions for clarity. A complete annotation is provided in Figure S2. (**b**) Overlay of BIRD-{^1^H,^13^C}HMQC-NMR spectra of metabolic extracts from HCT116 cells which were labeled with either [^13^C_6_]*myo*-inositol (black, same data as in (a)) or 1[^13^C_1_]*myo-*inositol (green). Annotation was limited to C1/3 positions (black arrows) and visible signals from Ins(2,3)P_2_ (bright green arrows) for clarity. The labeled 1-positions of GroPI and Ins(1)P confirm their enantiomeric identity. However, the lack of a 1-position label expected for Ins(1,2)P_2_ indicates that the InsP in question is the Ins(2,3)P_2_ enantiomer. (**c**) Scatter dot plot of quantified InsPs from metabolic extracts of various cells (HCT116, n = 3; H1975, n = 3; HT29, n = 1; HEK293, n = 6, biological replicates) with bars representing the means.

GroPI and Ins(1)P are established products of cellular phosphatidylinositide turnover [7,35], their detecton was therefore anticipated. The rather high abundance of Ins(2,3)P_2_ and Ins(2)P in the µM-range (esp. in HCT116 cells, see Figure 3c) was an unexpected observation. Ins(2,3)P_2_ and Ins(2)P have not been associated with any established InsP-related pathway so far. Although Ins(2)P and Ins(1/3,2)P_2_ have been detected in 1995 by Mitchell and colleagues [7,36] these metabolites received little attention and were neglected since then. Overall, the structural information contained in the HMQC-NMR spectra could be used to assign all detectable ^13^C-labeled species in mammalian cells and in combination with the asymmetrical inositol isotopomers, enantiomers could be resolved spectroscopically. This analysis uncovered high amounts of previously poorly characterized lower InsPs, which were not easily accessible with other analytical methods.

### The formation of Ins(2,3)P_2_ and Ins(2)P is dependent on MINPP1

In the biosynthetic pathway towards InsP_6_ there are no InsP intermediates that are phosphorylated at the 2-position. The 2-phosphate group of InsP_6_ is installed only in the last step, in which IPPK (inositol pentakisphosphate 2-kinase) converts InsP_5_[2OH] to InsP_6_. Ins(2,3)P_2_ and Ins(2)P may therefore be generated downstream of InsP_6_. A central InsP phosphatase is the mammalian phytase-like enzyme MINPP1, the only recognized InsP_6_ phosphatase. To investigate possible relationships between Ins(2,3)P_2_, Ins(2)P, and MINPP1, we turned our attention to cells lacking MINPP1. *MINPP1*^*-/-*^ HEK293 cells were labeled with [^13^C_6_]*myo-*inositol and the metabolites were analyzed by NMR (Figure 4). The *MINPP1*^*-/-*^ cells exhibited slightly elevated InsP_6_ levels and accumulated one new InsP species, which was initially assigned as InsP_5_[3OH] or its enantiomer InsP_5_[1OH] (Figure S4e). InsP_5_[1/3OH] was not present in any investigated WT cell line. Labeling of *MINPP1*^*-/-*^ HEK293 cells with the asymmetric isotopomers 1[^13^C_1_]*myo-*inositol and 4,5[^13^C_2_]*myo*-inositol revealed that the 1-position of the new InsP_5_ species is phosphorylated and the shift of the 4-position of the InsP_5_ in question would match InsP_5_[3OH] but not InsP_5_[1OH] (Figure S5).

**Figure 4:**
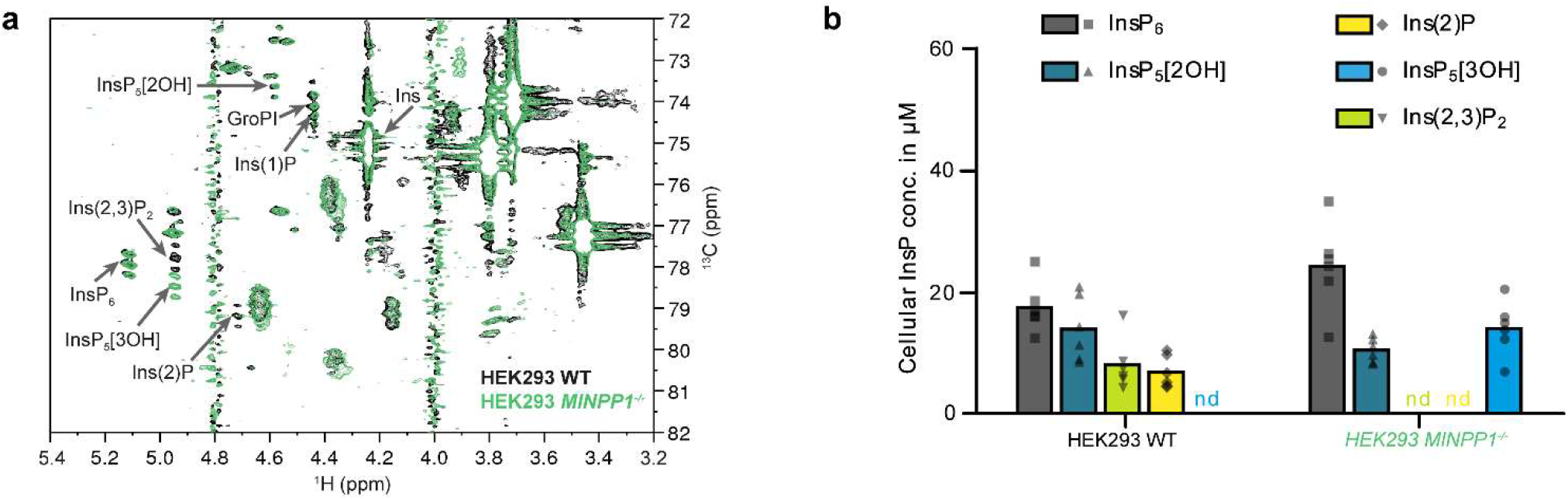
Identification of InsPs in HEK293 and *MINPP1*^*-/-*^ HEK293 cells. (**a**) Overlay of [^13^C_6_]*myo-*inositol-labeled HEK293 (black) and *MINPP1*^*-/-*^ HEK293 cells (green). Ins(2,3)P_2_ and Ins(2)P are not observable in *MINPP1*^*-/-*^ cells, instead InsP_5_[3OH] accumulates. (**b**) Scatter dot plot of quantified InsPs from these cell lines (WT, n = 6 same data as in Figure 3c for illustrative purposes; *MINPP1*^*-/-*^, n = 6, biological replicates). Bars represent the means, nd = not detected. Enantiomer-specific identification of InsP_5_[3OH] is shown in Figure S5.

Strikingly, another change observed in the *MINPP1*^*-/-*^ cell extracps was the complete absence of Ins(2,3)P_2_ and Ins(2)P, establishing a connection between MINPP1 and these lower phosphorylated InsPs. The lack of an undefined InsP_2_ species was also noted in previous analyses of the same cell line, using a radiolabeling approach.[21] Taking into consideration the only sparsely annotated intermediates and products of MINPP1-mediated dephosphorylation of InsP_6_, it seemed possible that MINPP1 could generate Ins(2,3)P_2_ and Ins(2)P directly from InsP_6_.

### MINPP1 dephosphorylates InsP_5_[2OH] and InsP_6_ *via* fully distinct pathways

To validate this hypothesis, we next sought to investigate the *in vitro* activity of MINPP1 against different InsPs. The expression and purification of recombinant MINPP1 in *E. coli* was optimized, to isolate protein yields compatible with biochemical reactions on an NMR scale (Figure S6). Next, MINPP1 was incubated with fully ^13^C_6_-labeled InsP_5_[2OH], and monitored the reaction using 2D NMR measurements. In first experiments we chose a substrate concentration of 50 µM, which is in the middle to upper range of physiological concentrations (Figure S7).[7,30,37] To enable the detection and assignment of all intermediates we subsequently increased the substrate concentration to 175 µM, which did not alter the overall outcome (Figure 5a). The structures of the intermediates were identified using the information from Figure 2 and additional cross-correlation NMR and spike-in experiments where necessary. In agreement with the annotation of MINPP1 as a 3-phosphatase, the first major intermediates for InsP_5_[2OH] dephosphorylation are Ins(1,4,5,6)P_4_ and subsequently Ins(1,4,5)P_3_. MINPP1 thereby directly reverses the phosphorylation reactions mediated by IPMK (inositol phosphate multikinase).[24,38,39] Ins(1,4,5)P_3_ is subsequently converted slowly to a mixture of different InsP_1/2_s (Figure 5b, a full scheme with all minor intermediates is shown in Figure S8).

**Figure 5:**
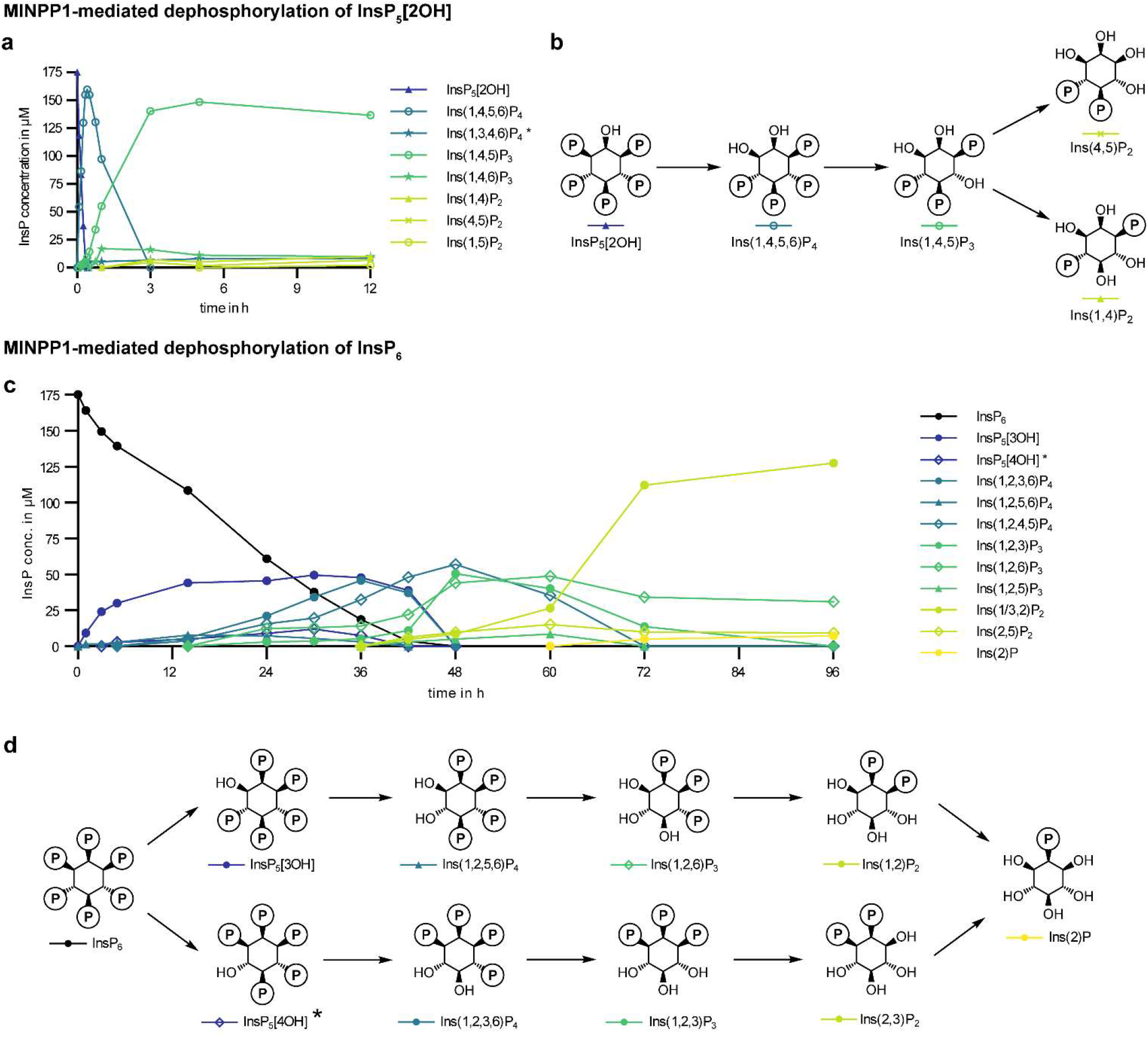
Dephosphorylation of InsP_5_[2OH] and InsP_6_ by MINPP1 *in vitro*. (**a**) Progress curves of MINPP1 reaction with 175 µM [^13^C_6_]InsP_5_[2OH] showing the first 12 h of the reaction (for full scope of progress curves see SI). The progress curves shown here are representative of two replicates. (**b**) Simplified reaction scheme of the MINPP1-mediated dephosphorylation of InsP_5_[2OH]. A complete reaction scheme that includes all minor intermediates is in Figure S8. (**c**) Progress curves of MINPP1 reaction with 175 µM [^13^C_6_]InsP_6_ with simplified reaction scheme depicting the two main reaction paths (**d**). The progress curves shown here are representative of two replicates. A complete reaction scheme that includes all intermediates is in Figure S9. Note that the two enantiomers Ins(1,3)P_2_ and Ins(2,3)P_2_ are quantified together. *: The structure of these InsPs could not be assigned with certainty due to low abundance and interference of more abundant signals.

We then proceeded to probe MINPP1-mediated dephosphorylation of InsP_6_. In contrast to InsP_5_[2OH] as a substrate, we observed a complex mixture of intermediates (Figure 5c). Additionally, the overall conversion of InsP_6_ was visibly slower. The two major reaction paths are depicted in Figure 5d (complete scheme in Figure S9): One dephosphorylation sequence proceeds *via* InsP_5_[3OH] and Ins(1,2,6)P_3_ as intermediates, and a second pathway generates Ins(1,2,3)P_3_ as an intermediate *via* an unidentified (due to low abundance) InsP_5_ isomer. Importantly, Ins(1/3,2)P_2_ and Ins(2)P were observed as the final products of the dephosphorylation of InsP_6_ validating that MINPP1 is responsible for generating these InsPs directly from InsP_6_.

To assess which enantiomers were formed during MINPP1-mediated dephosphorylation of InsP_6_ we synthesized 1[^13^C_1_]InsP_6_.[34] The InsP_5_, InsP_4_ and InsP_3_ intermediates which are produced by MINPP1 from 1[^13^C_1_]InsP_6_ are enantiopure, as no dephosphorylation of the 1-position was observed (detailed explanation in Figure S10a,b). But surprisingly, a mixture of Ins(1,2)P_2_ and Ins(2,3)P_2_ was formed during the later stage of the reaction (Figure S10c,d). The rather high ratio of Ins(2,3)P_2_ to Ins(1,2)P_2_ suggests that Ins(1,2)P_2_ is formed exclusively *via* Ins(1,2,6)P_3_ and Ins(1,2,3)P_3_ is selectively converted to Ins(2,3)P_2_. Both InsP_2_s are, in turn, substrates to form Ins(2)P. Our *in vitro* assessment of MINPP1 activity thus confirms the notion that MINPP1 can directly generate the observed new cellular InsP species from InsP_6_.

Another interesting observation, which runs counter to assumptions on MINPP1 activity,[6,25,40–42] is that the dephosphorylation sequences for InsP_6_ and InsP_5_[2OH] do not share any overlap (compare Figures S8, S9, S15, S21, S22), because MINPP1 seems to be incapable of removing the phosphoryl group at the 2-position. Likely, the charged phosphoryl group on the only axial position of the *myo-*inositol scaffold plays a role in positioning the InsPs inside MINPP1’s catalytic pocket.[43]

### MINPP1 exhibits different kinetic properties towards InsP_5_[2OH] and InsP_6_

To characterize the kinetic properties of MINPP1, we next numerically determined the reaction rates of the dephosphorylation steps from the respective experimental data based on a time-independent rates model. We formulated the kinetics of the reaction network as a Master equation and approximated the corresponding rate matrix with a least-square method that iteratively optimized the rates with respect to the scaled experimental data.[44,45] The reaction rates for the MINPP1 reaction starting with InsP_5_[2OH] as a substrate are shown in Figure 6b and were calculated from the experimental data (Figure 5a) and the corresponding network (Figure S8). The calculated rates predict progress curves (Figure 6a, S20) that are in good agreement with the experimental data which supports the assumption of time-independent rates and thus the absence of inhibition processes. The two highest reaction rates (k_20 and k_32) also correspond to the canonical MINPP1 activity towards InsP_5_[2OH] in the literature.[23,24]

**Figure 6:**
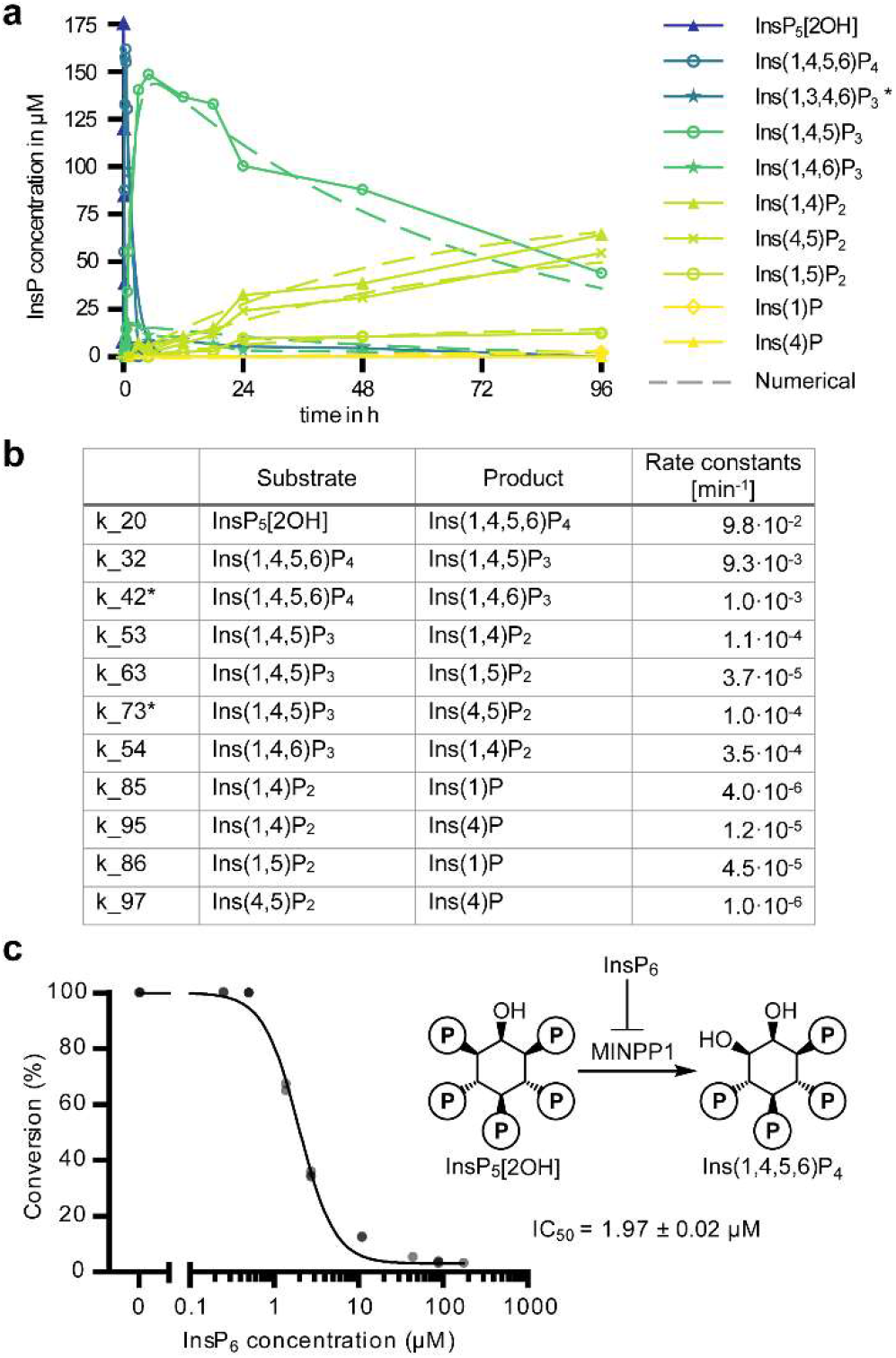
Numerical assessment of MINPP1 reaction rates. (**a**) Experimental and numerically approximated progress curves of MINPP1 dephosphorylation reactions with 175 μM InsP_5_[2OH]. Solid lines represent the experimental data (same data as in Figure 5a). The dashed lines represent the progress curves predicted by the numerically determined reaction rates. (**b**) Numerically determined reaction rates representative of two replicates. The *-marked reaction rates are subject to constraints. The SI also includes attempted numerical approximation of the MINPP1 reaction with InsP_6_. (**c**) Demonstration that InsP_6_ can inhibit dephosphorylation of InsP_5_[2OH] (175 µM) by MINPP1 (0.5 µM) with high potency.

However, in the case of InsP_6_, the computational analysis of the experimental data (Figure 5c) with the network assumption depicted in Figure S9 yielded poor results; only the consumption of InsP_6_ could be numerically analyzed with a rate of 9.3·10^−4^ min^-1^ (see SI). The poor fits indicate that the rates in the InsP_6_ dephosphorylation network might not be time-independent but are instead affected by inhibition processes that implicitly introduce a time dependence. Because of its relative stability and slow dephosphorylation, it seemed possible that InsP_6_ could act as an inhibitor for the dephosphorylation of the MINPP1-generated intermediates.[23] To test this, [^13^C_6_]InsP_5_[2OH] was incubated with MINPP1 in the presence of different amounts of [^13^C_6_]InsP_6_. Indeed, a clear inhibitory effect of InsP_6_ on the dephosphorylation of InsP_5_[2OH] by MINPP1 was observed, with an apparent IC_50_-value of 2 µM (Figure 6c).

### Ins(2,3)P_2_ and InsP_5_[3OH] are biosynthetically derived from InsP_6_ *in cellula*

With the biochemical confirmation that MINPP1 can generate InsP_5_[3OH], Ins(2,3)P_2_, and Ins(2)P *in vitro*, we sought to perform metabolic flux analysis, to confirm this reaction sequence in living cells. HEK293 or *MINPP1*^*-/-*^ HEK293 cells were labeled with [^13^C_6_]*myo-*inositol to equilibrium and subsequently exposed to medium containing 4,5[^13^C_2_]*myo-*inositol for different amounts of time before harvesting (Figure 7a). These two isotopomers were chosen to enable analysis by CE-MS: A mass difference of at least 2 Da allows the identification of the differently labeled InsPs from each other, but also the distinction of Ins(2,3)P_2_ from other, highly abundant, non-labeled sugar bisphosphates. Following cell lysis, InsP mixtures were extracted with TiO_2_ beads and analyzed *via* CE-MS to monitor the incorporation of the ^13^C_2_-isotopomers and the decrease of the ^13^C_6_-isotopomers simultaneously.

**Figure 7:**
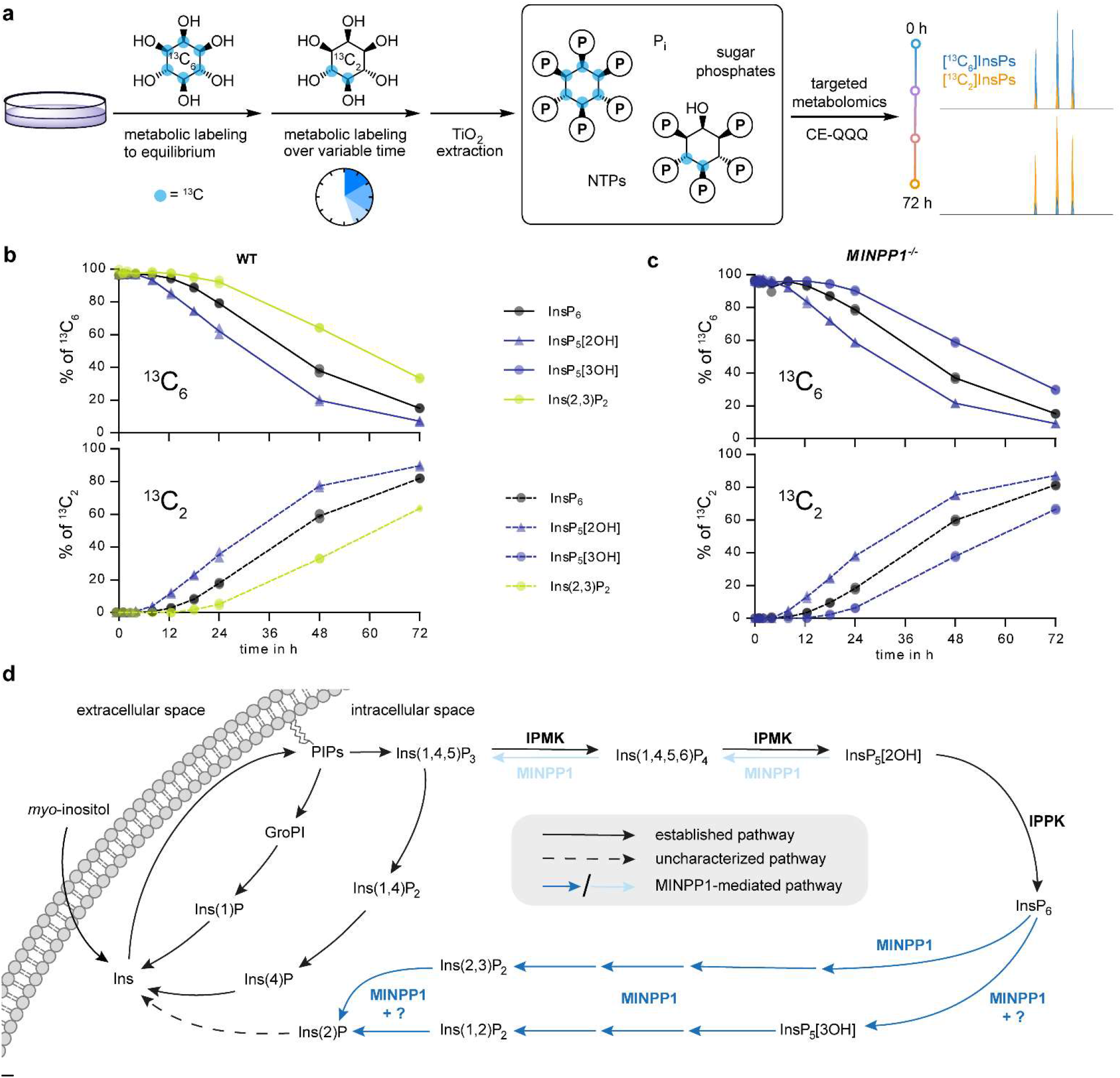
Metabolic flux analysis *via* time-dependent isotopic exchange of InsPs in HEK293 and *MINPP1*^*-/-*^ HEK293 cells. (**a**) General workflow of the metabolic flux analysis. (**b**) and (**c**) Ratios of six-fold ^13^C-labeled and doubly ^13^C-labeled InsPs HEK293 (**b**) and *MINPP1*^*-/-*^ HEK293 (**c**) cells in TiO2-extracted cell lysates. The data of two biological replicates are plotted individually and the means are connected with lines. All extracts contained a constant ∼3% of non-labeled InsP species which presumably stems from inositol neogenesis from non-labeled Glc6P (omitted, illustrated in Figure S12). (**d**): An updated overview of MINPP1-mediated InsP metabolism in human cells. As shown in this work, MINPP1 can dephosphorylate both InsP_6_ (blue arrows) and InsP_5_[2OH] (light blue arrows) *via* through two distinct, non-overlapping metabolic pathways. The question marks hint towards unknown or postulated phosphatases, that might explain the InsP_6_ 3-phosphatase activity observed in *MINPP1*^*-/-*^ cells or how Ins(2,3)P_2_ accumulates selectively in cells while both enantiomers are generated *in vitro*.

The metabolic flux analysis (Figure 7b) indicates that exogenous *myo*-inositol is incorporated first into the pool of InsP_5_[2OH], then into InsP_6_ and lastly into Ins(2,3)P_2_ (whose chemical identity was also confirmed with standards in CE-MS measurements, Figure S11). This supports the notion that Ins(2,3)P_2_ is indeed derived from InsP_6_ in human cells and not an intermediate in the biosynthesis of InsP_5_[2OH] or InsP_6_ (Figure 7d).

Interestingly, in HEK293 *MINPP1*^*-/-*^ cells, again, no Ins(2,3)P_2_ could be detected, although the sensitivity of CE-MS is superior to NMR. Thus, CE-MS analysis confirms the Ins(2,3)P_2_ dependency on MINPP1. Similarly, in the biosynthetic sequence InsP_5_[3OH] is generated after InsP_6_ (Figures 7c,d), hinting at an unidentified 3-phosphatase activity acting on InsP_6_, which has been suggested in the past.[16]

## Discussion

We have expanded the detection and identification of complex InsP mixtures using different isotopomers of *myo-* inositol, InsP_5_[2OH], and InsP_6_ in both cellular and biochemical settings. Detection *via* NMR spectroscopy provided important structural information, enabling the assignment of previously poorly characterized InsPs. Application of asymmetrically labeled 1[^13^C_1_]*myo-*inositol and 4,5[^13^C_2_]*myo-*inositol readily facilitated the distinction of enantiomers in a complex sample, which has remained an analytical challenge to this day. InsP isotopomers with different masses also proved useful tools when used in combination with CE-MS analysis, as the higher sensitivity of this technique allows for detailed metabolic flux analyses.

Taking advantage of our labeled *myo*-inositol isotopomers and InsPs, we uncovered a branch of human InsP metabolism mediated by MINPP1, which was confirmed through in-depth characterization of MINPP1’s reactivity *in vitro* and *in cellula*. The *in vitro* data illustrated that InsP_5_[2OH] is the preferred substrate for MINPP1, compared to InsP_6_. Under identical reaction conditions InsP_5_[2OH] was depleted with an apparent reaction rate that is two orders of magnitude higher than the rate for InsP_6_ (9.8·10^−2^ min^-1^ versus 9.3·10^−4^ min^-1^). Interestingly, depletion of cellular MINPP1 did not significantly alter InsP_5_[2OH] levels, suggesting that other enzymes are able to dephosphorylate InsP_5_[2OH] in a cellular setting.[46,47]

In contrast to the straightforward reaction paths for InsP_5_[2OH] dephosphorylation by MINPP1, the dephosphorylation of InsP_6_ occurs *via* an intricate network of intermediates. The first observable intermediates can be attributed to the 3-phosphatase activity of MINPP1, however, a significant portion of InsP_6_ must initially be dephosphorylated at a different position because the symmetrical Ins(1,2,3)P_3_ accumulates after ∼36h. Despite this complicated dephosphorylation network, the InsP_6_ dephosphorylation sequence converges to two final compounds, Ins(1/3,2)P_2_ and Ins(2)P *in vitro*. In all human cells we tested, Ins(2,3)P_2_ and Ins(2)P were present at significant concentrations and constitute a hitherto uncharacterized part of mammalian InsP metabolism. It was somewhat surprising, that Ins(2,3)P_2_ is the predominant InsP_2_ species within cells, given that MINPP1 is annotated as a 3-phosphatase. Our *in vitro* data demonstrate that MINPP1 is capable of producing both enantiomers, Ins(1,2)P_2_ and Ins(2,3)P_2_, *via* the aforementioned two dephosphorylation pathways from InsP_6_. It thus seems feasible that Ins(1,2)P_2_ can also be generated by MINPP1 in cells but may be depleted faster to Ins(2)P by either MINPP1 (which could be modified in its activity through posttranslational modifications or different isoforms[48]), or by a separate phosphatase altogether.

Remarkably, the many different dephosphorylation products of InsP_6_ do not overlap with any intermediates of InsP_5_[2OH] dephosphorylation, because MINPP1 appears incapable of removing the phosphoryl group at the 2-position of the inositol ring (Figure 7d). While MINPP1 converts InsP_5_[2OH] to its biosynthetic precursors Ins(1,3,4,5)P_4_ and Ins(1,4,5)P_3_ *in vitro*, InsP_6_ on the other hand is exclusively dephosphorylated to metabolites which keep the phosphoryl group at the 2-position. This data is in stark contrast to the common assumption that MINPP1 would convert InsP_6_ to InsP_5_[2OH], as is often depicted in overview schemes on InsP metabolism.[6,25,40–42] It was shown in the past that the phosphoryl group at the 2-position of the *myo-*inositol ring (the only axial position) can play an important role for proper recognition of InsPs by protein binding partners. [49,50] Our data further corroborates the importance of the phosphorylation status of the 2-position (and thus IPPK activity), because it appears that InsPs may be “sorted” into the known and reversible InsP network (when InsPs contain a free hydroxyl group at the 2-position), or InsPs enter the rather slow and potentially irreversible MINPP1-mediated circuit where they remain phosphorylated at the 2-position.

While this sorting could be accomplished solely by the preferred dephosphorylation of MINPP1, the accessibility to the two different substrates, InsP_5_[2OH] and InsP_6_ likely also plays a role. We found that the dephosphorylation of InsP_5_[2OH] was strongly inhibited by low concentrations of InsP_6_ *in vitro* (Figure 6c). In the cellular context, this potent inhibitory effect of the abundant InsP_6_ metabolite raises the question if, and how, MINPP1 can dephosphorylate InsP_5_[2OH] at all. MINPP1 would need to access localized pools of said InsP as substrates. Interestingly, MINPP1 is thought to predominantly localize to the ER, so how it accesses cytosolic (and presumably nuclear) InsPs is a question that has yet to be answered. While some studies have shown that MINPP1 (isoforms) might also be localized in cellular compartments other than the ER, or could be even secreted,[48,51,52] tools to measure intracellular concentrations of different InsPs with spatial resolution are currently not available.

Using asymmetrically isotope-labeled *myo*-inositol, it was possible to assign the uncharacterized InsP_5_ isomer that accumulates in *MINPP1*^*-/-*^ cells as InsP_5_[3OH]. This accumulation appears counter-intuitive, since MINPP1 is currently the only known enzyme in the human genome capable of generating InsP_5_[3OH]. Nevertheless, Chi *et al*. also observed a residual 3-phosphatase activity in *MINPP1*^*-/-*^ mice. [16] An analogous activity in human cells could be responsible for producing InsP_5_[3OH] from InsP_6_, as illustrated by our CE-MS-based metabolic flux analysis. Elucidating the identity of this 3-phosphatase will be of interest in the future as it constitutes an additional point of regulation within the InsP network. Furthermore, two recently reported cell lines with elevated intracellular phosphate levels were shown to contain a non-annotated InsP_5_ isomer (which we assume is also InsP_5_[1/3OH], based on the SAX-HPLC elution profiles).[53,54] Once the absolute configuration of these InsP_5_ isomers has been determined, and ideally the enzymatic activities responsible for generating these isomers, the impact of cellular phosphate homeostasis on InsP signaling could be further explored.

The physiological role of the herein described dephosphorylation pathway for InsP_6_ and its intermediates has yet to be explored. The InsPs produced by MINPP1 could be part of a recycling system converting InsP_6_ back to Ins(2)P, which might be converted to *myo*-inositol by an inositol monophosphatase (although the lithium-sensitive human enzymes IMPA1/2 are not known to act on Ins(2)P,[4,55]). In addition, it remains to be investigated which enzymes can utilize the herein identified Ins(1/3,2)P_2_ as substrates. If any of the InsP_6_-derived MINPP1 products have signaling functions themselves is also an open question. It is possible that some MINPP1-generated InsPs (or the lack thereof) could be important contributing factors in MINPP1-regulated processes, i.e. ER-stress, endochondral ossification and neuronal function. [17,19,21] For example, it would be interesting to investigate if the hyperaccumulation of InsP_5_[3OH] or the absence of Ins(2,3)P_2_ and Ins(2)P are partially responsible for causing pontocerebellar hypoplasia (PCH) in patients with MINPP1 loss-of-function mutations.[20,21] Ucuncu *et al*. proposed that hyperaccumulation of InsP_6_ in neuronal cells of PCH patients might be a mechanistic cause of this disease.[21] In contrast to their reported 3-to 4-fold increase of [^3^H]InsP_6_ levels (normalized against total tritiated PIPs) in HEK293 *MINPP1*^*-/-*^ cells compared to WT cells, we only observed a slight increase, using the same cell line but normalizing against packed cell volume. This discrepancy points towards several interesting possibilities: a) PIP levels, or the incorporation of exogenous *myo-*inositol, could be (indirectly) influenced by MINPP1 activity, b) radioactivity-induced cell stress could have an effect on MINPP1 expression [17] or c) knockout of MINPP1 changes the cell shape/volume. To differentiate between these possibilities, different quantification methods (e. g. normalization against total protein or DNA concentration) should be compared in the future, and the composition of PIP isotopomers during metabolic labeling experiments could be probed with mass spectrometry-based methods. [56–58]

As a next step, the combination of inositol isotopomers, NMR and CE-MS we used in this study could be useful to probe InsP metabolism in a variety of biological contexts. E.g. it could be investigated how the InsP pool changes during ER-related stress, during which MINPP1 is upregulated, and how this might correlate with the onset of apoptosis.[17] Other applications could be to determine the fate of inositol (phosphates) in pathogenic parasites such as *T. cruzi*, in which InsP metabolism is essential for the developmental cycle.[59] The question if, or how, the InsP metabolism of the host cell and the parasite have an influence on each other might lead to new therapeutic avenues for these parasitoses. It should be noted that our approach currently only probes the InsP pool generated from exogenously incorporated *myo-*inositol. However, in a low-inositol environment mammalian cells produce *myo-* inositol (and subsequently InsPs) *de novo* from glucose-6-phosphate, which depends strongly on ITPK1 (inositol tetrakisphosphate 1-kinase) activity.[53] To also include glucose-derived InsPs in the overall analysis, a different experimental setup would be required. The dissection of InsP degradation in an extracellular context, namely how InsPs contained in food are converted by digestive processes or the gut microbiome and if the resulting metabolites might have beneficial or detrimental effects on health, are also fascinating questions.[43,60,61] With the tools and methods reported here these topics now become addressable.

## Methods

All experimental and computational methods are described in the Supplementary Information.

## Supporting information

Supplementary file SI_main

Supplementary file SI_Numerical_Analysis_of_MINPP1-mediated_dephosphorylation

## Supplementary Information

Supplementary Information file “SI_Main” contains supplementary figures, methods for all *in vitro* and cellular experiments and NMR spectra.

Supplementary Information file “SI_Numerical_Analysis_of_MINPP1-mediated_dephosphorylation” contains methods and additional figures for the computational part of this work.

## Acknowledgments

M.N.T. and S.K. were funded by the Deutsche Forschungsgemeinschaft (DFG, German Research Foundation) under Germany’s Excellence Strategy – EXC 2008 – 390540038” – UniSysCat.

We thank Peter Schmieder for providing valuable guidance with NMR topics, Lena von Oertzen and Kathrin Motzny for their assistance in cell culture matters, Robert Puschmann for initial synthesis of *myo*-inositols and all members of the Fiedler lab for proof-reading.

## Author contributions

M.N.T. performed most biochemical and *in vitro* experiments, data analyses, and organic syntheses. S.K. performed all kinetic modeling analyses under supervision of B.K.. Z.F. contributed to initial testing of MINPP1 expression and reactivity optimizations. D.Q. and G.L. measured all CE-MS samples under supervision of H.J.. D.F. and M.N.T. conceived the project with input from A.S.. D.F. and M.N.T. supervised experimental work by Z.F.. M.N.T. and D.F. prepared the initial draft of the paper with S.K. contributing the kinetic modeling part and all authors contributed to the final version of the manuscript.

