## Supplementary file SI_main for "Stable isotopomers of *myo*-inositol to uncover the complex MINPP1-dependent inositol phosphate network"

[1]: Leibniz-Forschungsinstitut für Molekulare Pharmakologie, Robert-Rössle-Straße 10, 13125 Berlin, Germany

[2]: Institut für Chemie, Humboldt-Universität zu Berlin, Brook-Taylor-Straße 2, 12489 Berlin, Germany

[3]: Institut für Chemie und Biochemie, Freie Universität Berlin, Arnimallee 22, 14195 Berlin, Germany

[4]: Institut für Organische Chemie, Albert-Ludwigs-Universität Freiburg, Albertstraße 21, 79104 Freiburg, Germany

[5]: MRC Laboratory for Molecular Cell Biology, University College London, WC1E 6BT London, UK

### Abbreviations

| | |
|---|---|
| ACN | acetonitrile |
| BIRD | bilinear rotation decoupling |
| BIRD-HMQC | HMQC with BIRD pulse |
| BPG | 2,3-bisphoshpoglycerate |
| CD | circular dichroism spectroscopy |
| CE-MS | capillary electrophoresis electrospray mass spectrometry |
| DCI | 4,5-dicyanomidazole |
| DCM | dichloromethane |
| DMEM | Dulbecco's Modified Eagle Medium |
| DMSO | dimethylsulfoxide |
| DTT | dithiothreitol |
| EDTA | ethylenediaminetetraacetic acid |
| FBS | fetal bovine serum |
| GndHCl | guanidium hydrochloride |
| HMQC | heteronuclear multiple-quantum correlation |
| Ins | *myo*-inositol |
| InsPx | inositol phosphate |
| IPS | inositol phosphate synthase |
| IPTG | isopropyl *β*-D-1-thiogalactopyranoside |
| MINPP1 | multiple inositol polyphosphate phosphatase 1 |
| MWCO | molecular weight cut-off |
| $NAD^+$ | nicotinamide adenine dinucleotide (oxidized form) |
| NMR | nuclear magnetic resonance (spectroscopy) |
| $OD_{600}$ | optical density (at 600 nm) |
| ORF | open reading frame |
| PCV | packed cell volume |
| ppm | parts per million |
| rt | room temperature |
| SDS-PAGE | sodium dodecyl sulfate polyacrylamide gel electrophoresis |
| TB | terrific broth |
| TMPBr | tetramethylphosphonium bromide |

### Supporting Figures and Tables

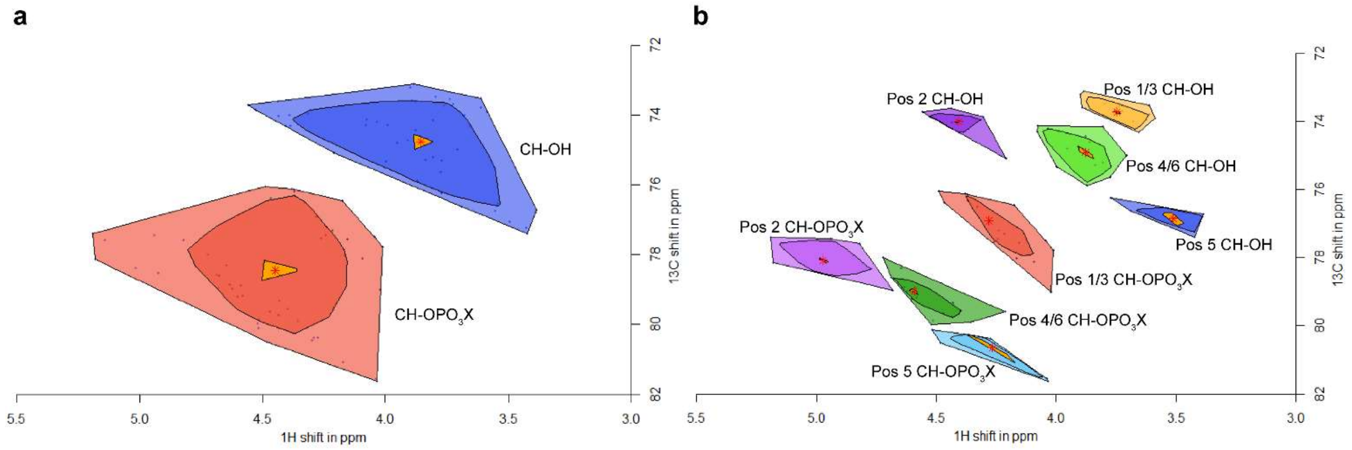

Figure S1: The bagplots illustrate the clustering depending on phosphororylation state (**a**) (blue: OH groups, red: phosphorylated groups) and position on the inositol ring (**b**). A fence factor of 6 was used to include all data points in the respective bags, the underlying data is the same as shown in Figure 2.

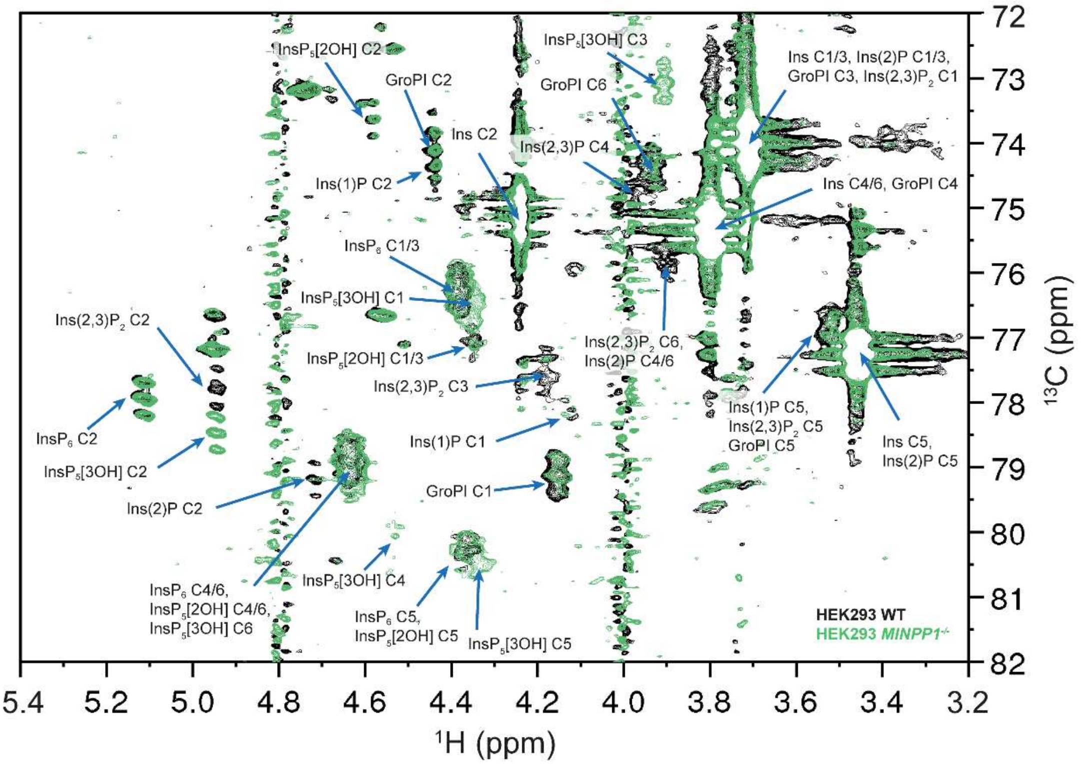

Figure S2: Complete annotation of [$^{13}C_6$]Ins-labeled HEK293 WT (black spectrum) and *MINPP1*$^{-/-}$ (green spectrum) metabolic extracts.

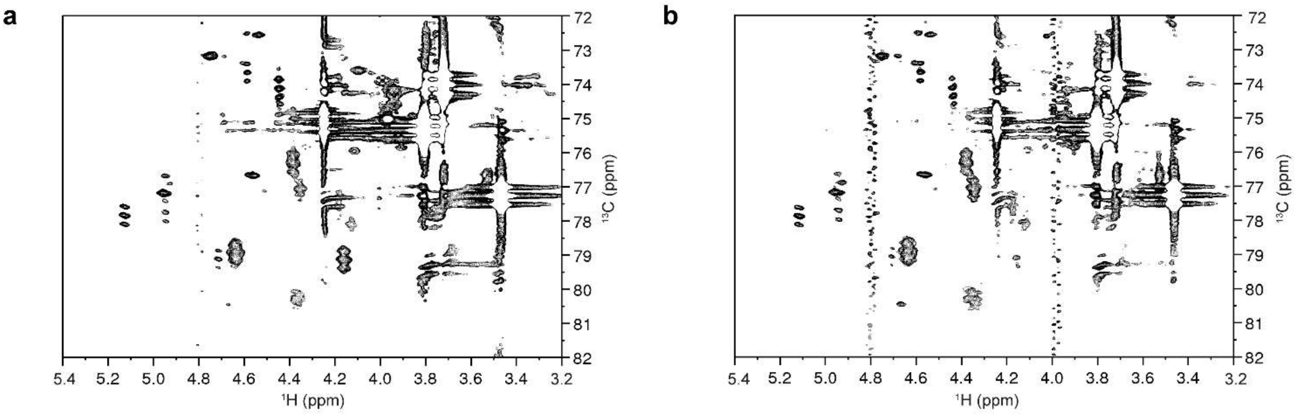

Figure S3: Additional NMR spectra of metabolic extracts from immortalized human wild-type cells. (**a**): HT29, (**b**): H1975. All labeled wild-type cells lines contain the same set of InsPs with varying concentrations.

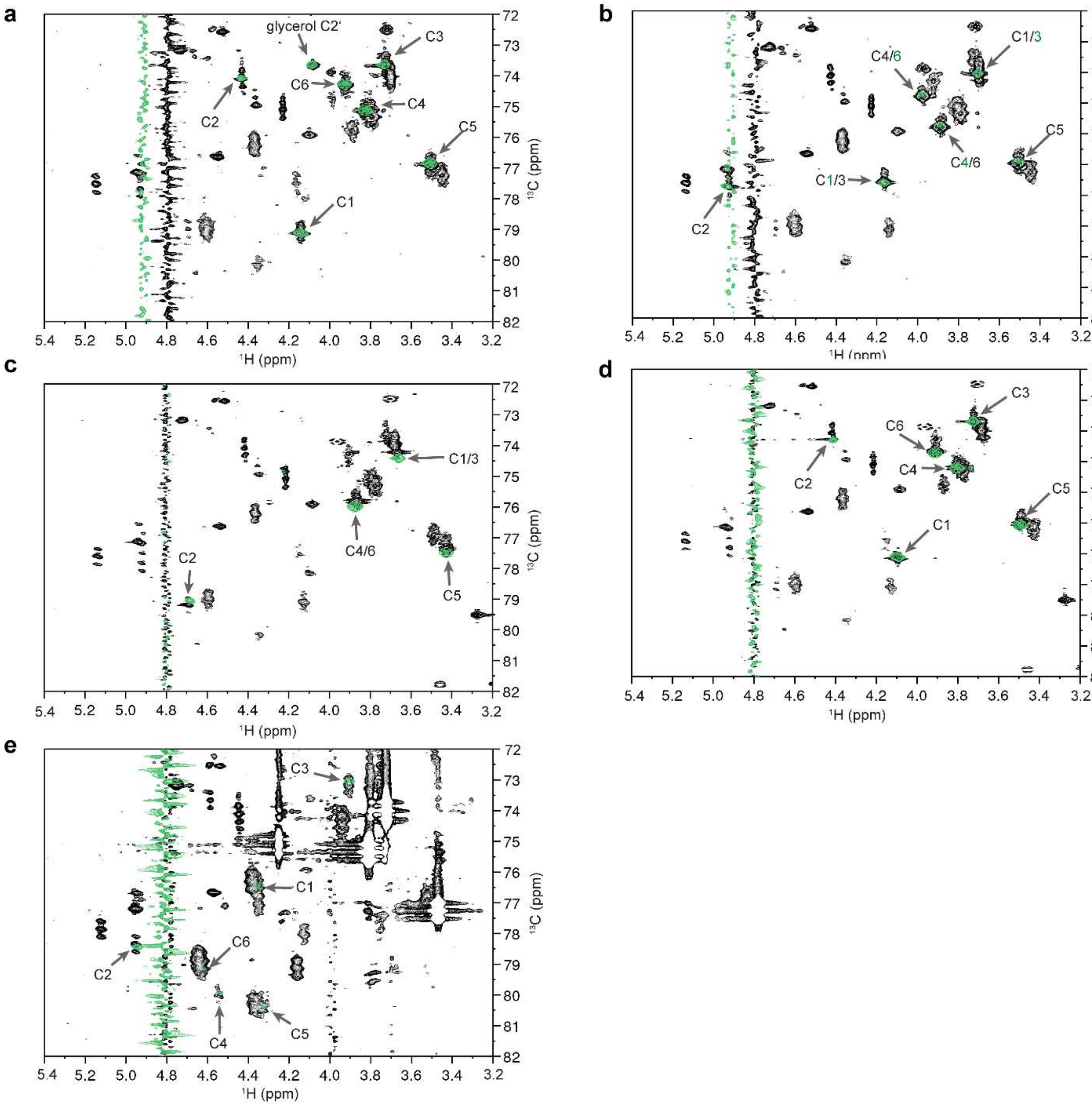

Figure S4: NMR spectra of [$^{13}C_6$]Ins-labeled H1Hela WT metabolic extracts spiked with InsP standards (black) overlayed with spectra of the same standard in saturated $KClO_4$ solution in $D_2O$, pH* = 6.0 (green). a: GroPI; b: Ins(1,2)$P_2$; c: Ins(2)P, d: Ins(1)P. e: NMR spectrum of [$^{13}C_6$]Ins-labeled HEK293 *MINPP1*$^{-/-}$ metabolic extract (black) overlayed with $InsP_5$[3OH] (red). The corresponding positions on the inositol ring are annotated with arrows. For Ins(1,2)$P_2$ the annotations for the spike-in standard are written in green while the annotation for the other enantiomer Ins(2,3)$P_2$, which is the species present in mammalian cells, are written in black. Note that in a and b the solvent signal is shifted between the extract and the InsP standards due to different sample temperatures during NMR measurement.

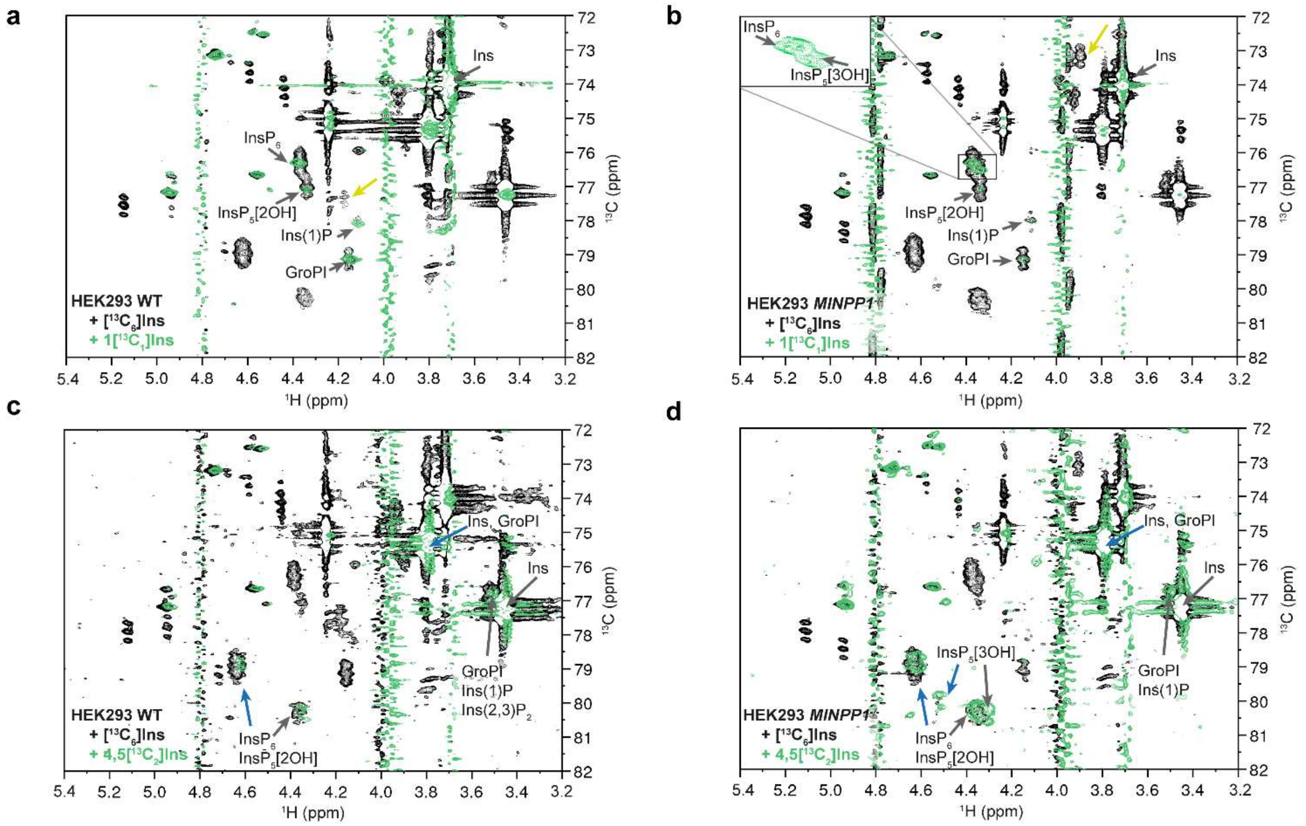

Figure S5: HMQC spectra of HEK cell lines which were metabolically labeled with asymmetrical isotopomers of *myo*-inositol (green) overlayed on the respective spectra of $[^{13}C_6]$*myo*-inositol-labeled cells. Annotations were limited to labeled positions. (a): $1[^{13}C_1]$*myo*-inositol-labeled HEK293 WT cells. This labeling experiment illustrates that the signal for the phosphorylated position of Ins(1/3,2)$P_2$ (yellow arrow) is not labeled, thus excluding Ins(1,2)$P_2$ as a possible enantiomer. Note that non-labeled positions of *myo*-inositol are still visible due to the large relative abundance of *myo*-inositol in the extract. (b): $1[^{13}C_1]$*myo*-inositol-labeled HEK293 *MINPP1*$^{-/-}$ cells. Here the signal for the 1-position of $InsP_5$[1/3OH] is clearly still phosphorylated (insert showing the magnified region), while the dephosphorylated 1/3-position (yellow arrow) is not labeled, indicating that the enantiomer present cannot be $InsP_5$[1OH], but must be $InsP_5$[3OH]. (c): $4,5[^{13}C_2]$*myo*-inositol-labeled HEK293 WT cells. For $4,5[^{13}C_2]$*myo*-inositol-labeled spectra the 4-positions are marked with a blue arrow and 5-positions with a black arrow. Note that the InsP signals now show a characteristic doublet pattern due to $^{13}C$-$^{13}C$ coupling. (d): $4,5[^{13}C_2]$*myo*-inositol-labeled HEK293 WT *MINPP1*$^{-/-}$ cells. Here, the signal for the 4-position of $InsP_5$[3OH] is shifted away from the signals of the 4/6-positions of $InsP_6$/$InsP_5$[2OH], which is consistent with the observed shifts for the $InsP_5$[3OH] standard, but not $InsP_5$[1OH] (see also Figure S4e].

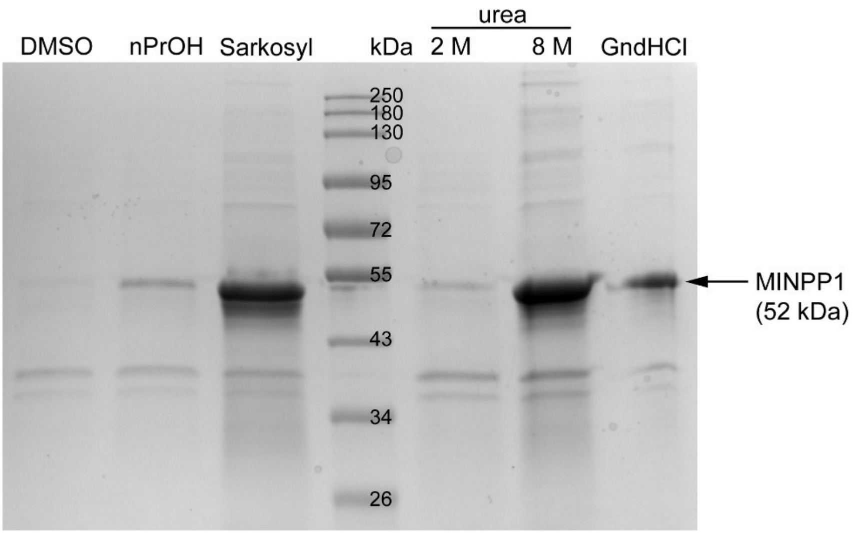

Figure S6: Testing resolubilization buffers for MINPP1 purification from inclusion bodies. MINPP1 was expressed according to the procedure described in the Experimental section. One part of the cell debris pellet obtained after lysis was washed only once with DI water, weighted, resuspended in little water and distributed into six 15 mL tubes (110 mg of wet pellet per tube). 4 mL of each resolubilization buffer (Experimental section under Cloning and production of MINPP1) were added to each tube, and incubated for 16 h at 4 °C on a reciprocal shaker. The tubes were centrifuged (30 min, 3000 g, 4 °C). 10 µL of supernatant were each diluted with 60 µL deionized water, 30 µL SDS running buffer, 40 µL Lämmli-buffer (incl. *β*-mercaptothanol) and all samples except for the guanidinium hydrochloride-based sample were boiled for 5 min at 90 °C. 30 µL of each sample were loaded on an SDS-PAGE gel, 150 V were applied until the loading marker completely ran into the gel. The wells were then flushed with SDS running buffer to remove excess guanidium hydrochloride to prevent gel distortions. Then the SDS-PAGE was continued (150 V, 45 min) and stained using colloidal Coomassie.

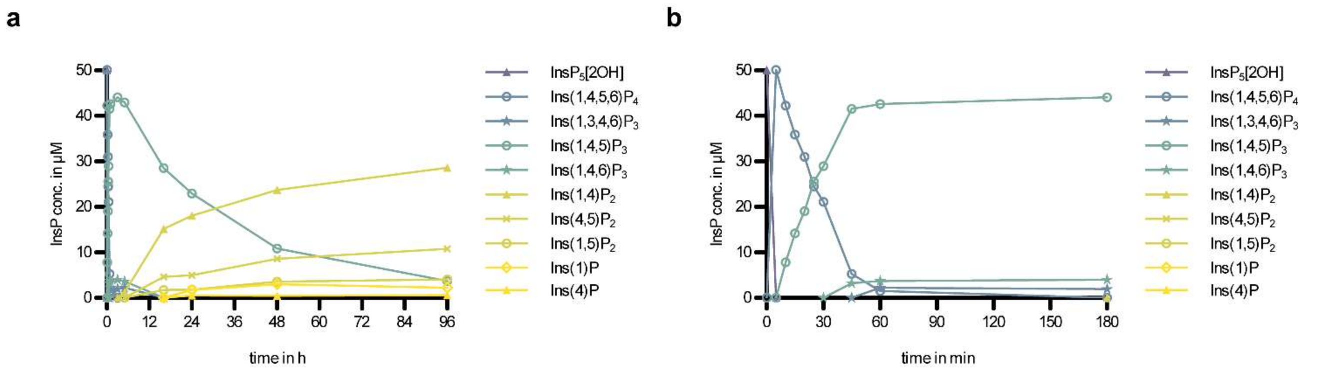

Figure S7 Progress curves of MINPP1 reaction with 50 µM $InsP_5$[2OH] (a and the first 180 min in b).

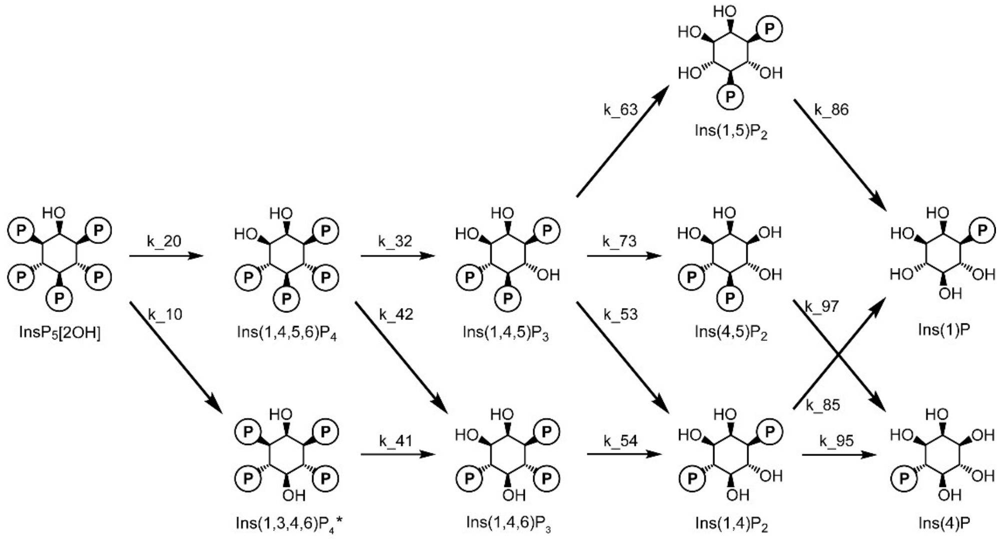

Figure S8: Complete MINPP1-mediated dephosphorylation pathway observed for $InsP_5$[2OH]

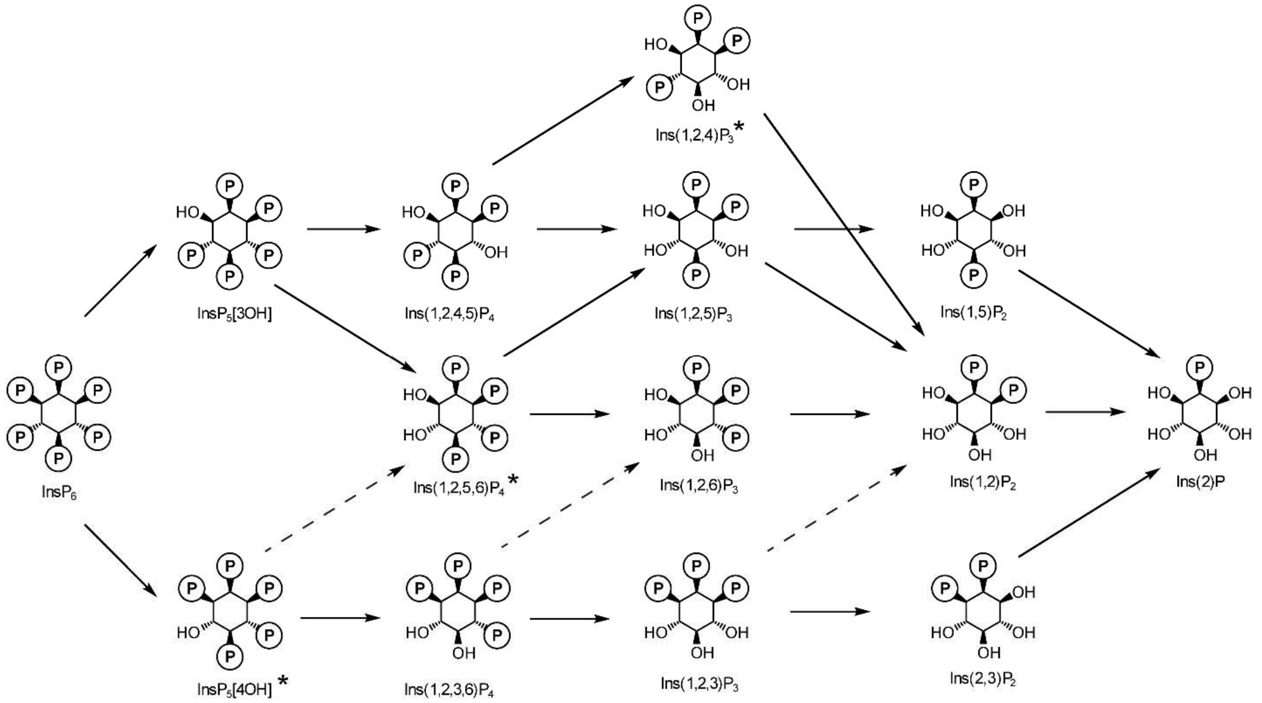

Figure S9: Complete MINPP1-mediated dephosphorylation pathway observed for $InsP_6$. The dashed arrows indicate theoretically possible paths which we assume are not relevant to the overall outcome. More investigation is needed to confirm this.

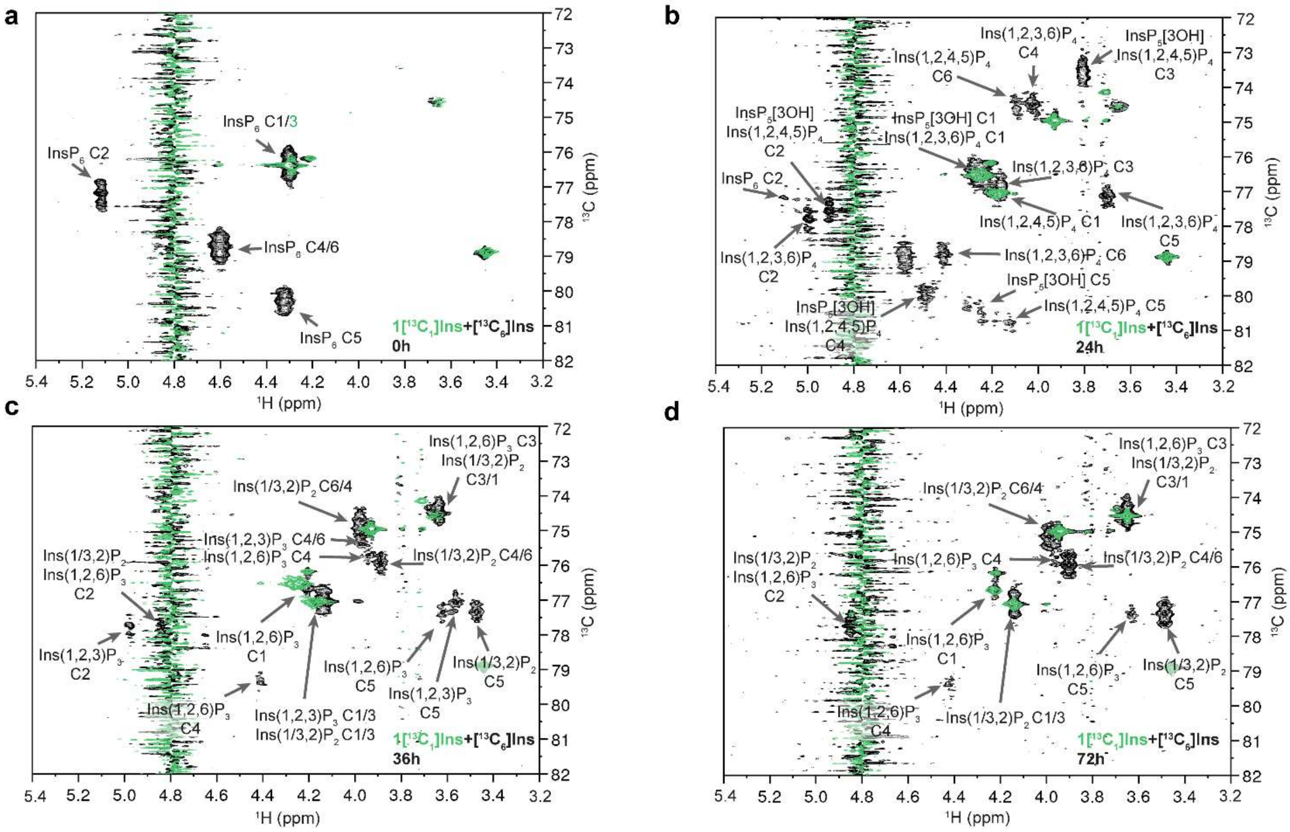

Figure S10: MINPP1 dephosphorylation of 1[$^{13}C_1$]$InsP_6$. The spectra show reactions in which MINPP1 was incubated with either 175 µM 1[$^{13}C_1$]$InsP_6$ (green) or a 1:1 mixture of [$^{13}C_6$]Ins:1[$^{13}C_1$]Ins. (**a**): Control sample without enzyme. $InsP_6$ is clearly shown to be labeled at the 1-position. The other visible signals belong to buffer components. (**b**): Reaction mixture after 24 h of incubation. The 1-position is not dephosphorylated at this stage. Thus, the shown enantiomers are enantiopure. The 3-position of Ins(1,2,3,6)$P_4$ is slightly shifted upfield with regards to the $^{13}$C-dimension, compared to the labeled 1-position of Ins(1,2,4,5)$P_4$. Also, the signals at ~75 and ~79 ppm ($^{13}$C dimension) are buffer components from the MINPP1 stock solution. (**c**): Reaction mixture after 36 h incubation. The 1-position of Ins(1,2,6)$P_3$ and 3/1-position of Ins(1/3,2)$P_2$ overlap with the buffer component at ~75 ppm which seems to increase in intensity. The labeling of the 1-position appearing in both the region for phosphorylated and the region for dephosphorylated positions indicate that a mixture of Ins(2,3)$P_2$ and Ins(1,2)$P_2$ is formed. (**d**): Reaction mixture after 72 h of incubation. The dephosphorylated 1-position of Ins(2,3)$P_2$ is now evident while the 1-position of Ins(1,2)$P_2$ is still phosphorylated, indicating that a mix of both enantiomers has been formed, despite the enantio-specific nature of the previous dephosphorylation steps in (b). A rough integration of all labeled 1-position signals (green spectrum) resulted in a near 1:1 ratio between the dephosphorylated 1-position of Ins(2,3)$P_2$ and the combined phosphorylated 1-position of Ins(1,2)$P_2$ and Ins(1,2,6)$P_3$. This high ratio suggests that MINPP1 likely converts Ins(1,2,3)$P_2$ likely exclusively into Ins(2,3)$P_2$, while all intermediates downstream of $InsP_5$[3OH] must result in the other enantiomer Ins(1,2)$P_2$.

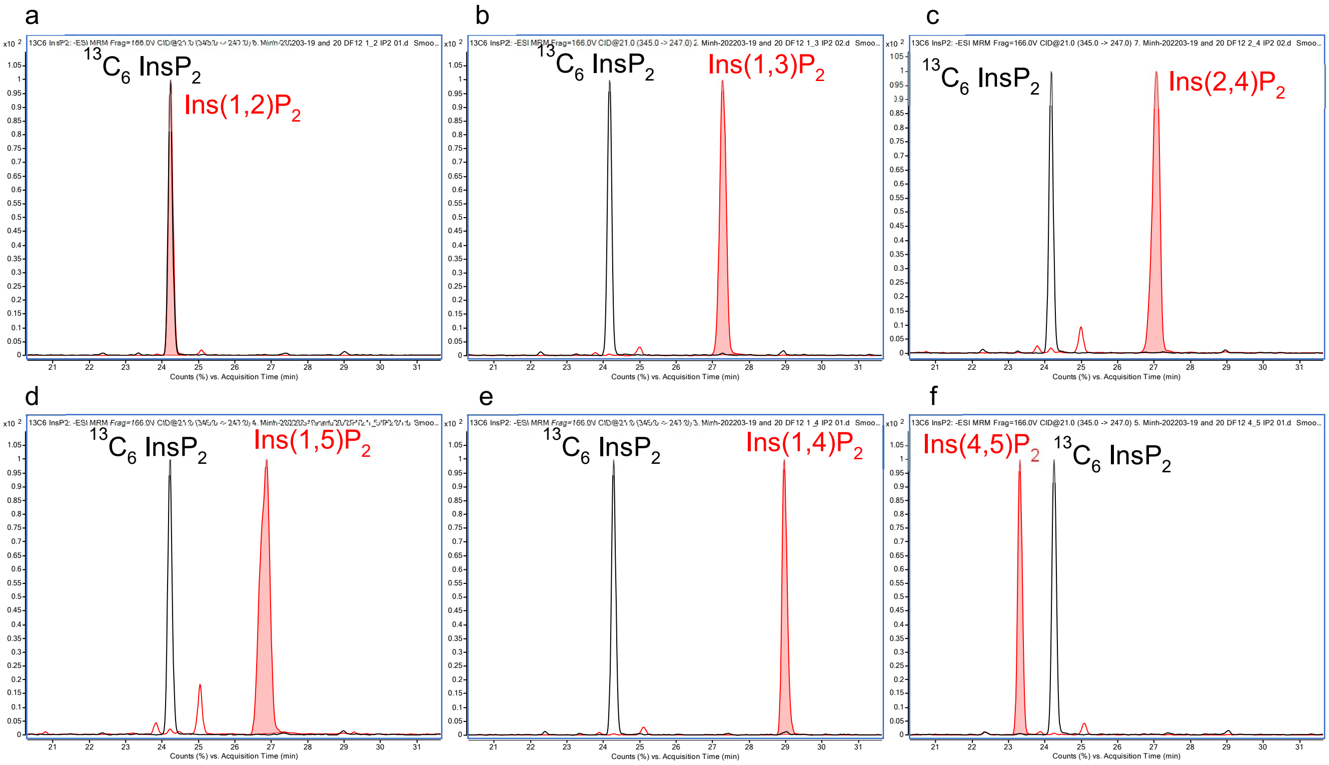

Figure S11: Confirmation of the identity of Ins(1/3,2)$P_2$ *via* CE-MS. The metabolic extract of [$^{13}C_6$]Ins metabolically-labeled HEK293 WT cells were spiked with commercial standards of different $InsP_2$ isomers and analyzed *via* CE-MS. Depicted are the extracted ion chromatograms corresponding to the masses of the intracellularly synthesized [$^{13}C_6$]$InsP_2$ (black) and the non-labeled $InsP_2$ standards. Only Ins(1,2)$P_2$ coelutes with the [$^{13}C_6$]$InsP_2$ signal in question (a) while all other tested $InsP_2$ standards (b: Ins(1,3)$P_2$, c: Ins(2,4)$P_2$, d: Ins(1,5)$P_2$, e: Ins(1,4)$P_2$, f: Ins(4,5)$P_2$) do not.

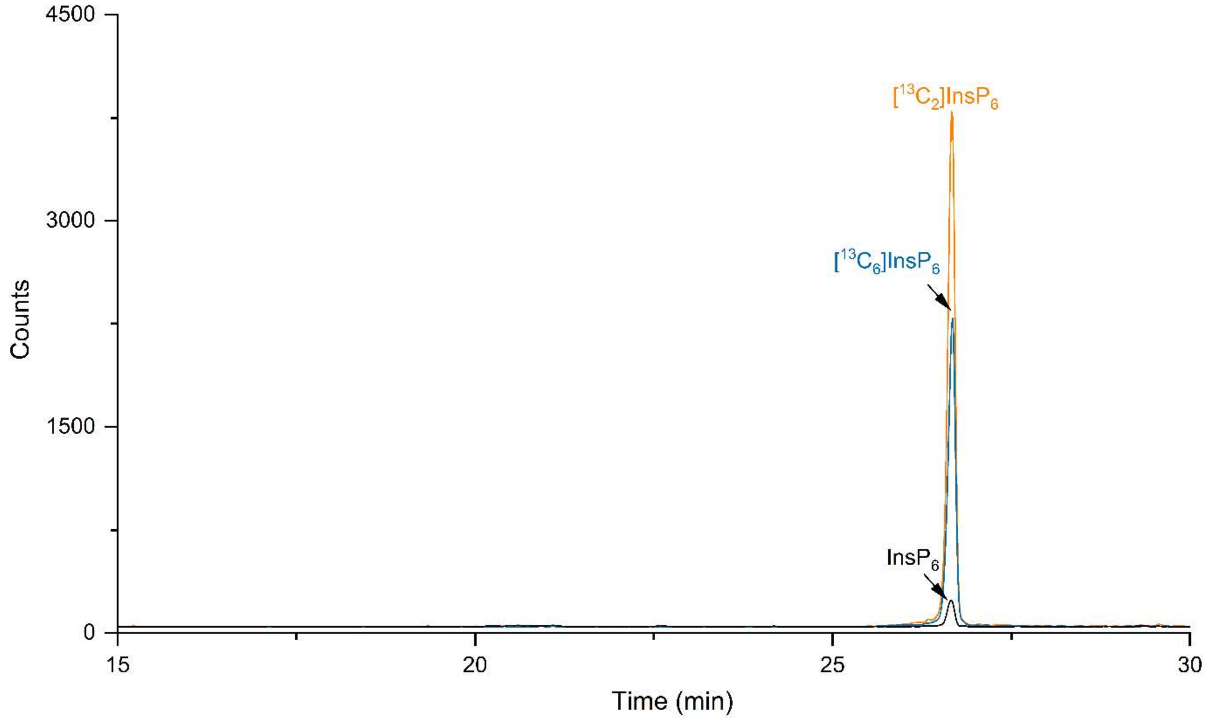

Figure S12: Example EICs (extracted ion chromatograms) of [13$C_{6/2/0}$]Ins$P_6$ in a HEK293 WT cells which were metabolically labeled with [$^{13}C_6$]*myo*-inositol to equilibrium and then with 4,5[$^{13}C_2$] *myo*-inositol for 48h. For the metabolic flux analysis (Figures 7b, 7c) the integrals of the respective isotopomer peaks (blue/ orange) were used for relative quantification. The InsP pools contained a constant ~3% of non-labeled InsPs (black) due to glucose-6-phosphate-dependent neogenesis of *myo*-inositol.

### Experimental section

#### General Information

Chemicals were obtained from Sigma Aldrich, VWR, Roth, TCI, Thermo Scientific or Roche and used without further purification unless stated otherwise.

InsP standards were purchased as sodium, potassium, ammonium or cyclohexylammonium salts from SiChem (Ins(3,4,5,6)$P_4$, Ins(1,4,5,6)$P_4$, Ins(1,4,5)$P_3$, Ins(1,3,4)$P_3$, $InsP_5$[3OH], $InsP_5$[1OH]), Cayman chemical (Ins(2,3,5)$P_3$, Ins(1,2)$P_2$), Echelon Bioscience (Ins(1,4)$P_2$, Ins(1,2,6)$P_3$, GroPI), Biomol (Ins(1,5)$P_2$, Ins(1,4,6)$P_2$) or Sigma-Aldrich (Ins(1)P, Ins(2)P) or synthesized in-lab ([$^{13}C_6$]$InsP_6$, [$^{13}C_6$]$InsP_5$[2OH], [$^{13}C_6$]1PP-$InsP_5$, [$^{13}C_6$]5PP-$InsP_5$, [$^{13}C_6$]1,5$(PP)_2InsP_4$) as described previously.[1] Non-labeled InsPs were dissolved in a saturated $KClO_4$ solution in $D_2O$ (pH* 6.0) to mimick the conditions of the metabolic extracts. Non-labeled standards were dissolved in the smallest volume possible for NMR measurements (min. 500 µL). All samples were adjusted to pH* 6.0 if necessary using DCl and NaOD solutions in $D_2O$ (all deuterated solutions obtained from Eurisotop).

For NMR-based quantification purposes standards (TMPBr (Sigma, 288268) or phosphonoacetic acid (TraceCert $^{31}$P-NMR standard, Supelco, 79251), respectively) were dissolved/ diluted in dry $D_2O$ (Eurisotop D215T) and aliquots are frozen until use.

#### NMR data acquisition and processing

For NMR measurements and NMR data analysis TopSpin 3.5 was used. Measurements were conducted on a Bruker AV-III spectrometer (Bruker Biospin, Rheinstetten, Germany) operating at 600 MHz for $^1$H and 151 MHz for $^{13}$C nuclei equipped with a cryo-QCI probe. The pulse sequence for BIRD-{$^1$H,$^{13}$C}HMQC is based on the hmqcbiph pulse program from Bruker. Measurement parameters are adapted depending on sample composition. Typically, metabolic extracts were recorded with TD($^{13}$C) = 1024, 140 scans, spectral width ($^{13}$C) limited to 40 – 100 ppm. Typically, samples from *in vitro* experiments were recorded with TD ($^{13}$C) = 512, 64 scans, spectral width ($^{13}$C) limited to 50 – 90 ppm. All samples were recorded at 310 K.

BIRD-{$^1$H,$^{13}$C}HMQC-NMR spectra were processed without digital water suppression with manual phasing and automatic baseline correction.

Quantification of NMR data were conducted as follows: For metabolic extracts InsPs were quantified against a known concentration of tetramethylphosphonium bromide (TMPBr). A standard curve for $InsP_6$ and $InsP_5$[2OH] against TMPBr was recorded earlier [2]. For other InsP species the standard curve for $InsP_6$ was used as an approximation as there are no fully $^{13}$C-labeled standards available. For the samples from the *in vitro* dephosphorylation of $InsP_{6/5}$ by MINPP1 the InsP signals were quantified relatively to each other and normalized to a total InsP concentration matching the initial substrate concentration. As the signals of the 2-positions are the sharpest and best resolved (due to the reduced coupling to the neighbouring CH groups), the 2-position signals were used for quantification. In the cases where the 2-position signals of two InsPs species are not baseline-separated, the signals were integrated together and split by the ratio of the 5-position signal integrals.

#### CE-MS measurement

CE-ESI-MS has been found to be an efficient platform for the analysis of inositol polyphosphate.[3] A CE-ESI-QQQ setup is used for this study, which consists of an Agilent 7100 CE, a triple quadrupole tandem mass spectrometry Agilent 6495c, connected to an Agilent Jet Stream (AJS) electrospray ionization (ESI) source. A commercial CE-MS sheath liquid coaxial interface was used, with an isocratic LC pump constantly delivering the sheath-liquid (*via* a splitter set with a ratio of 1:100). All experiments were performed on a bare fused silica capillary with a length of 100 cm (50 µm internal diameter and 365 µm outer diameter). 35 mM ammonium acetate titrated by ammonia solution to pH 9.7 was employed as background eletrolyte (BGE). Samples were injected by applying 100 mbar pressure for 15 s, corresponding to 1.5% of the total capillary volume (30 nL).

The sheath liquid is a mixture of water-isopropanol (1/1, v/v) and with a constant flow of 10 µL/min. The MS source parameters settings were as follows: nebulizer pressure was set to 8 psi, gas temperature was 150 °C with a flow of 11 L/min, sheath gas temperature was 175 °C and with a flow of 8 L/min, capillary voltage was -2000 V with nozzle voltage 2000 V. Negative high-pressure RF and low-pressure RF (Ion Funnel parameters) were 70 V and 40 V, respectively. Mass spectrometer parameters for MRM transitions are shown below.

| **Compound Name** | **Precursor Ion** | **Product Ion** | **dwell** | **Frag (V)** | **CE (V)** | **Cell Acc (V)** | **Polarity** |
|---|---|---|---|---|---|---|---|
| [$^{13}C_6$]InsP$_6$ | 331.9 | 486.9 | 60 | 166 | 13 | 4 | Negative |
| [$^{13}C_2$]InsP$_6$ | 329.9 | 482.9 | 60 | 166 | 13 | 4 | Negative |
| [$^{12}C_6$]InsP$_6$ | 328.9 | 480.9 | 60 | 166 | 13 | 4 | Negative |
| [$^{13}C_6$]InsP$_5$ | 292 | 504.9 | 60 | 166 | 9 | 3 | Negative |
| [$^{13}C_2$]InsP$_5$ | 290 | 500.9 | 60 | 166 | 9 | 3 | Negative |
| [$^{12}C_6$]InsP$_5$ | 289 | 498.9 | 60 | 166 | 9 | 3 | Negative |
| [$^{13}C_6$]InsP$_4$ | 252 | 424.9 | 60 | 166 | 5 | 1 | Negative |
| [$^{13}C_2$]InsP$_4$ | 250 | 420.9 | 60 | 166 | 5 | 1 | Negative |
| [$^{12}C_6$]InsP$_4$ | 249 | 418.9 | 60 | 166 | 5 | 1 | Negative |
| [$^{13}C_6$]InsP$_2$ | 345 | 247 | 60 | 166 | 21 | 4 | Negative |
| [$^{13}C_2$]InsP$_2$ | 341 | 243 | 60 | 166 | 21 | 4 | Negative |
| [$^{12}C_6$]InsP$_2$ | 339 | 241 | 60 | 166 | 21 | 4 | Negative |

#### Data handling

For plotting and other analyses Microsoft Excel, OriginPro 2016 and GraphPad Prism 5 were used. Bagplots were created in R (version 4.1.2) with the aplpack package (version 1.3.5).

For details of the kinetic modelling of MINPP1 see separate SI file “SI_Numerical_Analysis_of_MINPP1-mediated_dephosphorylation”.

#### Synthesis of $^{13}C$-labeled Ins and InsPs

The synthesis of $^{13}C$-labeled *myo*-inositol and its derivatization to InsPs were carried out based on published procedures for [$^{13}C_6$]Ins with slight improvements of the protocol as described below.[2]

#### Chemoenzymatic synthesis of *myo*-inositol isotopomers

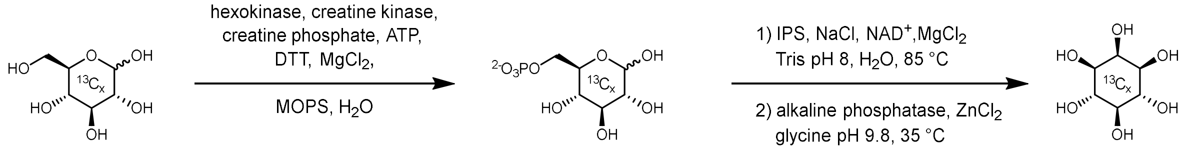

4[$^{13}C_1$]D-glucose (**S1a**)
1,2[$^{13}C_2$]D-glucose (**S1b**)
[$^{13}C_6$]D-glucose (**S1c**)

4[$^{13}C_1$]D-glucose-6-phosphate (**S2a**)
1,2[$^{13}C_2$]D-glucose-6-phosphate (**S2b**)
[$^{13}C_6$]D-glucose-6-phosphate (**S2c**)

1[$^{13}C_1$]Ins (**S3a**)
4,5[$^{13}C_2$]Ins (**S3b**)
[$^{13}C_6$]Ins (**S3c**)
(crude)

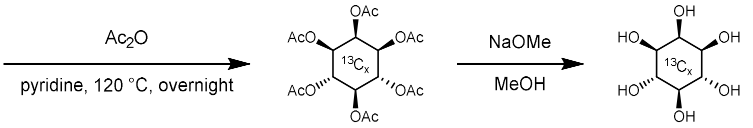

1[$^{13}C_1$]Ins hexakisacetate (**S4a**)
4,5[$^{13}C_2$]Ins hexakisacetate (**S4b**)
[$^{13}C_6$]Ins hexakisacetate (**S4c**)

1[$^{13}C_1$]Ins (**S3a**)
4,5[$^{13}C_2$]Ins (**S3b**)
[$^{13}C_6$]Ins (**S3c**)

Ins isotopomers were synthesized chemoenzymatically from the respective D-glucose isotopomer: 1[$^{13}C_1$]Ins (**S3a**) was synthesized from 4[$^{13}C_1$]D-glucose (**S1a**), 4,5[$^{13}C_2$]Ins (**S3b**) was synthesized by starting from 1,2[$^{13}C_2$]D-glucose (**S1b**), and [$^{13}C_6$]Ins (**S3c**) from [$^{13}C_6$]D-glucose (**S1c**). $^{13}$C-labeled material was obtained from Eurisotop/ Cambridge Isotope Labs. Generally, we observed improved yields with higher synthesis scale with 1 to 3 g glucose as starting material yielding up to 55% Ins. However, the asymmetric isotopomer **S3a** was synthesized only on a 500 mg scale.

Briefly, **S1a/b/c** is first converted enzymatically to the respective D-glucose-6-phosphate (**S2a/b/c**) with hexokinase and crudely purified *via* an anion exchange hand column. The subsequent lyophilization step of the eluate in the original procedure can be replaced by concentrating using a rotavap without reduction of yield while saving time. The resulting product/salt mixture is then converted to inositol-3-monophosphate (Ins(3)P) through the action of inositol monophosphate synthase (IPS), which is monitored *via* NMR. We recommend preparing recombinantly expressed IPS as closely to the protocol in [2] as possible to ensure sufficient activity of the IPS (esp. induction at high $OD_{600}$ and purification via heat-treatment); prolonged reaction times causes the $NAD^+$ cofactor to degrade, inhibiting IPS activity even after resupplementing more IPS and $NAD^+$. Subsequently. Ins(3)P is dephosphorylated to Ins (**S3a/b/c**) by alkaline phosphatase. The reaction progress is also monitored *via* NMR. The ion exchange treatment in the original procedure can be skipped upon complete conversion and the aqueous solution can be reduced on a rotavap instead, yielding a crude brown solid. The Ins is then purified through chemical derivatization by acetylation to *myo*-inositol hexakisacetate (**S4a/b/c**), purification *via* extraction and column chromatography on silica gel (~500 mL silica gel for a 3 g synthesis scale), followed by deacetylation and precipitation in acetonitrile to afford the desired *myo*-inositol isotopomer **S3a/b/c** in pure form following the published protocol.

**<u>1[$^{13}C_1$]Ins (S3a)</u>**: yield: 122 mg (starting from 500 mg **S1a**, 24%)

**$^1$H NMR** (600 MHz, $D_2O$) δ[ppm]: 3.99 (s, 1H, 2-position), 3.56 (ps-q, J = 9.8 Hz, 2.5H, 4/6-position and 1-position), 3.46 (d, J = 9.9 Hz, 1H, 3-position), 3.34 (d, J = 11.4 Hz, 0.5H, 1-position), 3.21 (t, J = 9.5 Hz, 1H, 5-position). Please note that the 1-position is coupling with $^{13}$C with a coupling constant of $^1J_{CH}$ = 143.4.

**$^{13}$C NMR** (151 MHz, $D_2O$) δ[ppm]: 77.09 (d, *J* = 6.7 Hz, 5-position), 75.17 (d, *J* = 33.4 Hz, 6-position), 75.14 (s, 4-position), 74.91 (d, *J* = 32.4 Hz, 2-position), 73.88 (large s, satellite d, J = 39.0 Hz, 1- and 3-position).

**HRMS** *m/z*: $[M – H]^-$ calcd. for $^{13}C_1{}^{12}C_5H_{11}O_6$ 180.0595; found 180.0593.

**4,5[$^{13}C_2$]Ins (S3b)**: yield: 450 mg (starting from 1 g **S1b**, 45 %)

**$^1$H NMR** (600 MHz, $D_2O$) δ[ppm]: 4.19 (t, *J* = 3 Hz, 1H, 2-position), 3.75 (tdd, J = 144.3, 9.9, 4.2 Hz, 1, 4-position) 3.75 (td, *J* = 9.7, 4.7 Hz, 1H, 6-postition), 3.66 (d, *J* = 9.8 Hz, 2H, 1/3-position), 3.40 (tdd, *J* = 140.7, 9.3, 4.1 Hz, 1H, 5-position).

**$^{13}$C NMR** (151 MHz, $D_2O$) δ[ppm]: 77.23 (d, *J* = 38.8 Hz, 5-position), 75.28 (d, *J* = 38.9 Hz, 4+6position), 75.05 (2-position), 74.02 (d, *J* = 6.8 Hz, 1-position), 74.00 (dd, *J* = 39.5, 7.1 Hz, 3-position).

**HRMS** *m/z*: [M – H]$^-$ calcd. for $^{13}C_2{}^{12}C_4H_{11}O_6$ 181.0628; found 181.0627.

**[$^{13}C_6$]Ins (S3c)**: yield: up to 1.55 g (starting from 3 g **S1c**, 52%)

Analytical data for [$^{13}C_6$]Ins were published previously. [2]

#### Synthesis of 1[$^{13}C_1$]InsP$_6$

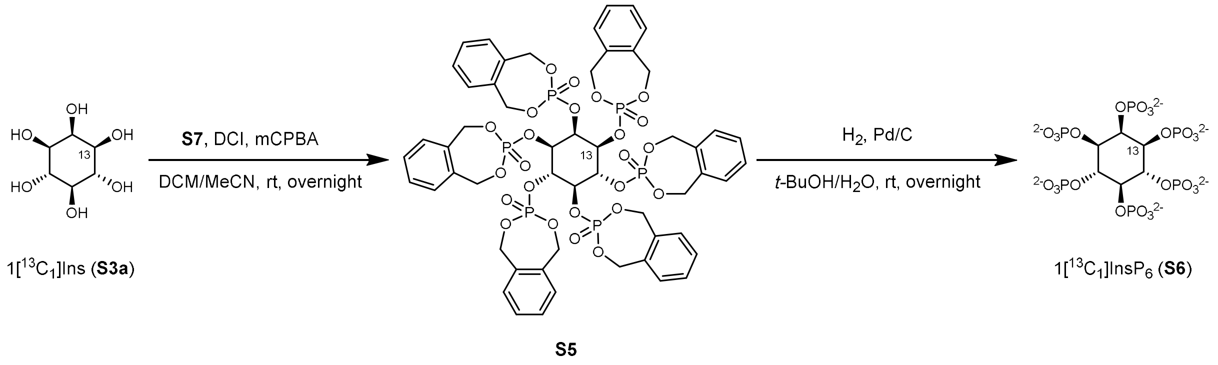

Synthesis of 1[$^{13}C_1$]InsP$_6$ (**S6**) was carried out following to published procedures [2] with slight modifications:

1[$^{13}C_1$]Ins (30 mg, 0.17 mmol) is resuspended together with commercial *o-xylylene N,N-diethylphosphoramidite (**S7**)* (Sigma Aldrich, 360 mg, 1.5 mmol) and a stirring bar in anhydrous acetonitrile under nitrogen atmosphere. To reduce water content further, the suspension is reduced and evaporated under high vacuum for an hour. The dried mixture is then resuspended in 5.5 mL of 1:1 anhydrous dichloromethane:acetonitrile and sonicated briefly. The mixture is cooled to 0 °C using an acetone bath to which dry ice was added in a controlled manner. DCI (254 mg, 2.15 mmol) was added and the gas phase was exchanged three times against nitrogen. The reaction is allowed to warm to room temperature and stirring is continued overnight under nitrogen/argon atmosphere. The subsequent workup is identical as described preciously [2] yielding **S5** in 68% yield (144 mg, 0.116 mmol) with slight impurities.

**S5**:

**$^1$H NMR** (600 MHz, $CDCl_3$) δ[ppm]: 7.40 – 7.31 (m, 20H), 7.27 – 7.25 (m, 4H, overlaps with solvent signal signal), 5.75 (dd, *J* = 13.8, 9.3 Hz, 2H), 5.65 (dt, *J* = 13.0, 8.0 Hz, 3H), 5.59 – 5.51 (m, 6H), 5.39 (dd, *J* = 13.8, 12.3 Hz, 2H), 5.30 – 4.93 (m, 18H).

**$^{13}$C NMR** (151 MHz, $CDCl_3$) δ[ppm:] 138.54, 138.51, 138.35, 138.16, 137.29, 132.34, 132.21, 132.08, 132.07, 132.04, 132.00, 131.83, 131.75, 80.01, 79.80, 79.59, 76.59, 76.56, 76.54, 72.46, 72.40, 72.29, 72.24, 72.19, 72.14, 72.11, 72.06.

**$^{31}$P NMR** (243 MHz, $CDCl_3$) δ[ppm]: -2.81 (d, *J* = 2.9 Hz, 1P), -3.37 (d, *J* = 3.0 Hz, 2P), -4.37 (s, 1P), -4.53 (s, 1P).

**HRMS** *m/z*: [M + H]$^+$ calcd. for $^{13}C_1{}^{12}C_{53}H_{55}O_{24}P_6$ 1274.1537; found 1274.1526.

144 mg (**S5**, 0.116 mmol, 1 eq.) was dissolved in 28 mL *t*-BuOH and Milli-Q® water 6:1, and 250 mg of palladium black (10% Pd/C) was added. The suspension was stirred overnight under hydrogen atmosphere. Upon depletion of starting material (according to LC-MS analysis) 2 ml Milli-Q® water was added to adjust the solvent to a ratio of 4:1 *t*-BuOH:Milli-Q® and stirring under hydrogen atmosphere was continued overnight. The catalyst was removed by centrifuging the suspension in 50 ml centrifugal tubes at 3000 g for 15 min and the supernatant was passed through a PTFE syringe filter (0.45 µm). The catalyst pellet is washed once with 5 ml Milli-Q® water, centrifuged and the supernatant is again filtered and the filtrates are united and tBuOH is removed on the rotavap before the aqueous solution is lyophilized. The resulting white solid is redissolved in 200 ml water and magnesium chloride solution is added to a final concentration of 26 mM (49 eq.). The solution is adjusted with sodium hydroxide solution to a final pH of 9.0 – 9.2 which initiates precipitation of **S6** as a $Mg^{2+}$-complex. The mixture was incubated at 4 °C overnight. The precipitate is pelleted by centrifugation (3000 g, 15 min) in a 50 ml tube and washed twice with 20 ml of 8 mM $MgCl_2$ solution at pH 9.0. The resulting pellet is resuspended in 10 ml of water resulting in a milky solution without any clumps. Meanwhile 15 ml bed volume of Amberlite® IRC-748 (chelating) ion exchange resin (Alfa Aesar, L19570), which was washed in advance extensively with deionized water and methanol and stored in methanol until use) is loaded into a 20 ml peptide reactor column (or another small column) and equilibrated by passing through 100 ml of water. 8 ml bed volume of this Amberlite are added to the $InsP_6$ suspension and incubated at rt on a shaking platform for 30 min until the supernatant turns clear. The content of the tube was transferred onto the remaining Amberlite column and the eluate (gravity-flow) was collected. Additional 20 ml of water pushed through the Amberlite column and the eluates are combined and lyophilized. The resulting clean material was redissolved in $D_2O$ for analysis, filtered through a 0.2 µm PTFE syringe filter and pH was adjusted by addition of DCl solution to 7.0 and dilution to a defined volume. The concentration of 1[$^{13}C_1$]$InsP_6$ was determined against a quantitative NMR-standard (phosphonoacetic acid). In total 0.105 mmol (91%) of clean 1[$^{13}C_6$]$InsP_6$ were obtained.

**1[$^{13}C_6$]$InsP_6$ (S6)**:

**$^1$H NMR** (600 MHz, Deuterium Oxide) δ 4.90 (dq, *J* = 7.7, 2.5 Hz, 1H, 2-position), 4.39 (qt, *J* = 9.5, 2.9 Hz, 2H, 4/6-position), 4.11 (q, *J* = 9.6 Hz, 1H, 3-position), 4.09 (t, *J* = 9.5 Hz, 1H, 5-position), 4.09 (dt, *J* = 144.0, 9.4 Hz, 1H, 1-position).

**$^{13}$C NMR** (151 MHz, $D_2O$) δ[ppm]: 80.16, 78.69, 76.91, 76.24 (1-position).

**$^{31}$P NMR** (243 MHz, $D_2O$) δ 1.94, 1.08, 0.73.

**HRMS** *m/z*: $[M – 2H]^{2-}$ calcd. for $^{13}C_1{}^{12}C_5H_{16}O_{24}P_6$ 329.4251; found 329.4242.

#### Cloning, expression and purification of recombinant human MINPP1

A gene sequence encoding for human MINPP1 (29-487, Uniprot Q9UNW1-1) lacking the N-terminal signal peptide was designed and ordered using Thermo Fisher's GeneArt service. The sequence was codon-optimized for expression in *E. coli* and contains a NdeI (at initial ATG) and XhoI (after the stop codon) restriction site. The MINPP1 gene was cloned into the vector pET-15b using the NdeI and XhoI restriction sites. The resulting plasmid (pET-15b-MINPP1) encodes an N-terminal His-tag with a thrombin cleavage site followed by MINPP1. For plasmid preparation the *E. coli* Top10 strain was used.

The complete nucleotide sequence of the ORF of pET-15b-MINPP1 is as follows:

CC**ATGGGCAGCAGCCATCATCATCATCATCACAGCAGCGGCCTGGTGCCGCGCGGCAGCCATAT
GCGTTGTAGCCTGCTGGAACCGCGTGATCCGGTTGCAAGCAGCCTGAGTCCGTATTTTGGTACAA
AAACCCGTTATGAAGATGTGAATCCGGTTCTGCTGAGCGGTCCGGAAGCACCGTGGCGTGATCCT
GAACTGCTGGAAGGCACCTGTACACCGGTTCAGCTGGTTGCACTGATTCGTCATGGCACCCGTTA
TCCGACCGTTAAACAAATTCGTAAACTGCGTCAGCTGCATGGTCTGCTGCAGGCACGTGGTAGCC
GTGATGGTGGTGCCAGCAGCACCGGTAGTCGTGATCTGGGTGCAGCACTGGCAGATTGGCCTCT**

**GTGGTATGCAGATTGGATGGATGGTCAGCTGGTAGAAAAAGGTCGTCAGGATATGCGTCAACTG**
**GCACTGCGTCTGGCAAGCCTGTTTCCGGCACTGTTTAGCCGTGAAAATTATGGTCGTCTGCGTCT**
**GATTACCAGCAGCAAACATCGTTGTATGGATAGCAGCGCAGCATTTCTGCAAGGTCTGTGGCAGC**
**ATTATCATCCGGGTCTGCCTCCGCCTGATGTTGCAGATATGGAATTTGGTCCGCCTACCGTTAATG**
**ATAAACTGATGCGTTTTTTTGACCATTGCGAGAAGTTTCTGACCGAGGTTGAAAAAAATGCAACCG**
**CACTGTATCATGTGGAAGCATTTAAAACAGGTCCGGAAATGCAGAACATCCTGAAAAAAGTTGCA**
**GCAACCCTGCAGGTTCCGGTTAATGATCTGAATGCCGATCTGATTCAGGTTGCCTTTTTTACCTGT**
**TCATTTGACCTGGCCATTAAAGGTGTTAAAAGCCCGTGGTGTGATGTGTTTGATATTGATGATGCA**
**AAGGTGCTGGAATATCTGAACGATCTGAAACAGTATTGGAAACGCGGTTATGGCTATACCATTAA**
**TAGCCGTAGCAGCTGTACCCTGTTTCAGGATATTTTTCAGCATCTGGATAAAGCCGTTGAACAGAA**
**ACAGCGTAGCCAGCCGATTAGCAGTCCGGTTATTCTGCAGTTTGGTCATGCGGAAACCCTGCTGC**
**CGCTGCTGAGCCTGATGGGTTATTTCAAAGATAAAGAACCGCTGACCGCCTACAACTATAAAAAG**
**CAGATGCATCGTAAATTTCGCAGCGGTCTGATTGTTCCGTATGCAAGCAATCTGATTTTTGTGCTG**
**TATCATTGCGAAAATGCGAAAACCCCGAAAGAACAGTTTCGTGTTCAGATGCTGCTGAATGAAAA**
**AGTTCTGCCGCTGGCATATAGCCAAGAAACCGTTAGCTTTTATGAGGACCTGAAAAACCACTACA**
**AAGATATCCTGCAGAGCTGTCAGACCAGCGAAGAATGTGAACTGGCACGTGCAAATAGCACCAG**
**TGATGAACTGTAACTCGAG**GATCC

Complete ORF of pET-15b-MINPP1. Restriction sites are highlighted (NcoI in yellow, NdeI in green, XhoI in light blue). The sequence encoding MINPP1 is shown in bold and the font colour for chosen component of the protein are changed (His-tag in blue, thrombin cleavage site in orange, catalytic histidine in red).

For protein expression *E. coli* BL21 (DE3) was used which was transformed with the MINPP1-encoding plasmid using the heat-shock method. A 5 ml-overnight culture of the transformed bacterial strain in terrific broth (TB, Formedium) and Ampicillin (100 µg/mL, Roth) at 37 °C was inoculated into 500 ml of TB and Ampicillin. The culture was cooled to 18 °C when $OD_{600nm}$ = 0.5 was reached (~160 min after inoculation). Protein expression was induced at $OD_{600nm}$ = 0.6 (~170 min after inoculation) with 0.6 mM Isopropyl β-D-1-thiogalactopyranoside (IPTG, Thermo Scientific). The culture was incubated at 18 °C for 18-20 h. The bacterial suspension was centrifuged (3000 g, 15 min, 4 °C) upon which a bacterial pellet of ~1.5 g wet weight was obtained. The pellet was resuspended in 50 ml ice-cold lysis buffer (150 mM NaCl, 10 mM Tris*HCl (Roth), pH 8.0, 1 mM DTT (Roth or VWR), 1X cOmplete™ protease inhibitor cocktail (Roche)) and a spatula tip of lysozyme (Roth) and DNAse I (Roche) were added. The bacterial cells were lysed using a homogenizer (LM10 Microfluidizer, Microfluidics, 15000 psi, 5 passages). The resulting suspension was centrifuged (20 000 g, 20 min, 4 °C). The supernatant was used for purification of soluble MINPP1 while the resulting pellet was used for MINPP1 isolation from inclusion bodies (see below). Inclusion body-purification of MINPP1 yielded higher amounts of enzyme and more active MINPP1.

The purification of soluble MINPP1 was adapted from Craxton *et al.* [4]: The supernatant was filtered (VWR vacuum filter, PES 0.45 µm) and the flowthrough was applied to a 5 ml Ni-NTA column (GE, HiTrap IMAC FastFlow) on a FPLC system (NGC Quest 10 Chromatography System, Bio-Rad) equilibrated to buffer A (150 mM NaCl, 10 mM Tris*HCl, pH 8.0, 1 mM DTT). The column was subsequently washed with 5 column volumes (CV) buffer A, 5 CV buffer A:B 10:7, 5 CV buffer B (1 M NaCl, 10 mM Tris*HCl, pH 8.0, 1 mM DTT), 5 CV buffer A with 2% buffer C (buffer C is identical to buffer A containing additional 500 mM imidazole (AppliChem), pH 8.0). For elution a gradient of 2% buffer C in A to 75% buffer C in A over 20 CV was applied. Fractions containing MINPP1 were united, concentrated using centrifugal filters (15 ml 10 kDa MWCO, Amicon Ultra) and dialyzed against 1 L of dialysis buffer 1 (150 mM NaCl, 10 mM Tris*HCl, pH 8.0, 1 mM DTT, 10 Vol-% glycerol (Roth), 0.25% CHAPS (Roth)) twice for 1.5 h. Protein concentration was determined using a BCA assay kit (Pierce™ BCA Protein Assay Kit). Protein solution was aliquoted and stored at -80 °C.

For purification of MINPP1 from inclusion bodies: After removing the lysate, the pellet was washed by thoroughly resuspending in 35 ml ice-cold deionized water, centrifugation (20 000 g, 20 min, 4 °C) and after discarding the supernatant the resulting pellet was washed in the same manner two more times after which a pellet of 1.4 g wet weight was obtained. Per 0.7 g pellet mass, the pellet was resuspended in 30 ml resolubilization buffer (0.2 w/v-% *N*-lauroylsarcosin sodium salt (Sarkosyl, Fisher Scientific), 10 mM Tris*HCl, pH 8.0, 1 mM DTT). The suspension was incubated overnight at 4 °C in a 50 ml-tube under light agitation on a reciprocal shaker. The tube was centrifuged (3000 g, 30 min, 4 °C). 20 ml of recovered supernatant containing MINPP1 was dialyzed first against 1 L dialysis buffer 2 (150 mM NaCl, 10 mM Tris*HCl, pH 8.0, 1 mM DTT, 10 Vol-% glycerol, 0.1 % Triton-X100 (Roth)) for 3 h at 4 °C and then again against fresh dialysis buffer overnight at 4 °C. Protein concentration was determined using a BCA assay kit (Pierce™ BCA Protein Assay Kit). The dialyzed protein solution was adjusted to a final glycerol content of 30 Vol-%, aliquoted, flash-frozen in liquid nitrogen and stored at -80 °C.

The activity of MINPP1 preparations were validated against its substrate 2,3-bisphophoglycerate (BPG) using a Malachite green assay. The activity of inclusion body-purified MINPP1 was determined to be 64 nmol $min^{-1}$ $mg^{-1}$ enzyme, which compares favorably to the value 16 nmol $min^{-1}$ $mg^{-1}$ enzyme reported in the literature.[5]

No decrease in activity was observed after over a year of storage (without freeze-thaw cycles).

**Alternative solubilization buffers**

Different solubilization buffers were also tested for the inclusion body purification of MINPP1 on a smaller scale. Several mild solubilization buffers [6] were unable to sufficiently resolubilize MINPP1 (40 mM TrisHCl, pH 8, with either 5 Vol-% DMSO or 5 Vol-% *n*-propanol (VWR); 90 mM TrisHCl, pH 8.6, 2 M urea). Among the resolubilization buffers only the following managed to solubilize MINPP1: 40 mM TrisHCl, pH 8 with a) 0.2% sarkosyl, b) 8 M urea or c) 6 M guanidinium chloride (see Figure S6).

The resolubilized MINPP1 solution from a) and b) were dialyzed against dialysis buffer with and without Triton-X. In general, it was observed that protein concentration with urea was higher than with Sarkosyl (~1 mg/mL vs. ~3 mg/mL) and with Triton-X-containing dialysis buffer the protein yield was also slightly higher by ~0.2mg/mL. CD spectroscopy suggest that Sarkosyl-based resolubilization increases the amount of folded protein (mainly α-helices as predicted by the Jpred4 tool [7]) as does Triton-X in the dialysis buffer. In comparison, soluble MINPP1 obtained from *e. coli* lysate seems to be even less folded.

For all *in vitro* experiments MINPP1 preparations were used based on resolubilization with sarkosyl and Triton-X-containing dialysis buffer (see above).

#### Enzymatic assays

For the *in vitro* dephosphorylation of $InsP_6$ and $InsP_5$[2OH] by **MINPP1** the following conditions were used unless stated otherwise:

The reaction buffer contained 100 mM NaCl, 100 mM $Na_2SO_4$, 25 mM HEPES, pH* = 7.4, 1 mM DTT, 1 mM EDTA (Sigma), 0.2 mg/mL BSA (Roth), 2 mM CHAPS, 175 µM (or 50 µM) of inositol phosphate substrate, 0.5 µM enzyme. The reactions were carried out in $D_2O$. For each sample (500 µL final volume), the reaction mixture was prepared without InsP substrate in a 1.5 ml microcentrifuge tube, prewarmed to 37 °C for 5 min before the reaction was started by adding the substrate. The reactions were quenched by boiling at 95 °C for 5 min. NMR spectra were recorded without further workup. For the substrate inhibition experiments, InsP5[2OH] was mixed with aliquots of a dilution series of $InsP_6$ before prior to addition to the reaction mixture.

For the dephosphorylation of 2,3-BPG, the same buffer conditions were used but 5 mM 2,3-BPG was used and the reaction was carried out in Milli-Q® water (100 µL total volume per sample) at 37 °C. After 90 min, 20 µL or 5µL of the reaction mixture were transferred into a clear, flat-bottom 96-well plate, diluted with Milli-Q® water to 80 µL final volume and quenched by addition of 20 µL of the Malachite green assay solution (Sigma) and incubated for 30 min at room temperature. Absorption was subsequently measured at 620 nm on a TECANInfinite 200 Pro M-Plex Plate Reader.

#### Cell culture and metabolic labeling

HT29 WT cells were a kind gift from the lab of Jan Carette [8] (William Kaiser laboratory, RRID: CVCL_0320) and HEK293 cell lines (WT and *MINPP1*$^{-/-}$) were a kind gift of the labs of Adolfo Saiardi and Vincent Cantagrel [9], HCT116 were obtained from ATCC. H1975 cells were a kind gift of the Klingmüller lab (originally ATCC, CRL-5908).

Unless stated otherwise all cell lines are cultivated in DMEM (Gibco DMEM high glucose, no glutamine, product no. 11960044) supplemented with streptomycin/penicillin (Gibco), L-glutamine (Gibco Glutamax) and 10% FBS (Pan Biotech), at 37 °C in an atmosphere with 5% $CO_2$, and 95% humidity.

**The metabolic labeling** was conducted as described in a previous publication.[2] Briefly, cells are seeded at a density of $3 \cdot 10^5$ on a 15 cm culture dish in custom DMEM containing no regular inositol nor FBS but 100 µM [$^{13}C_6$]*myo*-inositol (or the respective isotopomer) and 10% dialyzed FBS (Gibco, product no. 26400044) instead (from here on referred to as "labeling medium"). Upon reaching ~85 % confluency, the cells are split into five 15 cm culture dishes in labeling medium. Upon reaching confluency cells were harvested by trypsination, collected in 50 ml tubes and washed twice with 50 ml ice-cold PBS. Packed cell volumes were determined for quantification. The collected cell pellets were either processed immediately after harvest or flash-frozen and stored at -80 °C. Metabolites were extracted by $HClO_4$-extraction. Lyophilized metabolite extracts were redissolved in $D_2O$, re-lyophilized, and finally measured in $D_2O$ (dry $D_2O$ from ampulla, Eurisotop D215T). For quantification 100 µM TMPBr was added to each sample. Standard curves for $InsP_6$ and $InsP_5$[2OH] concentrations against TMPBr are reported previously.[2] The concentrations of other $^{13}C$-labeled InsP species for which no labeled standards were available were estimated using the standard curve for $InsP_6$. Cellular InsP concentrations were backcalculated from PCV.

**$TiO_2$ enrichment of InsPs for NMR samples** was adapted from published procedures.[10] Briefly, 500 µL of InsP containing sample is mixed 1:1 with ice-cold 1 M aq. perchloric acid and incubated for 30 min on ice (frozen samples are thawn directly in the perchloric acid and then incubated on ice). The sample is then centrifuged (10 min, 18 000 g, 4 °C) and the supernatant transferred into a separate 1.5 mL tube containing 5 mg of $TiO_2$ beads (Titanosphere 5 µm, GL Sciences), which were already washed with 500 µL Milli-Q® water and 500 µL 1 M perchloric acid ($HClO_4$, Supelco). The extract and $TiO_2$ beads were mixed on a rotary shaker on low speed for 5 min at 4 °C. The beads were briefly washed twice with 500 µL ice-cold 1 M perchloric acid (note: For centrifugation a table centrifuge (IKA miniG, 6000 rpm) was used at 1 min, and for transferring the supernatant without disturbing the $TiO_2$ beads a 2 µL Eppendorf tip attached to the tip of a 1 mL tip was used). Supernatans were united to check for unbound InsP species, neutralized roughly with 750 µL 2 M potassium hydroxide, centrifuged and the supernatant lyophilized. To eluate InsPs from the $TiO_2$ beads, the beads were incubated with 250 µL of 10% ammonia solution for 5 min at rt on a rotary shaker. After centrifugation the supernatant was collected in a separate tube. The elution step is repeated once more and the eluates are combined. The combined eluates are filtered through a 0.2 µM syringe filter (Sartorius Minisart RC4) which was subsequently rinsed with 150 µL of Milli-Q® water. The filtrate was collected in a new 1.5 mL tube and lyophilized. To reduce the water content for NMR analysis the lyophilized eluates were redissolved in 500 µL $D_2O$ and lyophilized again. For NMR measurement the eluates are redissolved in 500 µL $D_2O$, pH* was adjusted to 6.0.

For the **metabolic flux analysis** *via* CE-MS HEK293 cells were first metabolically labeled with [$^{13}C_6$]Ins as described above over two passages. One week prior to harvest, $4 \cdot 10^5$ of the [$^{13}C_6$]Ins-labeled cells were seeded into one 15 cm dish per time point in [$^{13}C_6$]Ins-labeling medium. For each time point (72, 48, 24, 18, 12.5, 8, 4, 2 and 1 h before harvest) one plate had its medium removed and washed once with 0.9% NaCl solution. The cells were then continued to incubate in 4,5[$^{13}C_2$]Ins-containing labeling medium. For harvesting, the cells of one plate were washed with 25 mL 0.9% NaCl solution, trypsinized (3 mL), then resuspended in 7 mL 0.9% NaCl solution, then pelleted, washed once with 15 mL 0.9% NaCl per pellet and kept on ice until flash-freezing and storage at -80 °C until further processing. For preparing CE-MS each cell pellet was processed as follows: Cells were lysed by resuspending in 1 mL ice-cold 1 M $HClO_4$ (4 °C, 10 min) and centrifuged (18 000 g, 5 min, 4 °C). The supernatant was added to 4 mg of $TiO_2$ beads (prepared as described above) and the $TiO_2$ enrichment protocol was followed as described above until the first lyophilization step. Lyophilized samples were stored at -20 °C until CE-MS measurement. CE-MS measurements were carried out as described above. InsP-isotopomers were quantified relatively to each other.

#### NMR spectra

1[$^{13}C_1$]Ins (**S3a**):

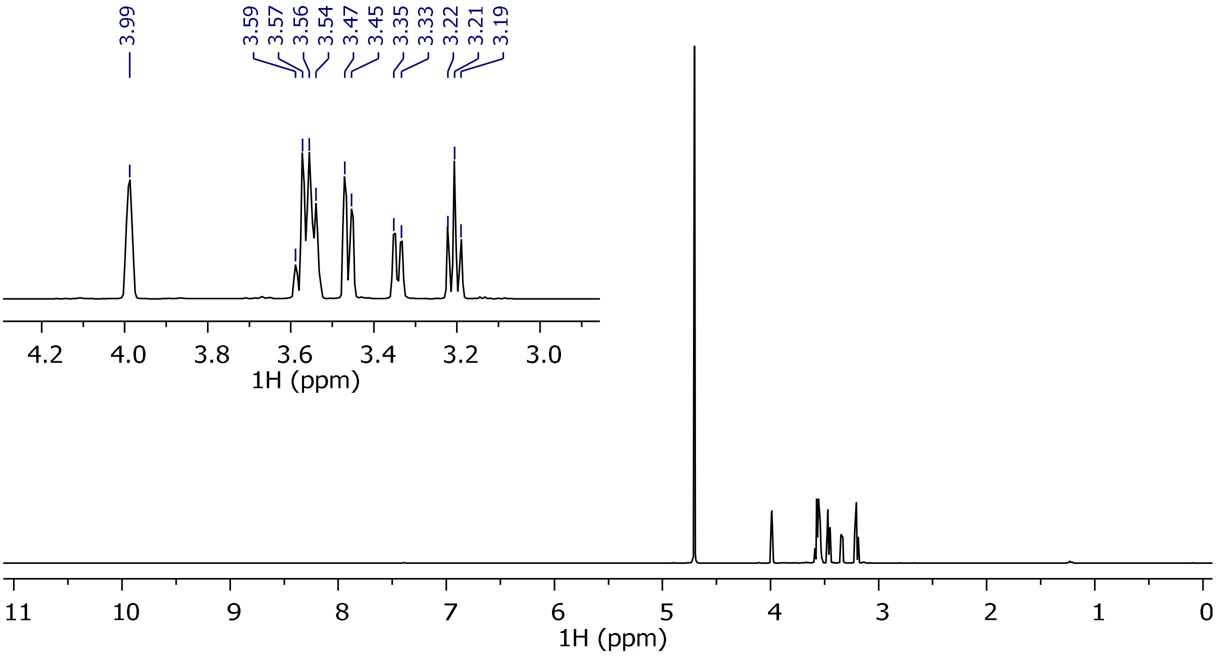

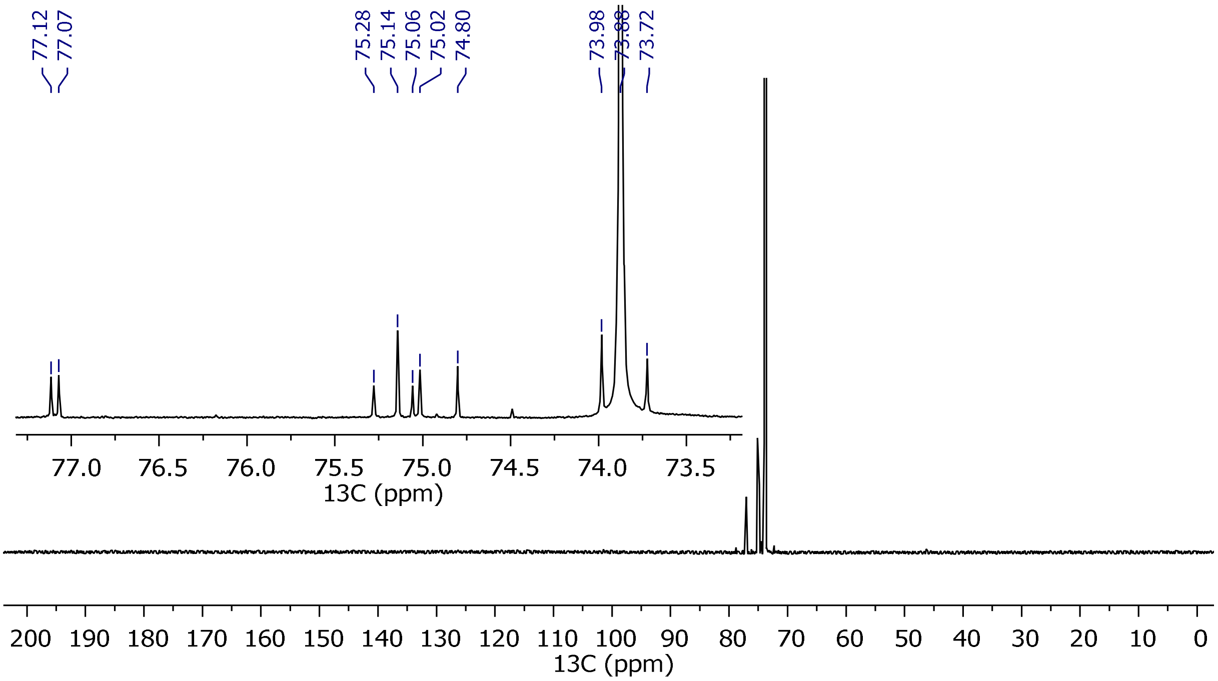

4,5[$^{13}C_2$]Ins (**S3b**):

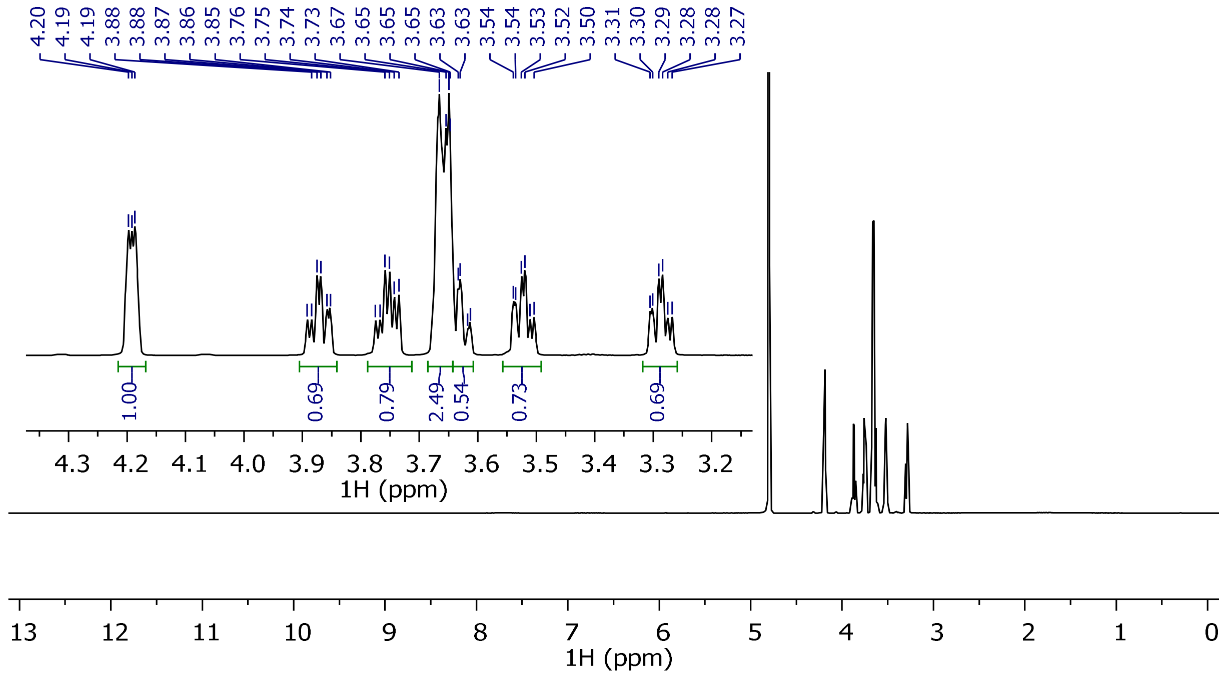

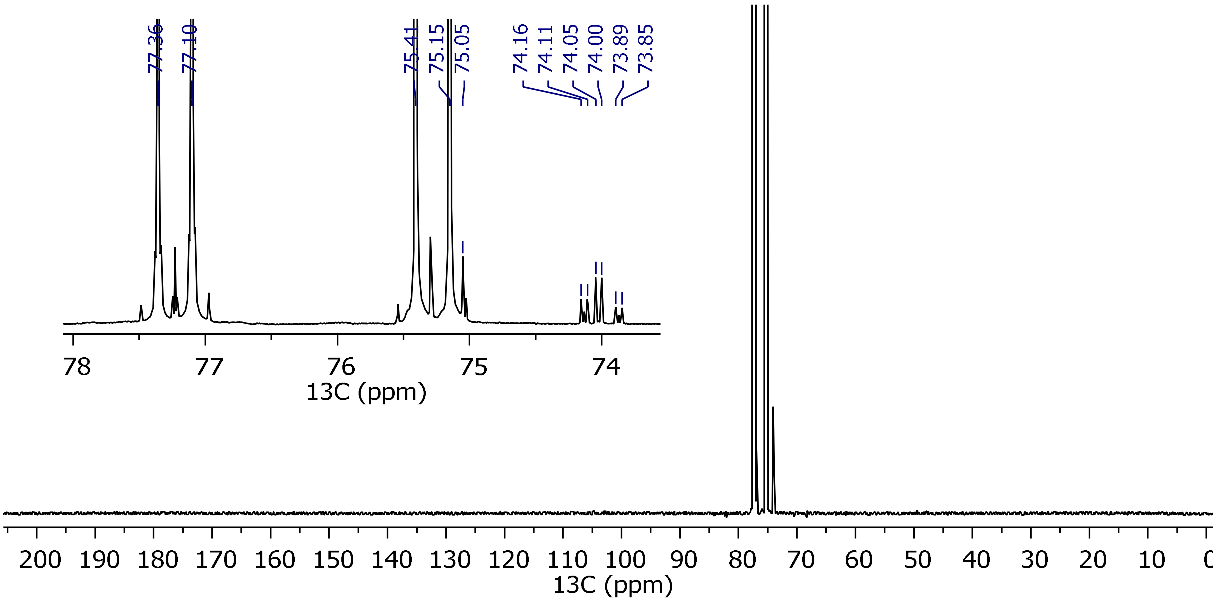

**S5:**

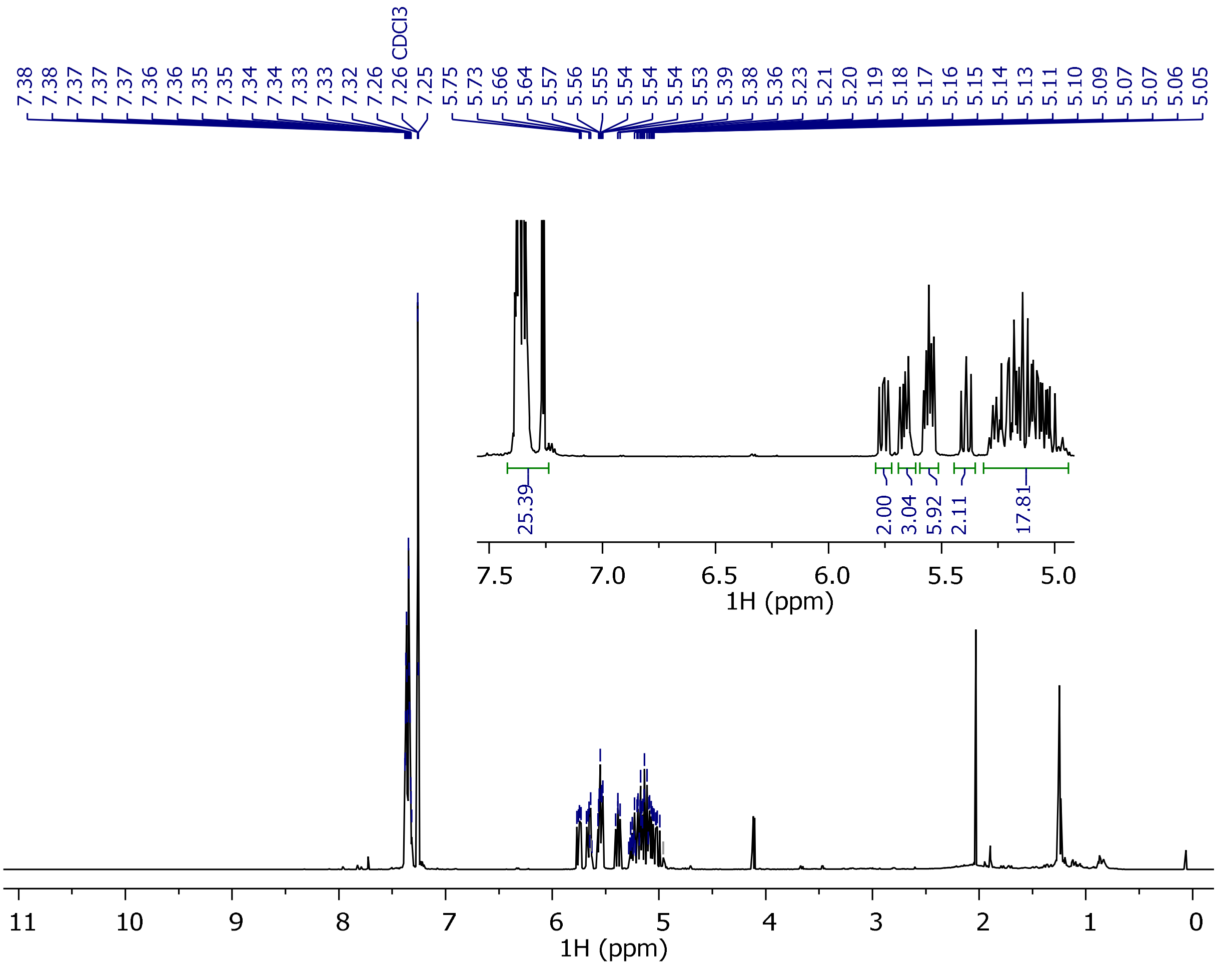

7.38
7.38
7.37
7.37
7.37
7.36
7.36
7.35
7.35
7.34
7.34
7.33
7.33
7.32
7.26
7.26 CDCl3
7.25
5.75
5.73
5.66
5.64
5.57
5.56
5.55
5.54
5.54
5.54
5.53
5.39
5.38
5.36
5.23
5.21
5.20
5.19
5.18
5.17
5.16
5.15
5.14
5.13
5.11
5.10
5.09
5.07
5.07
5.06
5.05
25.39
2.00
3.04
5.92
2.11
17.81
7.5
7.0
6.5
6.0
5.5
5.0
1H (ppm)
11
10
9
8
7
6
5
4
3
2
1
0
1H (ppm)

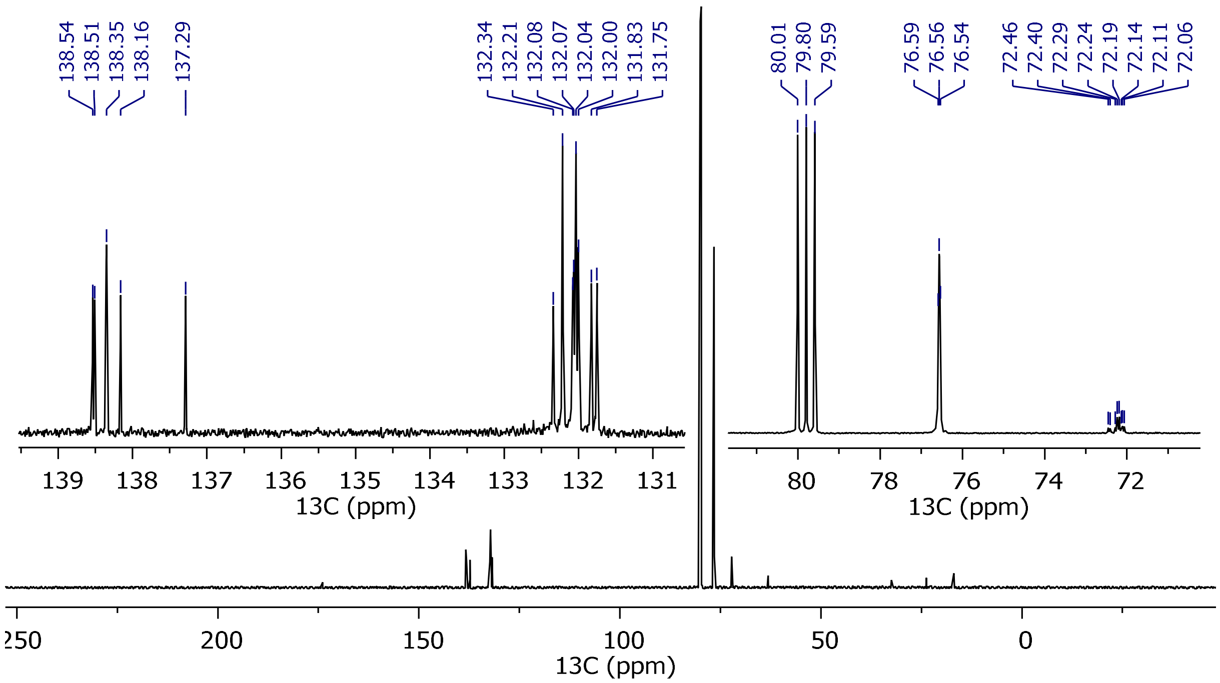
138.54
138.51
138.35
138.16
137.29
132.34
132.21
132.08
132.07
132.04
132.00
131.83
131.75
80.01
79.80
79.59
76.59
76.56
76.54
72.46
72.40
72.29
72.24
72.19
72.14
72.11
72.06
139 138 137 136 135 134 133 132 131
13C (ppm)
80 78 76 74 72
13C (ppm)
250 200 150 100 50 0
13C (ppm)

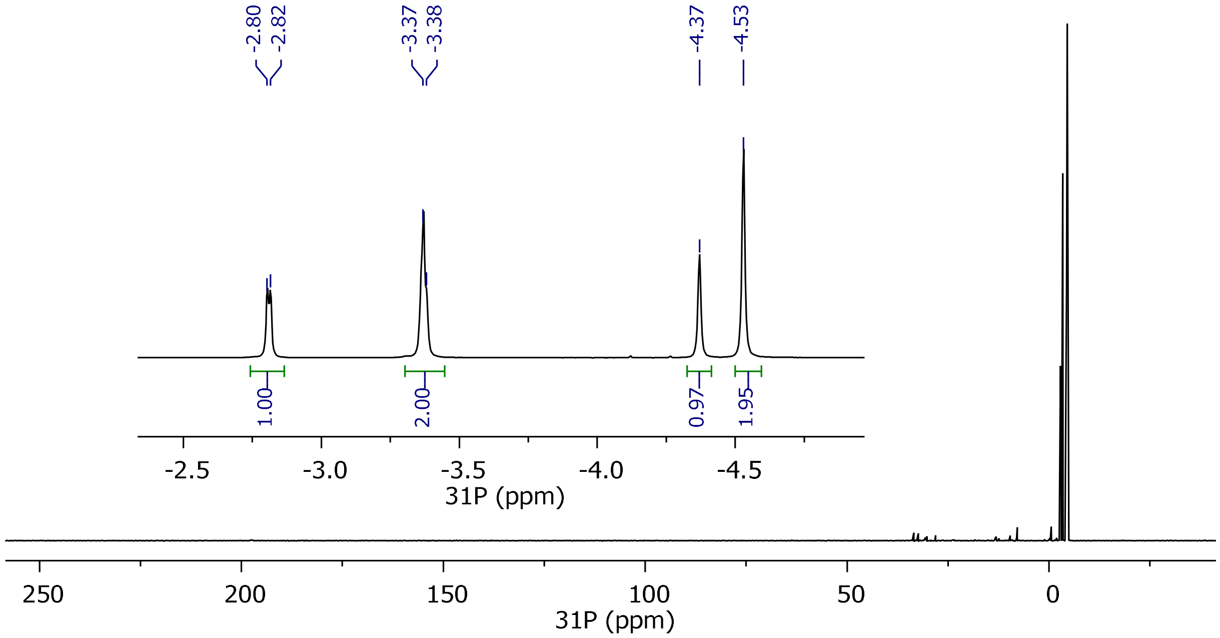
-2.80
-2.82
-3.37
-3.38
-4.37
-4.53
1.00
2.00
0.97
1.95
-2.5 -3.0 -3.5 -4.0 -4.5
31P (ppm)
250 200 150 100 50 0
31P (ppm)

1[$^{13}C_6$]InsP$_6$ (**S6**):

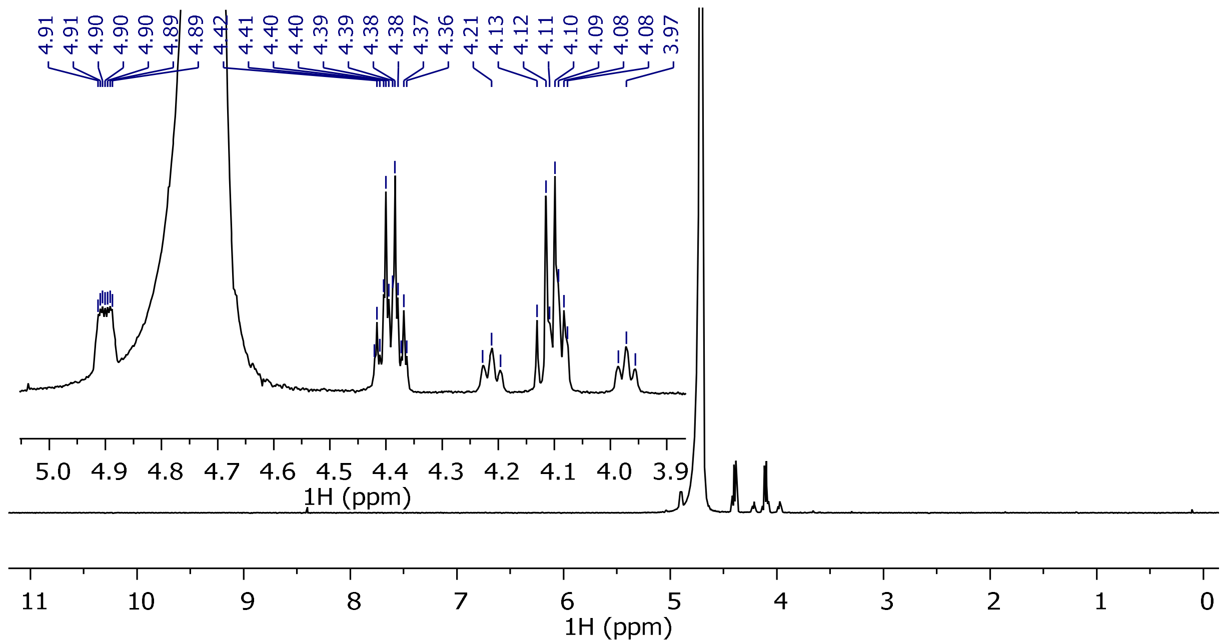

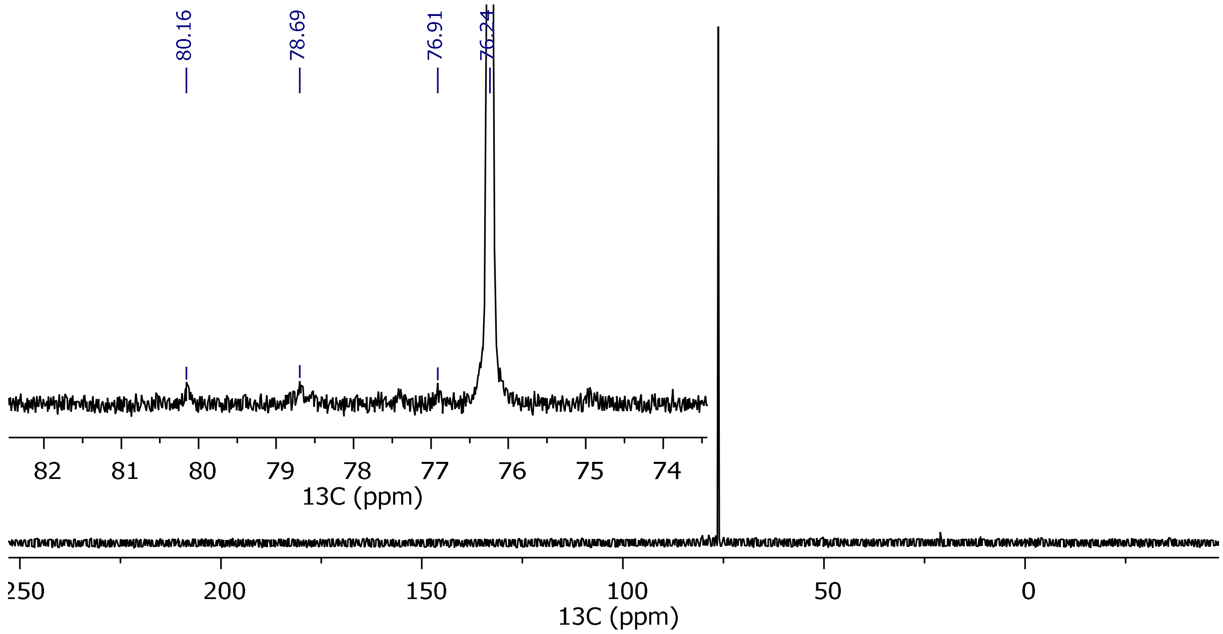

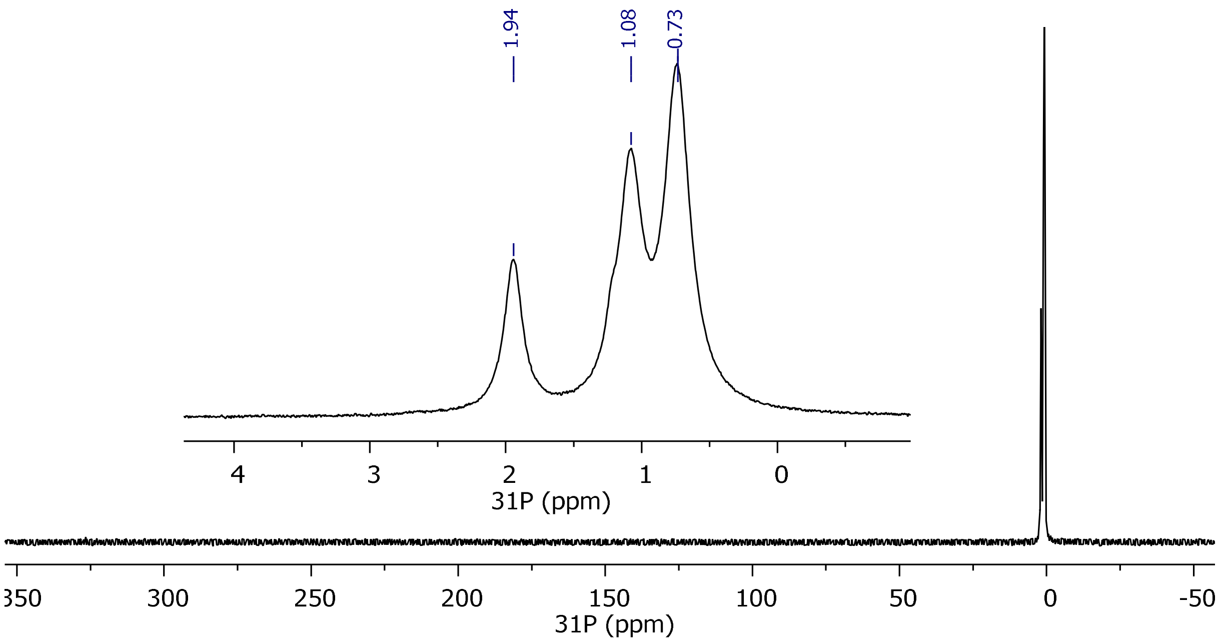
1.94
1.08
0.73
4
3
2
1
0
31P (ppm)
350
300
250
200
150
100
50
0
-50
31P (ppm)
