## Supplementary file SI_Numerical_Analysis_of_MINPP1-mediated_dephosphorylation for "Stable isotopomers of *myo*-inositol to uncover the complex MINPP1-dependent inositol phosphate network"

#### Stable isotopomers of *myo*-inositol to decipher the complex inositol phosphate network and substrate scope of MINPP1

#### List of Abbreviations

| | |
|---|---|
| $InsP_5[2OH]$ | inositol-1,3,4,5,6-pentakisphosphate |
| $InsP_6$ | inositol hexakisphosphate |
| MINPP1 | Multiple Inositol Polyphosphate Phosphatase 1 |
| InsPx | inositol polyphosphate (in general) |
| NMR | nuclear magnetic resonance (spectroscopy) |

### 1 Introduction

This Supplementary Information (SI) contains all information on the numerical evaluation of the reaction rates for the $InsP_5$[2OH]- and $InsP_6$-dephosphorylation from the experimental data. For additional information on the experimental part and figures S1 - S12, please consult the other SI file. Here, we explain the theoretical background of the applied model and the procedure and assumptions that let to the final results presented in main part Fig. 6a and 6b. In this SI, we introduce a six-digit binary representation of the structure names as shown in S13. The numbers in the binary code represent the groups attached to the Cyclohexane scaffold, where the number "0" encodes the hydroxyle group -OH and the number "1" encodes the phosphoryl group $-OPO_3^{2-}$. The position of the number in the binary code (read from left to right) corresponds to the position of the corresponding group in the Cyclohexane scaffold (see S13). For example, the binary representation of Ins(1,2,5,6)$P_4$ reads 110011 and the binary representation of Ins(4,6)$P_2$ is 000101.

| **name representation:** | $InsP_5$[2OH] | $InsP_6$ |
|---|---|---|
| **binary representation:** | 101111 | 111111 |
| **structure:** R - $OPO_3^{2-}$ | 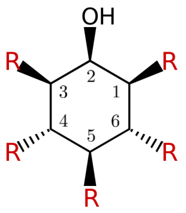  | 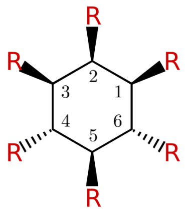  |

S 13:
$InsP_5$[2OH] and $InsP_6$ and their corresponding binary representations and structures.

### 2 Theory

#### 2.1 Full $InsP_5$[2OH] dephosphorylation network

S14 depicts the full reaction pathway of the MINPP1-mediated dephosphorylation of $InsP_5$[2OH]. The network contains all possible intermediates and products including their connection pattern. Each line in S14 represents a rate that describes the reaction from the higher phosphorylated InsPx to the lower phosphorylated InsPx. Since MINPP1 is a phosphatase, the respective reverse reactions are neglected and the network is to be read from top to bottom. The full network contains a total of 75 rates and 31 species, 10 of

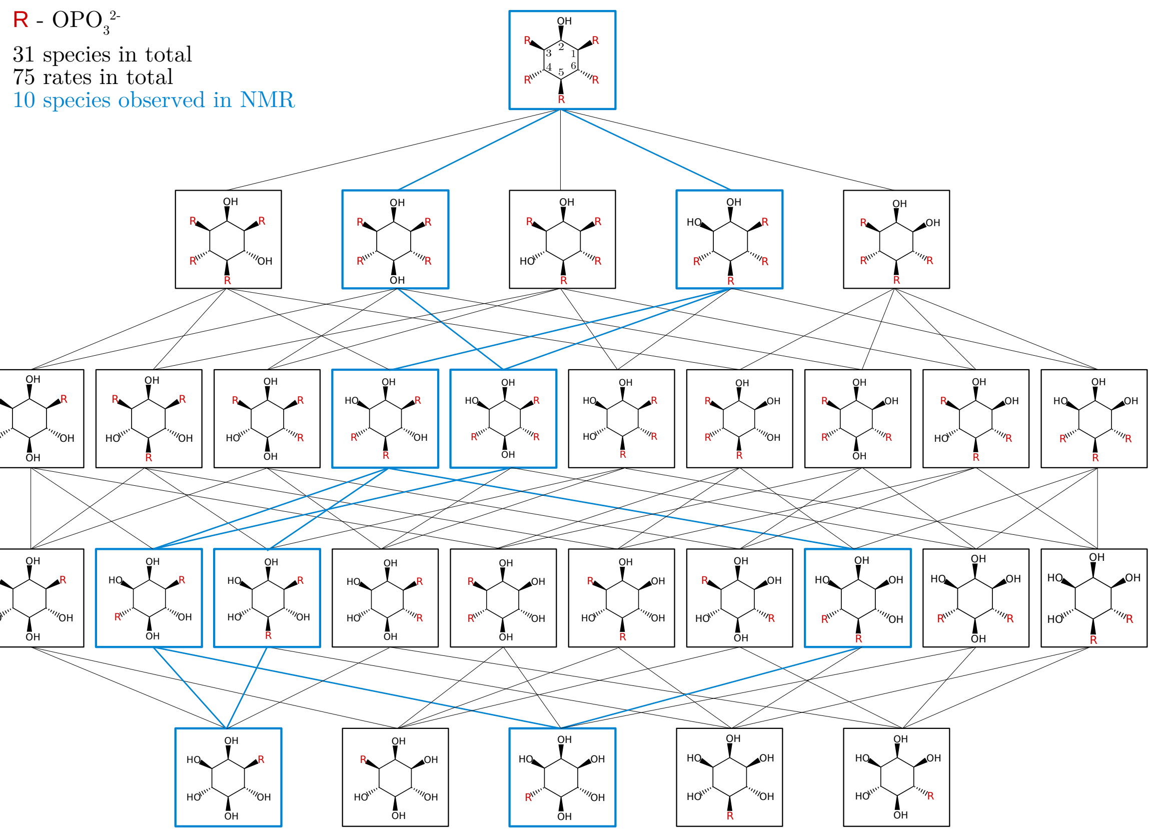

S 14:
Full reaction network with all theoretically possible intermediates and reaction rates of the MINPP1-mediated dephosphorylation of $InsP_5$[2OH]. The structures enclosed in a blue box have been identified in the NMR-experiments with the blue lines depicting the corresponding connection pattern.

which have been identified in the NMR-experiments (blue boxes). We assume that the dephosphorylation network of $InsP_5$[2OH] is dominated by these 10 observed InsPx (see S8) and the corresponding 13 rates (blue lines).

#### 2.2 Full $InsP_6$ dephosphorylation network

S15 depicts the full reaction network of the MINPP1-mediated dephosphorylation of $InsP_6$ in the binary representation. The network contains all possible intermediates and products including their connection pattern. Each line represents a rate that describes the reaction from the higher phosphorylated InsPx to the lower phosphorylated InsPx. Since MINPP1 is a phosphatase, the respective reverse reactions are neglected and the network is to be read from top to bottom. The full network contains a total of 186 rates and 63 species, 14 of which have been identified in the NMR-experiments (blue highlighted boxes) with symmetrically and asymmetrically $^{13}$C-labeled $InsP_6$. We assume that the

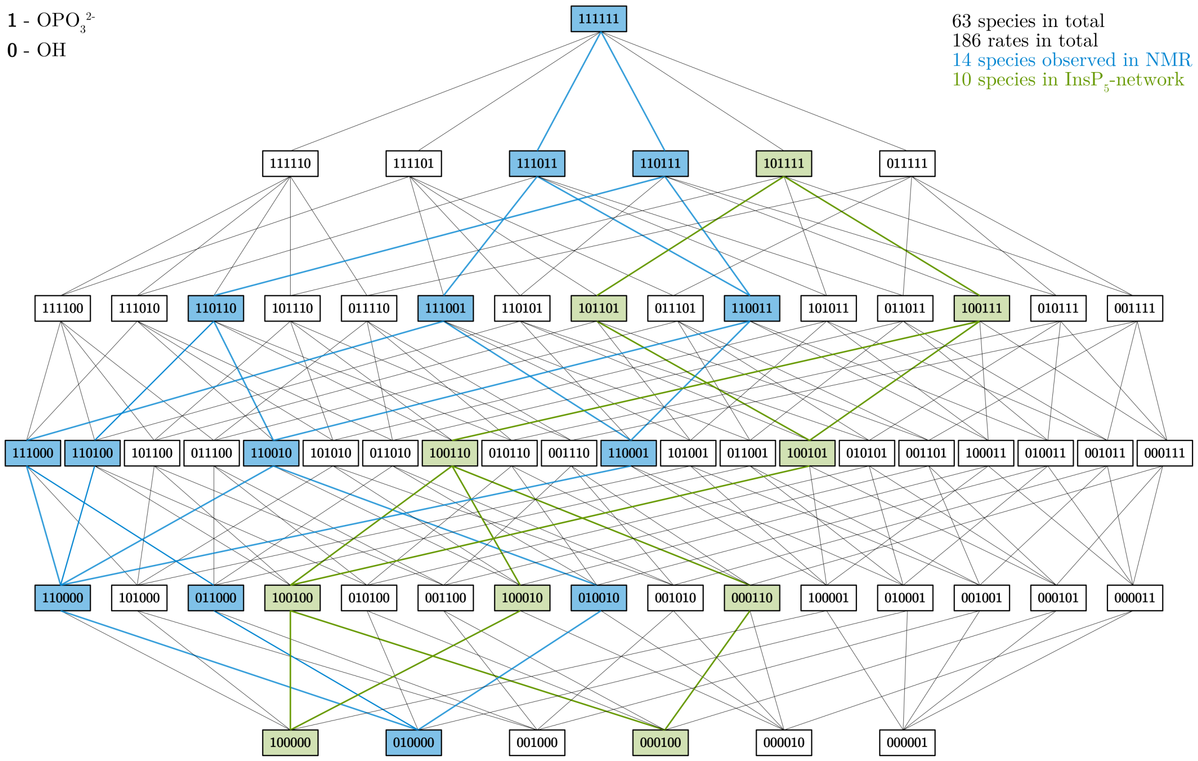

S 15:
Full reaction network with all theoretically possible intermediates and reaction rates of the MINPP1-mediated dephosphorylation of $InsP_6$. The structures highlighted in blue have been identified in the NMR-experiments with the blue lines depicting the corresponding connection pattern. The $InsP_5$ dephosphorylation network (S14) is highlighted in green.

$InsP_6$ dephosphorylation pathway is dominated by the observed 14 InsPx (see S9) and the corresponding 21 rates (blue lines). Additionally, S15 compares the $InsP_6$ dephosphorylation network (highlighted in blue) to the $InsP_5$[2OH] dephosphorylation network (highlighted in green). We can clearly see that the two networks do not overlap and therefore do not share a single structure or rate.

#### 2.3 Master equation formalism

All processes in the $InsP_5$[2OH] dephosphorylation network (SI Fig 14) as well as in the $InsP_6$ dephosphorylation network (S15) are irreversible chemical reactions of the type

$$A_j \xrightarrow{k_{ij}} A_i \quad \text{with} \quad i, j = 0, 1, \ldots, N-1,\ i \neq j\,, \tag{2.1}$$

where species $A_j$ reacts to species $A_i$ with the rate constant $k_{ij}$. $N$ is the total number of species within the network. Please note that we start counting from zero to present the theory in line with the implementation of our analysis in Python3. The rate constants $k_{ij}$ are the matrix elements of the rate matrix $\mathbf{K} \in \mathbb{R}^{N\times N}$. To ensure mass conservation the diagonal elements of $\mathbf{K}$ are defined as the negative of the sum of all other elements in the same column (constraints to master equation)

$$k_{ii} = -\sum_{j=1}^{N} k_{ji} \quad \text{for} \quad i = 0, 1, \ldots, N-1 \text{ and } i \neq j \tag{2.2}$$

such that the sum over each column evaluates to zero. In other words, $\mathbf{K}$ is column-normalized to zero. The vector $\boldsymbol{\phi}(t) \in \mathbb{R}^N$ collects the density (or concentration) $\phi_{A_i}(t)$ at time $t$ of all species. The master equation that corresponds to scheme 2.1 reads

$$\dot{\boldsymbol{\phi}}(t) = \mathbf{K}\,\boldsymbol{\phi}(t)\,, \tag{2.3}$$

where $\dot{\boldsymbol{\phi}}(t)$ is the first derivative of $\boldsymbol{\phi}(t)$ with respect to time. Eq. 2.3 yields $N$ coupled linear homogeneous first order differential equations which describe the kinetics of the entire network. Given the time series $\boldsymbol{\phi}(t)$ (e.g. from experimental data), the master equation can be used to numerically determine the corresponding rates $k_{ij}$ and incorporate additional constraints.[1,2]

##### 2.4 Propagator formalism

In the previous subsection, we describe the kinetics with the corresponding master equation, that is via the change of the density (or concentration) with respect to time. Next, we want to introduce the propagator formalism with which the time evolution of the density (or concentration) for $N$ species can be described directly without the use of time derivatives.
The solution of eq. 2.3 is given as

$$\phi(\tau) = \exp(\mathbf{K}\tau)\phi_0\,, \tag{2.4}$$

where $\phi_0 = \phi(\tau = 0)$ denotes the initial condition at time $\tau = 0$. Eq. 2.4 contains the operator

$$\mathbb{P}(\tau) = \exp(\mathbf{K}\tau) \quad \text{with: } \mathbb{P}(\tau) \in \mathbb{R}^{N\times N}\,, \tag{2.5}$$

which is called propagator and acts on the initial density $\boldsymbol{\phi}(0) = \boldsymbol{\phi}_0 \in \mathbb{R}^N$ to yield the density $\boldsymbol{\phi}_\tau$ after time $\tau$. A given propagator $\mathbb{P}(\tau)$ can only propagate the density in increments of the lag time $\tau$ to yield the time series $\boldsymbol{\phi}_0, \boldsymbol{\phi}_\tau, \boldsymbol{\phi}_{2\tau}, \ldots, \boldsymbol{\phi}_{n\tau}$ with $n \in \mathbb{N}$

according to

$$\begin{aligned}
\boldsymbol{\phi}_{\tau} &= \mathbb{P}(\tau)\,\boldsymbol{\phi}_0 \\
\boldsymbol{\phi}_{2\tau} &= \mathbb{P}(\tau)\,\boldsymbol{\phi}_{\tau} \\
\boldsymbol{\phi}_{3\tau} &= \mathbb{P}(\tau)\,\boldsymbol{\phi}_{2\tau} \\
&\vdots \\
\boldsymbol{\phi}_{n\tau} &= \mathbb{P}(\tau)\,\boldsymbol{\phi}_{(n-1)\tau}
\end{aligned} \tag{2.6}$$

By recursively inserting each equation into the other we get

$$\begin{aligned}
\boldsymbol{\phi}_{n\tau} &= \overbrace{\mathbb{P}(\tau)\mathbb{P}(\tau)\cdots\mathbb{P}(\tau)}^{n \text{ times}}\,\boldsymbol{\phi}_0 \\
\boldsymbol{\phi}_{n\tau} &= \mathbb{P}^n(\tau)\,\boldsymbol{\phi}_0\,.
\end{aligned} \tag{2.7}$$

We want to emphasize that, similar to the master equation formalism, conservation of mass is automatically incorporated into the propagator formalism via the rate matrix $\mathbf{K}$ (see eq. 2.2) such that $\mathbb{P}$ is column-normalized to one. In summary, given a propagator $\mathbb{P}(\tau)$ we can compute the density $\boldsymbol{\phi}_{n\tau}$ at time $n\tau$ either by computing all intermediate steps as described in eqs. 2.6 or evaluate $\boldsymbol{\phi}_{n\tau}$ directly via eq. 2.7. In other words, given all rates $k_{ij}$ we can use the propagator formalism to predict the progress curves of all $N$ species in the network.[2,3]

#### 2.5 Minimization method to numerically determine rates

Let's assume we experimentally obtained the concentration of all $N$ species within a network at different discrete times. In other words, we know the density (concentration) vectors $\boldsymbol{\phi}_0^{\text{exp}}, \boldsymbol{\phi}_{\tau}^{\text{exp}}, \boldsymbol{\phi}_{2\tau}^{\text{exp}}, \ldots, \boldsymbol{\phi}_{n\tau}^{\text{exp}} \in \mathbb{R}^N$ at times $0, \tau, 2\tau, \ldots, n\tau$. From this time series, we can numerically determine the time-derivatives as a finite difference

$$\dot{\boldsymbol{\phi}}_{m\tau}^{\text{exp}} = \frac{\boldsymbol{\phi}_{(m+1)\tau}^{\text{exp}} - \boldsymbol{\phi}_{m\tau}^{\text{exp}}}{\tau} \qquad \text{with} \qquad m = 0, 1, \ldots, n-1\,. \tag{2.8}$$

Additionally, we can define a set of $n$ master equations

$$\dot{\boldsymbol{\phi}}_{m\tau} = \mathbf{K}\,\boldsymbol{\phi}_{m\tau}^{\text{exp}}\,, \tag{2.9}$$

where the elements of $\mathbf{K}$ are unknown. With eq. 2.9 we can predict $\dot{\boldsymbol{\phi}}_{m\tau}$ for a specific choice of $\mathbf{K}$. By selecting one value of $m$ we obtain one master equation for this specific

$m$ as

$$\mathbf{y} = \mathbf{K}\mathbf{x}^{\text{exp}}\,, \tag{2.10}$$

where we abbreviate $\boldsymbol{\phi}_{m\tau}^{\text{exp}} = \mathbf{x}^{\text{exp}} = (x_0^{\text{exp}}, x_1^{\text{exp}}, \ldots, x_{N-1}^{\text{exp}})^T$ and $\dot{\boldsymbol{\phi}}_{m\tau} = \mathbf{y} = (y_0, y_1, \ldots, y_{N-1})^T$. Let $\dot{\boldsymbol{\phi}}_{m\tau}^{\text{exp}} = y^{\text{exp}} = (y_0^{\text{exp}}, y_1^{\text{exp}}, \ldots, y_{N-1}^{\text{exp}})^T$ be the density vector at time $m\tau$ which was calculated numerically from the experimental data via eq. 2.8. We use the mean squared error $\Delta(m\tau)$

$$\Delta(m\tau) = \sum_{i=0}^{N-1} (y_i - y_i^{\text{exp}})^2\,, \tag{2.11}$$

to measure the error of the prediction of $\mathbf{y}$ described in eq. 2.10 and the experimentally obtained $\mathbf{y}^{\text{exp}}$ (eq. 2.8).

$\mathbf{K}$ is column-normalized to zero such that we can substitute the diagonal matrix elements $k_{ii}$ by eq. 2.2. The error in eq. 2.11 then only depends on the off-diagonal elements of $\mathbf{K}$. We can now use a least-square method to minimize $\Delta(m\tau)$ with respect to these off-diagonal elements to get a rate matrix $\mathbf{K}$ that produces $\mathbf{y}$ as close as possible to the experimentally observed $\mathbf{y}^{\text{exp}}$.

With eqs. 2.10 and 2.11 we only made use of the experimental data $\boldsymbol{\phi}_{(m+1)\tau}^{\text{exp}}$ and $\boldsymbol{\phi}_{m\tau}^{\text{exp}}$ at two distinct times $m$ and $m+1$ to determine $\mathbf{K}$. Next, we extend our approach to all values of $m$ such that we can include the entire experimental time series as described in eq. 2.9. In this context, we define the overall error $\Delta$ as a sum over all individual errors $\Delta(m\tau)$ (eq. 2.11)

$$\Delta = \Delta(\tau) + \Delta(2\tau) + \cdots + \Delta(n\tau)\,. \tag{2.12}$$

and minimize eq. 2.12 with respect to the off-diagonal elements $k_{ij}$ in order to get a good estimate for the rate matrix $\mathbf{K}$. Please note, that the described least-square method is particularly effective if the unknown $\mathbf{K}$ is sparse, meaning if it contains a lot of zeros. Furthermore, the method allows for additional constraints (additional to column-normalization, e.g. fixing certain reaction rates to a predefined value) and upper and lower limits for the value of the unknown parameters (e.g. for reaction rates we have $k_{ij} \in [0, 1]$). As indicated in S15, the $InsP_5$[2OH] dephosphorylation is dominated by 10 different InsPx forming a network that includes 13 different rates. Consequently, the corresponding rate matrix $\mathbf{K}$ is $10 \times 10$-dimensional and sparse, which makes the minimization process described above a very well suited tool to determine the reaction rates of the kinetic network. The same argument holds for the $InsP_6$ dephosphorylation network which consists of 12 species and 17 rates, yielding a sparse $12 \times 12$-dimensional rate matrix. Finally, we want to emphasize that a good initial guess for all elements of $\mathbf{K}$ is crucial for the convergence behaviour of a minimization routine as described above.

#### 2.6 Consecutive first-order kinetics

We consider the simplest example of consecutive first order kinetics which is given as

$$A_1 \xrightarrow{k_{21}} A_2 \xrightarrow{k_{32}} A_3\,. \tag{2.13}$$

where one irreversible reaction from species $A_1$ to $A_2$ with the reaction rate $k_{21}$ is followed by a second irreversible reaction from $A_2$ to $A_3$ with the reaction rate $K_{32}$. The naming convention of the reaction scheme follows eq. 2.1. The corresponding master equation is defined in eq. 2.3 with $\boldsymbol{\phi}(t) = (\phi_{A_1}(t), \phi_{A_2}(t), \phi_{A_3}(t))^T \in \mathbb{R}^3$ and $\mathbf{K} \in \mathbb{R}^{3\times3}$. The diagonal elements of $\mathbf{K}$ are given as $k_{11} = -k_{21}$ and $k_{22} = -k_{32}$ (see eq. 2.2). The master equation yields a system of three coupled linear differential equations

$$\dot{\phi}_{A_1}(t) = \frac{\mathrm{d}\phi_{A_1}(t)}{\mathrm{d}t} = -k_{21}\phi_{A_1}(t) \tag{2.14}$$

$$\dot{\phi}_{A_2}(t) = \frac{\mathrm{d}\phi_{A_2}(t)}{\mathrm{d}t} = k_{21}\phi_{A_1}(t) - k_{32}\phi_{A_2}(t) \tag{2.15}$$

$$\dot{\phi}_{A_3}(t) = \frac{\mathrm{d}\phi_{A_3}(t)}{\mathrm{d}t} = k_{32}\phi_{A_2}(t)\,, \tag{2.16}$$

with the analytic solution

$$\phi_{A_1}(t) = \phi^0_{A_1}\, \exp(-k_{21}t) \tag{2.17}$$

$$\phi_{A_2}(t) = k_{21}\phi^0_{A_1}\frac{\exp(-k_{21}t) - \exp(-k_{32}t)}{k_{32} - k_{21}} \tag{2.18}$$

$$\phi_{A_3}(t) = \phi^0_{A_1}\left(1 + \frac{k_{21}\exp(-k_{32}t) - k_{32}\exp(-k_{21}t)}{k_{32} - k_{21}}\right)\,, \tag{2.19}$$

for the case $k_{21} \neq k_{32}$ and the initial condition $\phi_{A_1}(0) = \phi^0_{A_1}$.[4] S16 shows an example for the progress curves defined in eqs. 2.17-2.19 for randomly selected rates.

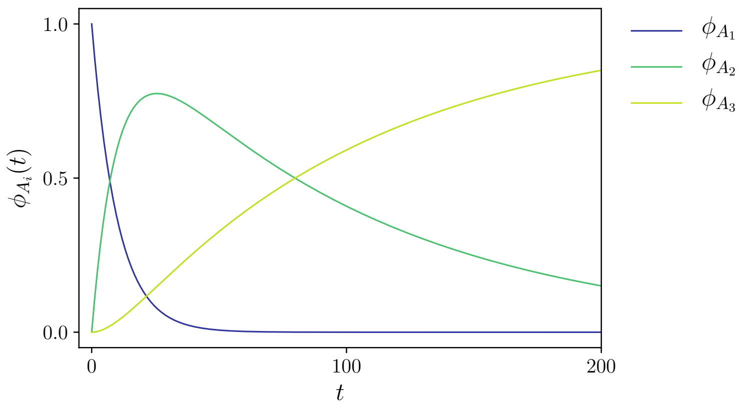

S 16:
The progress curves for two consecutive first order reactions as defined in eq. 2.13 for a set of example rates.

### 3 Analysis

#### 3.1 Analysis protocol

To extract the reaction rates from the scaled experimental data shown in S17 and S17 we use the least-square minimization routine described in SI section 2.5 following the protocol presented below for both replicas of the $InsP_5$[2OH] dephosphorylation respectively as well as for the $InsP_6$ dephosphorylation.

1. scale experimental data such that conservation of mass is fulfilled
2. fit scaled experimental data with analytic fit functions
3. create network assumption
4. use fit functions to generate time-equidistant series $\boldsymbol{\phi}_0^{\text{exp}}, \boldsymbol{\phi}_\tau^{\text{exp}}, \ldots, \boldsymbol{\phi}_{n\tau}^{\text{exp}}$ with resolution $\tau = 1$ min
5. use analytical time-derivatives to compute time series $\dot{\boldsymbol{\phi}}_0^{\text{exp}}, \dot{\boldsymbol{\phi}}_\tau^{\text{exp}}, \ldots, \dot{\boldsymbol{\phi}}_{n\tau}^{\text{exp}}$
6. use corresponding network to set-up rate matrix $\mathbf{K}$ and identify all elements that are not equal zero
7. set-up corresponding master equation and extract set of coupled differential equations
8. determine boundary conditions (bounds) and constraints
9. generate initial guess
10. write numerical program using `scipy.optimize.minimize`
11. compute all rates
12. use the rates to predict corresponding progress curves (eq. 2.7) and compare to scaled experimental data

#### 3.2 Experimental data $InsP_5$[2OH] dephosphorylation

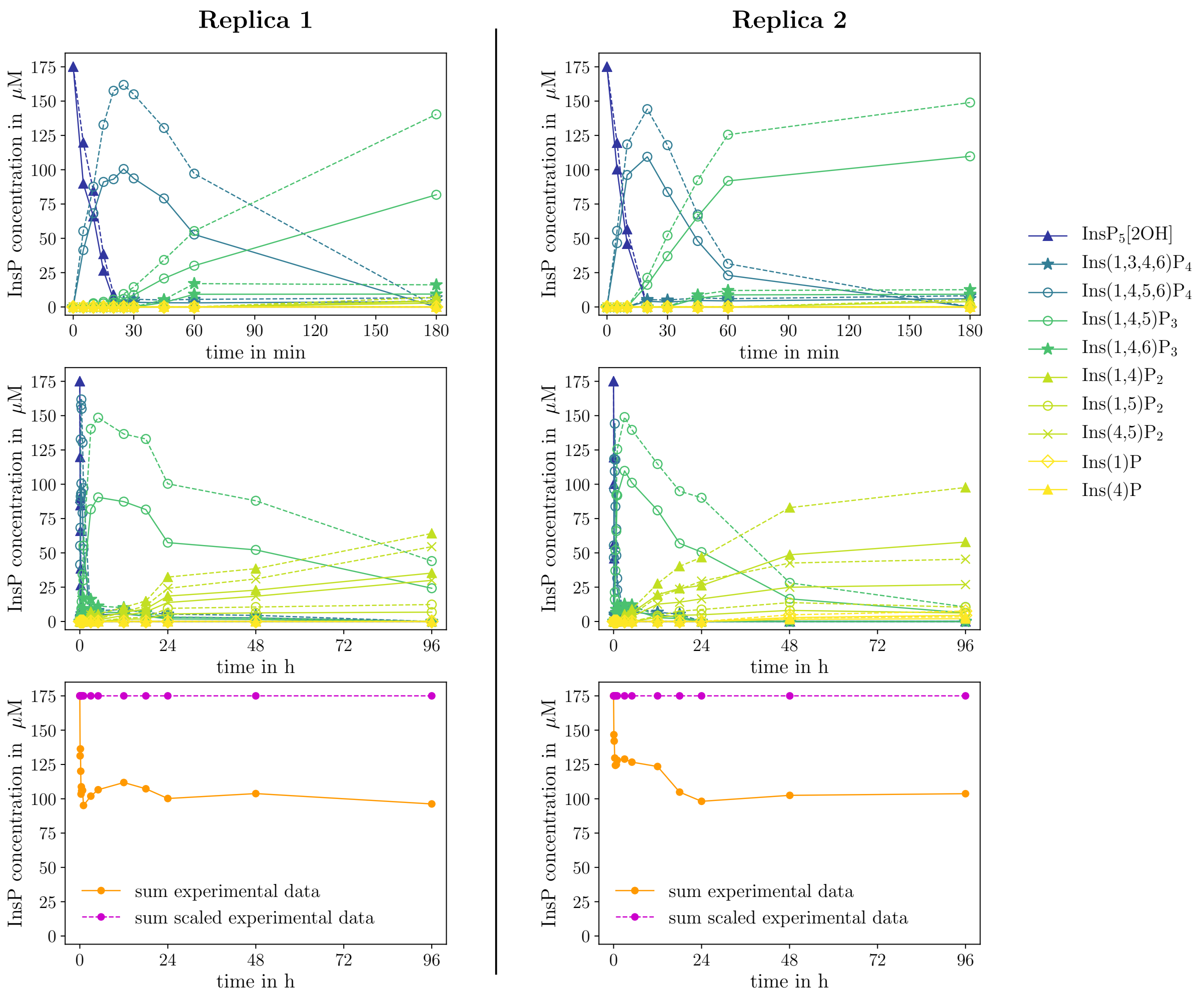

S 17:
Measured progress curves (solid lines) and scaled progress curves (dashed lines) of MINPP1 reaction with 175 $\mu$M [$^{13}C_6$]$InsP_5$[2OH] as a concentration time series for two replicas with identical experimental setup. The top row magnifies the first 180 min, the middle row shows the full 96 hours and the bottom row represents the sum over all progress curves, for each of the experiments respectively. The dashed lines in the left column top and middle represent the same data set as main part Fig. 5a.

S17 shows the progress curves of two replicas of the MINPP1 reaction with 175 $\mu$M [$^{13}C_6$]$InsP_5$[2OH] (columns) with identical experimental setup, where we conducted NMR-measurements at 10 different points in time for replica 1 and at 8 different points in time for replica 2. The plot at the top magnifies the first 180 min of the experiment and the plot in the middle shows the full 96 hours time interval of the measurements, respectively. The solid lines represent the experimentally measured concentration time series $\phi_i^{\mathrm{exp}}(t)$ with $i = 0, \ldots, N-1$ of the $N = 10$ species that could be identified in the NMR-experiments. The orange line in the bottom plot represents the corresponding sum $S^{\mathrm{exp}}(t)$ over the concentrations of all 10 species at each point in time

$$S^{\text{exp}}(t) = \sum_{i=0}^{N-1} \phi_i^{\text{exp}}(t)\,, \tag{3.1}$$

We can clearly see that $S^{\text{exp}}(t) \neq 175\ \mu\text{M}$ for all $t$, meaning that we "loose" mass during the course of the experiment and conservation of mass is not fulfilled by the original experimental data. Since conservation of mass is crucial for the kinetic model we use to extract rates from the experimental data, we correct for the loss of mass by scaling the experimental data according to

$$\phi_i^{\text{scaled}}(t) = \frac{\phi_i^{\text{original}}(t)}{S^{\text{exp}}(t)} \cdot 175\ \mu\text{M} \tag{3.2}$$

such that

$$S^{\text{scaled}}(t) = \sum_{i=0}^{N-1} \phi_i^{\text{scaled}}(t) = 175\ \mu\text{M} \qquad \forall t\,. \tag{3.3}$$

The scaled progress curves $\phi_i^{\text{scaled}}(t)$ are shown as dashed lines in S17 and in main part Fig. 5a. The results of of both replicas exhibit similar behaviour but the progress curves of replica 2 indicate slightly faster kinetics. In the main part we chose replica 1 as representative for both replicas. To extract kinetic of the $\text{InsP}_5$ dephosphorylation, we perform the numerical analysis on both replicas separately and compare the resulting rates in SI section 4.1. Please note that we solely use the scaled progress curves for the numerical analysis.

To prepare the scaled experimental data for the numerical analysis, we fitted the progress curves of each species with an analytic fit function. The fit functions provide access to more and time-equidistant data points and analytical derivatives for each progress curve (no numerical derivatives necessary). S18 compares the fit function to the scaled experimental data for both replicas and SI table 1 summarizes the fit functions and the corresponding fit parameters. Please note, that we used the kinetic function defined in eq. 2.18 to fit the progress curve of $\text{Ins}(1,4,5)\text{P}_3$ (dark green circles) meaning that the fit parameters $k_1$ and $k_2$ can already be interpreted as reaction rates.

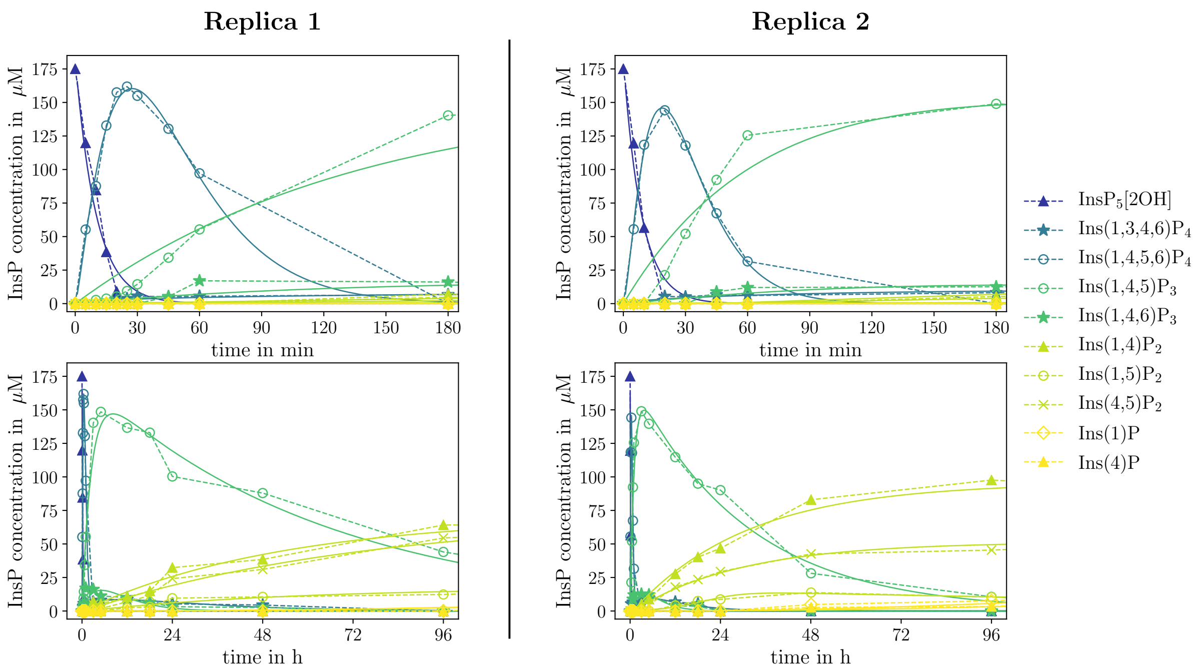

S 18:
Scaled progress curves (dashed lines) of MINPP1 reaction with 175 $\mu$M [$^{13}C_6$]$InsP_5$[2OH] as a concentration time series and corresponding fit functions (solid lines). The top row magnifies the first 180 min and the bottom row shows the full 96 hours.

SI Table 1:
Fit functions and parameters used to fit the scaled experimental data of $[^{13}C_6]InsP_5[2OH]$ dephosphorylation for two replicas.

| **species** | **Replica 1** | **Replica 2** |
|---|---|---|
| $InsP_5[2OH]$ | $f(t) = a\exp(-kt)$<br>$a = 182.834$<br>$k = 0.100$ | $f(t) = a\exp(-kt)$<br>$a = 182.020$<br>$k = 0.114$ |
| $Ins(1,3,4,6)P_4$ | $f(t) = a \cdot t^b \cdot \exp(-kt)$<br>$a = 0.734$<br>$b = 0.459$<br>$k = 0.0008$ | $f(t) = a \cdot t^b \cdot \exp(-kt)$<br>$a = 0.575$<br>$b = 0.612$<br>$k = 0.002$ |
| $Ins(1,4,5,6)P_4$ | $f(t) = a \cdot t^b \cdot \exp(-kt)$<br>$a = 7.508$<br>$b = 1.322$<br>$k = 0.048$ | $f(t) = a \cdot t^b \cdot \exp(-kt)$<br>$a = 7.722$<br>$b = 1.520$<br>$k = 0.080$ |
| $Ins(1,4,5)P_3$ | $f(t) = \frac{c_0\, k_1}{k_2 - k_1}(\exp(-k_1 t) - \exp(-k_2 t))$<br>$k_1 = 0.006$<br>$k_2 = 0.000262$<br>$c_0 = 167.378$ | $f(t) = \frac{c_0\, k_1}{k_2 - k_1}(\exp(-k_1 t) - \exp(-k_2 t))$<br>$k_1 = 0.0153$<br>$k_2 = 0.000553$<br>$c_0 = 169.494$ |
| $Ins(1,4,6)P_3$ | $f(t) = a \cdot t^b \cdot \exp(-kt)$<br>$a = 0.181$<br>$b = 0.937$<br>$k = 0.003$ | $f(t) = a \cdot t^b \cdot \exp(-kt)$<br>$a = 0.111$<br>$b = 1.115$<br>$k = 0.005$ |
| $Ins(1,4)P_2$ | $f(t) = S - (S - a) \cdot \exp(-bt)$<br>$a = -0.052$<br>$b = 0.0003$<br>$S = 70.446$ | $f(t) = S - (S - a) \cdot \exp(-bt)$<br>$a = -2.263$<br>$b = 0.0005$<br>$S = 95.617$ |

| species | new experiment | old experiment |
| --- | --- | --- |
| $\mathrm{Ins(1,5)P_2}$ | $f(t) = S-(S-a)\cdot\exp(-bt)$<br>$a = -0.289$<br>$b = 0.0003$<br>$S = 17.523$ | $f(t) = \frac{d}{a+b\exp(-ct)}\exp(-gt)$<br>$a = 2.029$<br>$b = 78.948$<br>$c = 0.003$<br>$d = 33.623$<br>$g = 0.00007$ |
| $\mathrm{Ins(4,5)P_2}$ | $f(t) = S-(S-a)\cdot\exp(-bt)$<br>$a = -0.582$<br>$b = 0.0002$<br>$S = 71.352$ | $f(t) = S-(S-a)\cdot\exp(-bt)$<br>$a = -1.303$<br>$b = 0.0005$<br>$S = 51.389$ |
| $\mathrm{Ins(1)P_1}$ | $f(t) = a\,t^2$<br>$a = 7.89\cdot 10^{-8}$ | $f(t) = a\,t^2$<br>$a = 1.7\cdot 10^{-7}$ |
| $\mathrm{Ins(4)P_1}$ | $f(t) = a\,t^2$<br>$a = 7.06\cdot 10^{-8}$ | $f(t) = a\,t^2$<br>$a = 9.64\cdot 10^{-8}$ |

#### 3.3 Experimental data $InsP_6$ dephosphorylation

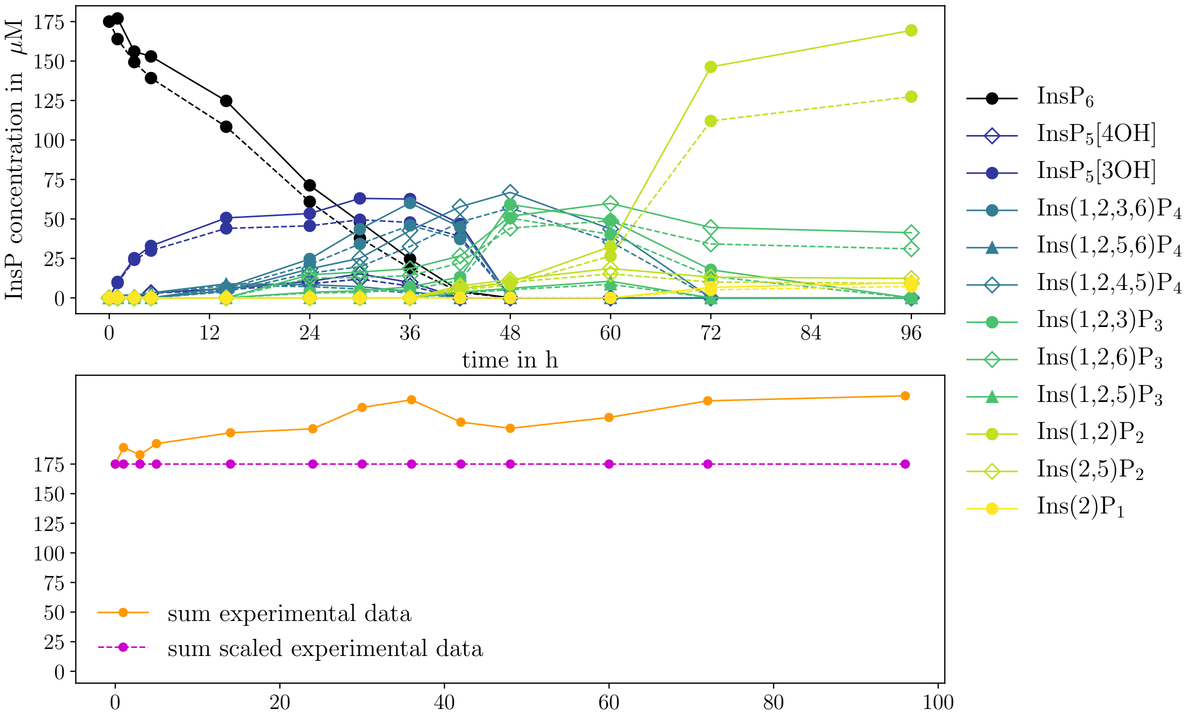

S 19:
Measured progress curves (solid lines) and scaled progress curves (dashed lines) of MINPP1 reaction with 175 $\mu$M [$^{13}C_6$]$InsP_6$ as a concentration time series (top) and the sum over all progress curves at each point in time (bottom). The dashed lines in the top plot represent the same data set as main part Fig. 5c.

S9 and S15 depict our assumption of the complete MINPP1-mediated $InsP_6$ dephosphorylation pathway and main part Fig. 5d shows the corresponding simplified version. The pathway contains the enantiomers Ins(1,2,4)$P_3$ and Ins(1,2,6)$P_3$ and the enantiomers Ins(1,2)$P_2$ and Ins(2,3)$P_2$ which can only be distinguished in asymmertrically $^{13}$C-labeled NMR experiments. However, we base our numerical analysis on the progress curves shown in S19,top which resulted from the MINPP1 reaction with symmetrically labeled [$^{13}C_6$]$InsP_6$. Consequently, both pairs of enantiomers are represented by one progress curve each, which we labeled with one representative for each pair of enantiomers. This reduces the network in S15 from 14 to 12 different species. The solid lines in S19,top, represent the experimentally measured concentration time series $\phi_i^{\text{exp}}(t)$ with $i = 0, \dots, N-1$ and $N = 12$. The orange line in S19, bottom represents the sum $S^{\text{exp}}(t)$ over the concentrations of all 12 species at each point in time. Since the original experimental data does not obey conservation of mass over the entire time axis, we scale the data according to eq. 3.2. The scaled progress curves are shown as dashed lines and are equivalent to the solid lines in main part Fig. 5c. We want to emphasize that we solely use the scaled progress curves for all further analysis. To prepare the scaled experimental data for the numerical analysis, we fitted the progress curves of each species with an analytic fit function. S20 compares the fit function to the scaled experimental data and SI table 2 summarizes the fit functions and the corresponding fit parameters. We used eq. 2.17 as fit function to fit the $InsP_6$ progress curve which means that we can interpret the fit parameter $k$ as the reaction rate that quantifies the depletion of $InsP_6$ over time. Moreover, we used eq. 2.18 to fit the $InsP_5$[3OH] progress curve and thus we can interpret the fit parameters $k_1$ and $k_2$ as reaction rates that dictate the growth and the depletion of $InsP_5$[3OH] concentration in time.

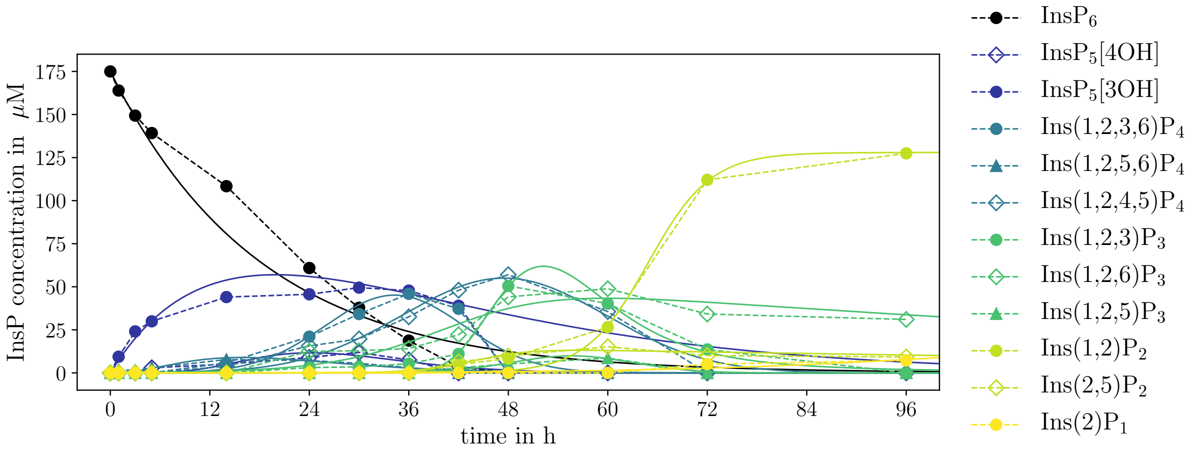

S 20:
Scaled progress curves (dashed lines) of MINPP1 reaction with 175 $\mu$M [$^{13}C_6$]$InsP_6$as a concentration time series and corresponding fit functions (solid lines).

SI Table 2:
Fit functions and parameters used to fit the scaled experimental of $[^{13}C_6]InP_6$ dephosphorylation.

| **species** | **fit data** |
|---|---|
| $InsP_6$ | $f(t) = a\exp(-kt)$<br>$a = 175.00$<br>$k = 0.000932$ |
| $InsP_5[4OH]$ | $f(t) = at^b\exp(-kt)$<br>$a = 6.2 \cdot 10^{-18}$<br>$b = 6.697$<br>$k = 0.00460$ |
| $InsP_5[3OH]$ | $f(t) = \frac{k_1\,c_0}{k_2 - k_1}(\exp(-k_1\,t) - \exp(-k_2\,t))$<br>$k_1 = 0.00074$<br>$k_2 = 0.00093$<br>$c_0 = 175$ |
| $Ins(1,2,3,6)P_4$ | $f(t) = a \cdot \exp\left(-\frac{(t-\mu)^2}{\sigma^2}\right)$<br>$a = 45.149$<br>$\mu = 2057.099$<br>$\sigma = 709.875$ |
| $Ins(1,2,5,6)P_4$ | $f(t) = at^b\exp(-kt)$<br>$a = 0.0000021$<br>$b = 2.603$<br>$k = 0.00272$ |
| $Ins(1,2,4,5)P_4$ | $f(t) = a \cdot \exp\left(-\frac{(t-\mu)^2}{\sigma^2}\right)$<br>$a = 55.045$<br>$\mu = 2849.46$<br>$\sigma = 1045.39$ |

| species | fit data |
|---|---|
| Ins(1,2,3)$P_3$ | $f(t) = a\ \exp\left[-b\Big(1-\exp(-c(t-d))\Big)^2\right]$<br>$a = 61.89$<br>$b = 4.578$<br>$c = 0.00076$<br>$d = 3133.40$ |
| Ins(1,2,6)$P_3$ | $f(t) = \dfrac{d}{a+b\ \exp(-c\,t)}\ \exp(-e\,t)$<br>$a = 0.0973$<br>$b = 45.655$<br>$c = 0.00249$<br>$d = 7.480$<br>$e = 0.00014$ |
| Ins(1,2,5)$P_3$ | $f(t) = a\cdot\exp\left(-\dfrac{(t-\mu)^2}{\sigma^2}\right)$<br>$a = 9.909$<br>$\mu = 3338.044$<br>$\sigma = 626.472$ |
| Ins(1,2)$P_2$ | $f(t) = \dfrac{d}{a+b\ \exp(-c\,t)}$<br>$a = 0.000046$<br>$b = 1394.199$<br>$c = 0.00442$<br>$d = 0.00590$ |
| Ins(2,5)$P_2$ | $f(t) = \dfrac{d}{a+b\ \exp(-c\,t)}\ \exp(-e\,t)$<br>$a = 11.970$<br>$b = 0.000024$<br>$c = -0.00497$<br>$d = 0.000468$<br>$e = -0.00486$ |
| Ins(2)$P_1$ | $f(t) = \dfrac{d}{a+b\ \exp(-c\,t)}$<br>$a = -8.741$<br>$b = -3609.11$<br>$c = 0.000978$<br>$d = -168.34$ |

#### 3.4 Analysis setup $InsP_5$[2OH] dephosphorylation

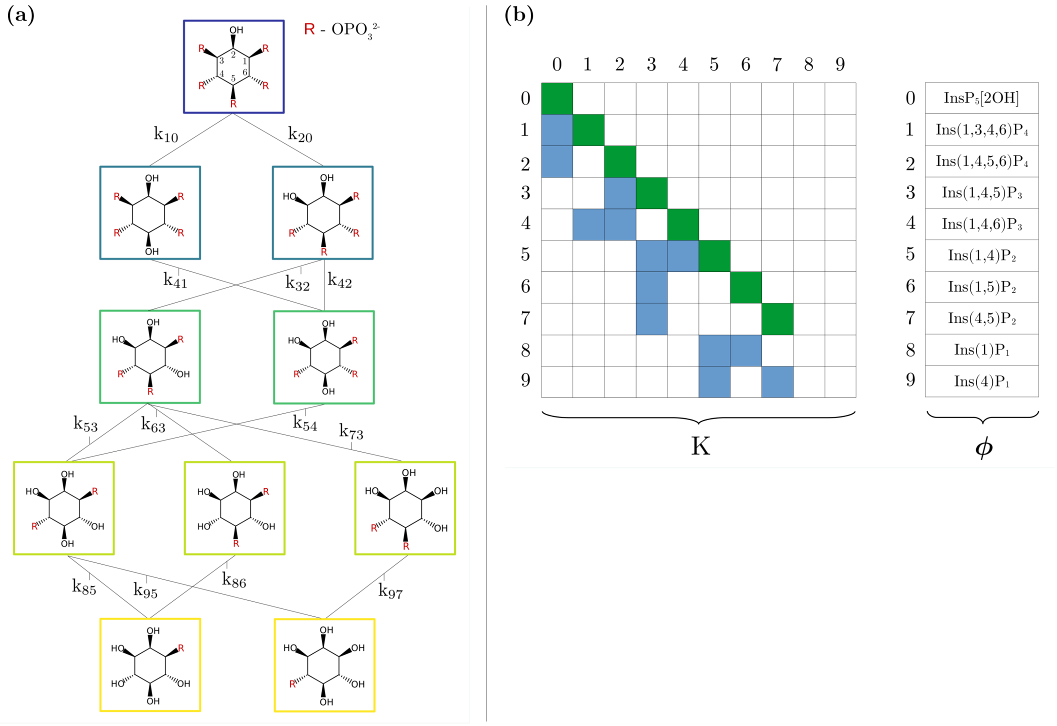

S 21:
(a) Assumed network for MINPP1 reaction with 175 $\mu$M $InsP_5$[2OH] including all reactions rates. A copy of this network is shown in S8 and a simplified version is depicted in main part Fig. 5b.
(b) Schematic representation of the corresponding rate matrix and density (concentration) vector. Matrix: The white squares mark all matrix elements that are equal to zero, the blue squares all elements that are not zero and the green squares represent the diagonal elements defined via eq. 2.2. Vector: The representation indicates which vector element is associated with which InsP.

**Network**:
Based on the NMR-data (see main part Fig. 3 and S17), we assume that the $InsP_5$[2OH] dephosphorylation network is dominated by 10 different InsPx that form the network depicted in S21, a. All possible reactions from a higher phosphorylated InsPx to a lower phosphorylated InsPx are indicated with a line and are associated with a reaction rate $k_{ij} \neq 0$.

**Density (concentration) vector and corresponding time derivative**:
We use the fit functions (SI table 1) to create time-equidistant data points $\boldsymbol{\phi}_0^{\text{exp}}, \boldsymbol{\phi}_\tau^{\text{exp}}, \ldots, \boldsymbol{\phi}_{n\tau}^{\text{exp}}$ and $\dot{\boldsymbol{\phi}}_0^{\text{exp}}, \dot{\boldsymbol{\phi}}_\tau^{\text{exp}}, \ldots, \dot{\boldsymbol{\phi}}_{n\tau}^{\text{exp}}$ (eq. 2.9) with a resolution of $\tau = 1$ min for each replica.

**Rate matrix:**
To build the rate matrix $\mathbf{K}$, we number all species in the network from zero to nine in a left-to-right and top-to-bottom fashion and assign the corresponding rates according to eq. 2.1. These rates are represented in S21, b as blue squares. The diagonal elements

(green squares) are defined via eq. 2.2 and given as

$$
\begin{aligned}
k_{00} &= -(k_{10} + k_{20}) \\
k_{11} &= -k_{41} \\
k_{22} &= -(k_{32} + k_{42}) \\
k_{33} &= -(k_{53} + k_{63} + k_{73}) \\
k_{44} &= -k_{54} \\
k_{55} &= -(k_{85} + k_{95}) \\
k_{66} &= -k_{86} \\
k_{77} &= -k_{97}\,.
\end{aligned} \tag{3.4}
$$

All other matrix elements are equal to zero (white squares in S21, b).

**Set of differential equations:**
With the rate matrix **K** defined, we can now formulate the corresponding master equation (eq. 2.3) which yields the following set of 10 coupled first-order differential equations

$$
\begin{array}{lcllllllll}
\dot{\phi}_0 & = & k_{00}\phi_0 & & & & & & & \\
\dot{\phi}_1 & = & k_{10}\phi_0 & +k_{11}\phi_1 & & & & & & \\
\dot{\phi}_2 & = & k_{20}\phi_0 & & +k_{22}\phi_2 & & & & & \\
\dot{\phi}_3 & = & & & +k_{32}\phi_2 & +k_{33}\phi_3 & & & & \\
\dot{\phi}_4 & = & & +k_{41}\phi_1 & +k_{42}\phi_2 & & +k_{44}\phi_4 & & & \\
\dot{\phi}_5 & = & & & & +k_{53}\phi_3 & +k_{54}\phi_4 & +k_{55}\phi_5 & & \\
\dot{\phi}_6 & = & & & & +k_{63}\phi_3 & & & +k_{66}\phi_6 & \\
\dot{\phi}_7 & = & & & & +k_{73}\phi_3 & & & & +k_{77}\phi_7 \\
\dot{\phi}_8 & = & & & & & & +k_{85}\phi_5 & +k_{86}\phi_6 & \\
\dot{\phi}_9 & = & & & & & & +k_{95}\phi_5 & & +k_{97}\phi_7
\end{array} \tag{3.5}
$$

**Constraints and bounds:**
Here, we report the applied constraints that yielded the best results for the reaction rates reported in main part Fig. 6a and 6b. In total, we constrained 4 rates ($k_{10}, k_{41}, k_{73}$ and $k_{42}$), which leaves us with 9 reaction rates that have to be optimized during the minimization routine.

$k_{10}$ and $k_{41}$:
Since the progress curve of Ins(1,3,4,6)$P_4$ evolves at very low concentrations (less than 9 $\mu$M for the entire time series), we decided to exclude this species from the analysis and

set the corresponding rates to zero, $k_{10}, k_{41} = 0$, for both replicas.

$\underline{k_{73}}$:
As mentioned in section 3.2 , we use eq. 2.18 as fit function for the progress curve of Ins(1,3,4)$P_3$ (SI table 1). Since this function emerges from a kinetic model, we can interpret the corresponding fit parameters $k_1$ and $k_2$ as kinetic rates with $k_1$ describing the increase and $k_2$ the decrease of concentration. The increase in Ins(1,3,4)$P_3$ concentration is determined by $k_{32}$ and the decrease is determined by $k_{53} + k_{63} + k_{73}$. We use the fit parameter $k_2$ ($2.62 \cdot 10^{-4}$ min$^{-1}$ for replica 1 and $5.53 \cdot 10^{-4}$ min$^{-1}$ for replica 2) to constrain the rate $k_{73}$ as $k_{73} = k_2 - k_{53} - k_{63}$ and leave $k_{32}$ unconstraint for each replica respectively.

$\underline{k_{42}}$:
During our analysis, we found that the increase in Ins(1,4,6)Ins$P_3$ concentration was generally overestimated by the minimization routine and thus we decided to constrain the rate $k_{42}$ towards Ins(1,4,6)Ins$P_3$ by hand. In an iterative procedure, we found that the constraint $k_{42} = 0.001$ yields the most promising results for both replicas.

bounds:
Since reaction rates are a real number between zero and one, we bound all rates to the interval $k_{ij} \in [10^{-6}, 1]$.

**Initial guess:**
We built the rate matrix **K** by formulating an initial guess for each rate, used eq. 2.7 to predict the corresponding progress curves and compared these prediction to the scaled experimental data (S17, dashed lines). In an iterative procedure, we corrected the rates by hand until the set of rates produced progress curves that roughly matched the scaled experimental progress curves. The set of rates is summarized in SI table 3 and serves as initial guess for our minimzation routine.

**Technical details:**
We used Python3 and `scipy.optimize.minimize`[5] to implement the minimization routine described in SI section 2.5, where we passed eq. 2.12 as objective function to be minimized, the initial guess and bounds as described above and left all other parameters at their default settings. Since we constraints 4 of the 13 rates, the implemented minimization routine optimizes the remaining 9 rates such that the resulting rate matrix **K** yields progress curves that are in excellent agreement with the scaled experiment data.

SI Table 3:
Initial guess for all rates of the $InsP_5$[2OH] dephosphorylation network in $min^{-1}$ for both replicas.
*) The marked rates are subject to constraints.

| **rate** | **replica 1** | **replica 2** |
|---|---|---|
| $k_{10}$ | $0.00^*$ | $0.00^*$ |
| $k_{20}$ | $9.76 \cdot 10^{-2}$ | $9.76 \cdot 10^{-2}$ |
| $k_{41}$ | $0.00^*$ | $0.00^*$ |
| $k_{32}$ | $6.84 \cdot 10^{-3}$ | $6.84 \cdot 10^{-3}$ |
| $k_{42}$ | $1.00 \cdot 10^{-3*}$ | $1.00 \cdot 10^{-3*}$ |
| $k_{53}$ | $1.09 \cdot 10^{-4}$ | $1.09 \cdot 10^{-4}$ |
| $k_{63}$ | $4.16 \cdot 10^{-5}$ | $4.16 \cdot 10^{-5}$ |
| $k_{73}$ | $(2.62 \cdot 10^{-4} - k_{53} - k_{63})^*$ | $(5.53 \cdot 10^{-4} - k_{53} - k_{63})^*$ |
| $k_{54}$ | $6.11 \cdot 10^{-4}$ | $6.11 \cdot 10^{-4}$ |
| $k_{85}$ | $1.00 \cdot 10^{-5}$ | $1.00 \cdot 10^{-5}$ |
| $k_{95}$ | $1.08 \cdot 10^{-5}$ | $1.08 \cdot 10^{-5}$ |
| $k_{86}$ | $5.77 \cdot 10^{-5}$ | $5.77 \cdot 10^{-5}$ |
| $k_{97}$ | $2.40 \cdot 10^{-6}$ | $1.00 \cdot 10^{-6}$ |

#### 3.5 Analysis setup $InsP_6$ dephosphorylation

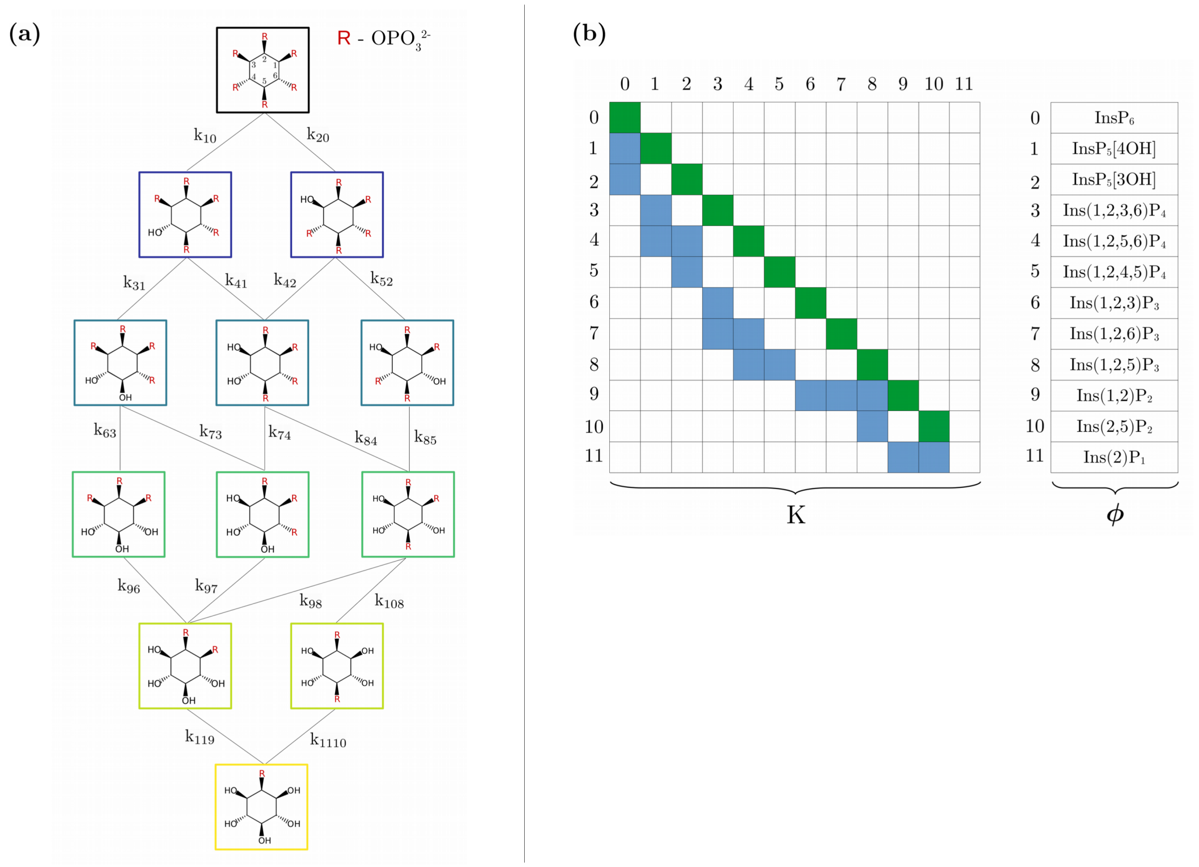

S 22:
(a) Assumed network for MINPP1 reaction with 175 $\mu$M $InsP_6$ including all reactions rates.
(b) Schematic representation of the corresponding rate matrix and density (concentration) vector. Matrix: The white squares mark all matrix elements that are equal to zero, the blue squares all elements that are not zero and the green squares represent the diagonal elements defined via eq. 2.2. Vector: The representation indicates which vector element is associated with which InsP.

##### Network:

S22a depicts the network assumption on which we base our numerical analysis. Ins(1,2,6)$P_3$ and Ins(1,2)$P_2$ are chosen as representatives for their respective pair of enantiomiers (see also section 3.3). All possible reactions from a higher phosphorylated InsPx to a lower phosphorylated InsPx are indicated with a line and are associated with a reaction rate $k_{ij} \neq 0$. The network consits of 12 different InsPx and 17 reaction rates.

##### Density (concentration) vector and corresponding time derivative:

We use the fit functions (SI table 2) to create time-equidistant data points $\boldsymbol{\phi}_0^{\text{exp}}, \boldsymbol{\phi}_\tau^{\text{exp}}, \ldots, \boldsymbol{\phi}_{n\tau}^{\text{exp}}$ and $\dot{\boldsymbol{\phi}}_0^{\text{exp}}, \dot{\boldsymbol{\phi}}_\tau^{\text{exp}}, \ldots, \dot{\boldsymbol{\phi}}_{n\tau}^{\text{exp}}$ (eq. 2.9) with a resolution of $\tau = 1$ min.

**Rate matrix:**
To build the rate matrix $\mathbf{K}$, we number all InsPx included in the network from zero to eleven in a left-to-right and top-to-bottom fashion and assign the corresponding rates according to eq. 2.1. S22,b represents these rates as blue squares. The diagonal elements (green squares) are defined via eq. 2.2 and given as

$$\begin{aligned}
k_{00} &= -(k_{10} + k_{20}) \\
k_{11} &= -(k_{31} + k_{41}) \\
k_{22} &= -(k_{42} + k_{52}) \\
k_{33} &= -(k_{63} + k_{73}) \\
k_{44} &= -(k_{74} + k_{84}) \\
k_{55} &= -k_{85} \\
k_{66} &= -k_{96} \\
k_{77} &= -k_{97} \\
k_{88} &= -(k_{98} + k_{108}) \\
k_{99} &= -k_{119} \\
k_{1010} &= -k_{1110} \, .
\end{aligned} \tag{3.6}$$

All other matrix elements are equal to zero (white squares).

**Set of differential equations:**
With the rate matrix $\mathbf{K}$ we can formulate the corresponding master equation (eq. 2.3) which yields the following set of coupled first-order differential equations

$$\begin{aligned}
y_0 &= +k_{00}x_0 \\
y_1 &= +k_{10}x_0 + k_{11}x_1 \\
y_2 &= +k_{20}x_0 + k_{22}x_2 \\
y_3 &= +k_{31}x_1 + k_{33}x_3 \\
y_4 &= +k_{41}x_1 + k_{42}x_2 + k_{44}x_4 \\
y_5 &= +k_{52}x_2 + k_{55}x_5 \\
y_6 &= +k_{63}x_3 + k_{66}x_6 \\
y_7 &= +k_{73}x_3 + k_{74}x_4 + k_{77}x_7 \\
y_8 &= +k_{84}x_4 + k_{85}x_5 + k_{88}x_8 \\
y_9 &= +k_{96}x_6 + k_{97}x_7 + k_{98}x_8 + k_{99}x_9 \\
y_{10} &= +k_{108}x_8 + k_{1010}x_{10} \\
y_{11} &= +k_{119}x_9 + k_{1110}x_{10}
\end{aligned} \tag{3.7}$$

**Constraints and bounds:**
Here, we report the applied constraints that yielded the best results for the reaction rates reported in SI table 5. In total, we constrain 3 reaction rates ($k_{10}, k_{20}$ and $k_{52}$) which left us with 14 reaction rates that have to be optimized during the minimization routine.

$k_{20}$ and $k_{52}$:
As mentioned in section 3.3, we used eq. 2.18 as fit function for the progress curve of $InsP_5$[3OH] (SI table 2). Consequently, we can interpret the fit parameter $k_1$ as reaction rate that describes build-up of concentration and $k_2$ as reaction rate that describes the decrease of concentration. According to the network in S22a, the increase of $InsP_5$[3OH] concentration is solely determined by $k_{20}$ and thus we constrain $k_{20} = k_1 = 7.4 \cdot 10^{-4}$ $\text{min}^{-1}$. The decrease of $InsP_5$[3OH] concentration is determined by $k_{42} + k_{52}$ and we set the constraint $k_{52} = k_2 - k_{42} = 9.3 \cdot 10^{-4} - k_{42}$.

$k_{10}$:
We used eq. 2.17 as fit function for the progress curve of $InsP_6$ (SI table 2) and thus the fit parameter $k$ represents the reaction rate that dictates the decrease of $InsP_6$ concentration over time. According to the network in S22,a, this decrease is described by $k_{10} + k_{20}$ and we constrain $k_{10} = k - k_{20} = 1.9 \cdot 10^{-4}$ $\text{min}^{-1}$.

bounds: We set the bounds $k_{108} \in [10^{-3}, 1]$, $k_{119} \in [10^{-5}, 1]$ and $k_{1110} \in [10^{-4}, 1]$ to brute-force increase the influence of these rates on the network and prevent the minimization routine from setting all of them to the lowest possible value $10^{-6}$. All other rates were bound to the interval $k_{ij} \in [10^{-6}, 1]$.

**Initial guess:**
To generate a good initial guess for the unconstrained rates, we started with a reduced network that included $InsP_6$, $InsP_5$[4OH], $InsP_5$[3OH], Ins(1,2,3,6)$P_4$, Ins(1,2,5,6)$P_4$ and Ins(1,2,4,5)$P_4$ and performed a minimization run. Next, we increased the network by including Ins(1,2,3)$P_3$, Ins(1,2,6)$P_3$ and Ins(1,2,5)$P_3$ and repeated the minimization, where we used the results from the previous run for $k_{10}, k_{31}$ and, $k_{41}$ and the default initial guess for the remaining rates. Finally, we repeated this step with the full network which yielded a good initial guess for all 13 unconstrained rates as shown in SI table 4.

**Technical details:**
For all technical details, the reader is referred to section 3.4.

SI Table 4:
Initial guess (in $min^{-1}$) and bounds for all rates in the [$^{13}C_6$]Ins$P_6$ dephosphorylation network.
*) The marked rates are subject to constraints.

| rate | initial guess | bounds |
|---|---|---|
| $k_{10}$ | $1.9 \cdot 10^{-4*}$ | - |
| $k_{20}$ | $7.4 \cdot 10^{-4*}$ | - |
| $k_{31}$ | $2.9 \cdot 10^{-3}$ | $[10^{-6}, 1]$ |
| $k_{41}$ | $1.0 \cdot 10^{-5}$ | $[10^{-6}, 1]$ |
| $k_{42}$ | $3.0 \cdot 10^{-4}$ | $[10^{-6}, 1]$ |
| $k_{52}$ | $9.3 \cdot 10^{-4*} - k_{42}$ | - |
| $k_{63}$ | $5.0 \cdot 10^{-4}$ | $[10^{-6}, 1]$ |
| $k_{73}$ | $3.7 \cdot 10^{-4}$ | $[10^{-6}, 1]$ |
| $k_{74}$ | $1.4 \cdot 10^{-3}$ | $[10^{-6}, 1]$ |
| $k_{84}$ | $1.5 \cdot 10^{-4}$ | $[10^{-6}, 1]$ |
| $k_{85}$ | $4.2 \cdot 10^{-4}$ | $[10^{-6}, 1]$ |
| $k_{96}$ | $2.0 \cdot 10^{-4}$ | $[10^{-6}, 1]$ |
| $k_{97}$ | $2.0 \cdot 10^{-4}$ | $[10^{-6}, 1]$ |
| $k_{98}$ | $3.9 \cdot 10^{-3}$ | $[10^{-6}, 1]$ |
| $k_{108}$ | $1.0 \cdot 10^{-5}$ | $[10^{-3}, 1]$ |
| $k_{119}$ | $1.0 \cdot 10^{-3}$ | $[10^{-5}, 1]$ |
| $k_{1110}$ | $1.0 \cdot 10^{-3}$ | $[10^{-4}, 1]$ |

### 4 Results

#### 4.1 Results Ins$P_5$[2OH] dephosphorylation

The numerically determined set of rates for both replicas are presented in S24. We can see that replica 2 exhibits slightly faster kinetics than replica although both share an identical experimental setup. Excluding the rates $k_{95}$ and $k_{97}$, both sets of rates are in good agreement (S24, b). The fastest process is described by $k_{20} \approx 10^{-1}$ $min^{-1}$ which governs the reaction Ins$P_5$[2OH] $\rightarrow$ Ins(1,4,5,6)$P_4$. This result is in good agreement with MINPP1's annotation as a phosphatase that predominantly removes the phosphoryl group at the 3-position.[6] The reaction rates of the subsequent dephosphorylation steps are separated by at least one order of magnitude, where we get $k_{ij} \approx 10^{-2}$ $min^{-1}$ for reactions of the type Ins$P_4$ $\rightarrow$ Ins$P_3$, $k_{ij} \approx 10^{-4}$ $min^{-1}$ for reactions of the type Ins$P_3$ $\rightarrow$ Ins$P_2$ and $k_{ij} \approx 10^{-5}$ $min^{-1}$ for reactions of the type Ins$P_2$ $\rightarrow$ Ins$P_1$.
In S23, we compare the scaled experimental data (dotted lines ) to the progress curves (solid lines) predicted from the numerically determined set of rates (eq. 2.7) for each replica. The predicted progress curves match the experimental data both qualitatively and quantitatively which strongly supports the assumption that the reaction rates in the Ins$P_5$[2OH] dephosphorylation network are time independent. Furthermore, the results

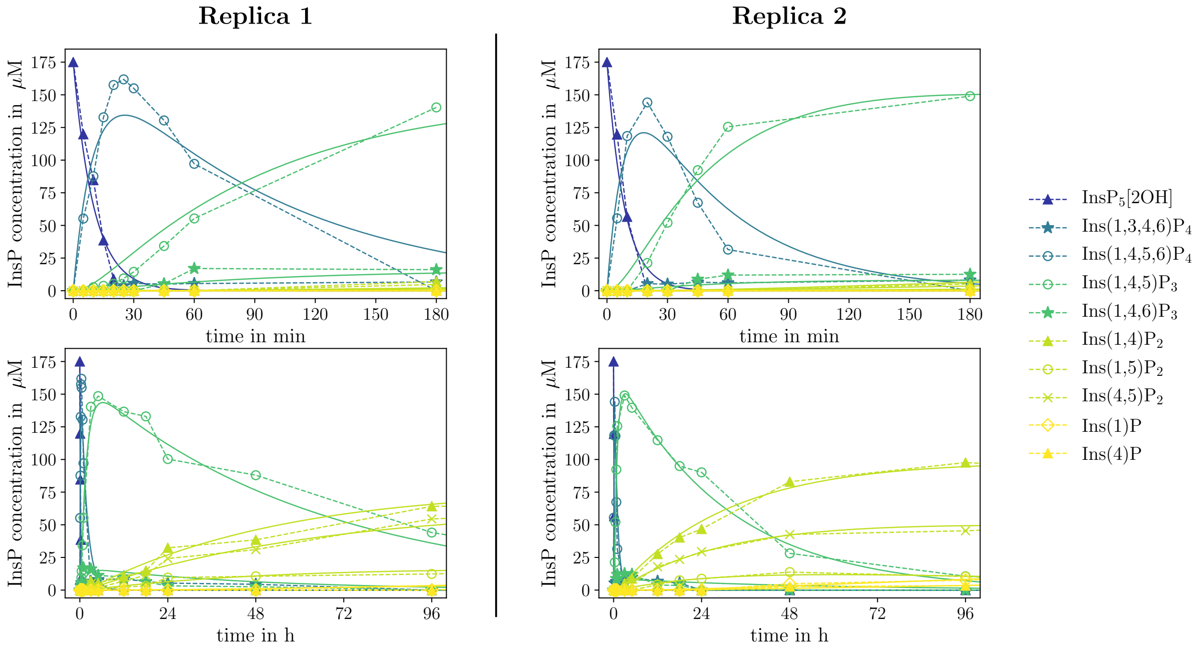

S 23:
Predicted progress curves (solid lines) obtained via minimization routine and scaled experimental data (dashed lines) for two replicas (columns) of MINPP1 reaction with 175 $\mu$M $[^{13}C_6]InsP_5[2OH]$, where the top row magnifies the first 180 min and the bottom row the entire time axis of the experiment. The results for replica 1 are a copy of the results shown in main part Fig. 6a.

**(a)**

| rate | Replica 1 | Replica 2 |
|---|---|---|
| $k_{10}$ | $0.00^*$ | $0.00^*$ |
| $k_{20}$ | $9.8 \cdot 10^{-2}$ | $1.1 \cdot 10^{-1}$ |
| $k_{41}$ | $0.00^*$ | $0.00^*$ |
| $k_{32}$ | $9.3 \cdot 10^{-3}$ | $1.9 \cdot 10^{-2}$ |
| $k_{42}$ | $1.0 \cdot 10^{-3*}$ | $1.0 \cdot 10^{-3*}$ |
| $k_{53}$ | $1.1 \cdot 10^{-4}$ | $3.1 \cdot 10^{-4}$ |
| $k_{63}$ | $3.7 \cdot 10^{-5}$ | $5.9 \cdot 10^{-5}$ |
| $k_{73}$ | $1.0 \cdot 10^{-4}*$ | $1.8 \cdot 10^{-4*}$ |
| $k_{54}$ | $3.5 \cdot 10^{-4}$ | $3.0 \cdot 10^{-4}$ |
| $k_{85}$ | $4.0 \cdot 10^{-6}$ | $4.3 \cdot 10^{-6}$ |
| $k_{95}$ | $1.2 \cdot 10^{-5}$ | $1.0 \cdot 10^{-6}$ |
| $k_{86}$ | $4.5 \cdot 10^{-5}$ | $1.1 \cdot 10^{-4}$ |
| $k_{97}$ | $1.0 \cdot 10^{-6}$ | $1.5 \cdot 10^{-5}$ |

**(b)**

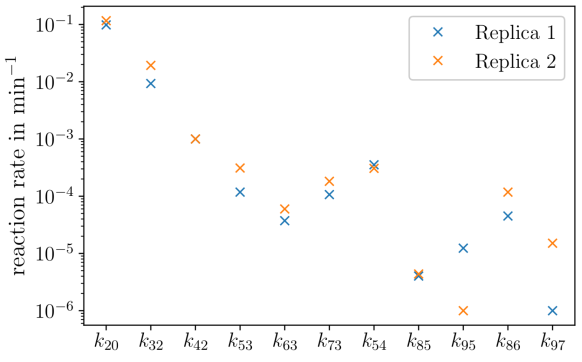

S 24:
(a) Computed rates in $\text{min}^{-1}$ for both replicas of the $[^{13}C_6]InsP_5[2OH]$ dephosphorylation. The column for replica 1 is a copy of the results presented in main part Fig. 6b. *) The marked rates are subject to constraints.
(b) Visual comparison of the rates computed from the scaled experimental data of replica 1 and replica 2, respectively. The rates $k_{10}, k_{41} = 0$ are not included in the representation.

confirm that the network depicted in S21 accurately describes the MINPP1-mediated dephosphorylation of $InsP_5[2OH]$.

#### 4.2 Results $InsP_6$ dephosphorylation

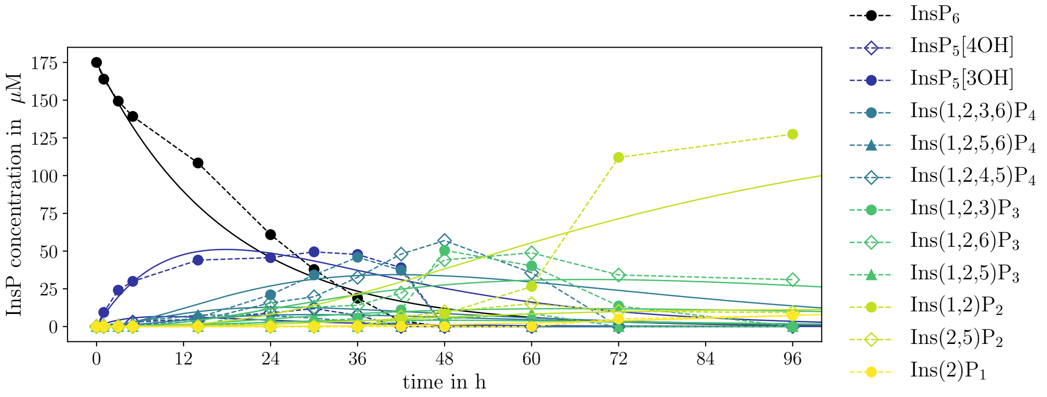

S 25:
Predicted progress curves (solid lines) obtained via minimization routine and scaled experimental data (dashed lines) of MINPP1 reaction with 175 $\mu$M [$^{13}C_6$]$InsP_6$.

SI Table 5:
Computed reaction rates in $min^{-1}$ for [$^{13}C_6$]$InsP_6$ dephosphorylation network.
*) The marked rates were subject to constraints.

| rate | reaction rate |
|---|---|
| $k_{10}$ | $1.90 \cdot 10^{-4*}$ |
| $k_{20}$ | $7.42 \cdot 10^{-4*}$ |
| $k_{31}$ | $2.95 \cdot 10^{-3}$ |
| $k_{41}$ | $1.00 \cdot 10^{-6}$ |
| $k_{42}$ | $2.88 \cdot 10^{-4}$ |
| $k_{52}$ | $6.49 \cdot 10^{-4*}$ |
| $k_{63}$ | $5.10 \cdot 10^{-4}$ |
| $k_{73}$ | $3.67 \cdot 10^{-4}$ |
| $k_{74}$ | $1.56 \cdot 10^{-3}$ |
| $k_{84}$ | $2.13 \cdot 10^{-5}$ |
| $k_{85}$ | $5.99 \cdot 10^{-4}$ |
| $k_{96}$ | $2.02 \cdot 10^{-4}$ |
| $k_{97}$ | $2.22 \cdot 10^{-4}$ |
| $k_{98}$ | $3.88 \cdot 10^{-3}$ |
| $k_{108}$ | $1.00 \cdot 10^{-3}$ |
| $k_{119}$ | $1.00 \cdot 10^{-5}$ |
| $k_{1110}$ | $1.49 \cdot 10^{-4}$ |

S25 shows the comparison between the scaled experimental data (dashed lines) and the progress curves predicted by the numerically determined rates (solid lines). We can clearly see that the computed rates yield a poor representation of the experimental progress curves which strongly indicates that the applied time-constant rates model is insufficient to describe the $InsP_6$ dephosphorylation. The shapes of the experimental progress curves already indicate a kinetic network with time-dependent rates, e.g. the $InsP_6$ progress curve does not represent an exponential decay as we would expect from a first-order reaction. Instead, we observe a damped decrease which could emerge from a inhibition process. As mentioned in main part (Fig. 6c) we suggest that $InsP_6$ itself could act as an inhibitor for the dephosphorylation of its own MINPP1-mediated intermediates.[7] This assumption is further supported by the fact, that the kinetics clearly accelerate as soon as $InsP_6$ is fully depleted. However, we can roughly approximate the rate at which $InsP_6$ is depleted as $k_{10} + k_{20} = 9.3 \cdot 10^{-4}\ \mathrm{min}^{-1}$. Based on our results and the discussion above, we conclude that our master equation ansatz (SI section 2.5) is not capable to capture the true kinetics for the MINPP1-mediated dephosphorylation of $InsP_6$ and thus does not provide any more insight into the main pathways that generate the enantiomers Ins(1,2)$P_2$ and Ins(2,3)$P_2$. For the sake of completeness, we report the numerically determined rates in SI table 5 but did not include these results in the main part of our work. We renounce to extend our ansatz in order to include inhibition processes but this kind of analysis is beyond the scope of this paper.
